# Multidirectional Cellular Plasticity in Ewing Sarcoma Reveals a Tumor–CAF Continuum and Novel Immunotherapeutic Targets

**DOI:** 10.64898/2026.08.03.742600

**Authors:** Mohamed A. Sharif, Abdul S. Khan, Renier J. Brentjens, Scott H. Olejniczak, Henry G. Withers, Joyce E. Ohm

## Abstract

Ewing sarcoma (EwS) is an aggressive pediatric malignancy with poor outcomes for patients with metastatic or relapsed disease. Effective immunotherapeutic approaches, including CAR T-cell therapy, are limited by intratumoral heterogeneity, an incompletely characterized tumor microenvironment (TME), and a lack of well-defined, tumor-restricted target antigens. To address these limitations, we performed an integrated analysis of EwS tumor samples using both single-nucleus and single-cell RNA sequencing datasets derived exclusively from patient samples, including matched primary tumors and orthotopic patient-derived xenograft (PDX) models. Our analyses reveal that primary EwS tumors are largely composed of highly heterogeneous malignant cell populations occupying multiple, multidirectional transcriptional states, including neuronal-like, proliferative, angiogenic, and fibroblast-like. We demonstrate that the EwS TME contains both classical cancer-associated fibroblasts (CAFs) and abundant EwS CAF-like tumor cells that transcriptionally resemble stromal cells while retaining tumor identity. Trajectory analyses define a progressive and coordinated tumor–CAF continuum, marked by gradual loss of neuronal programs and activation of mesenchymal and extracellular matrix remodeling programs, suggesting dynamic tumor cell reprogramming that may promote invasion, immune evasion, and therapeutic resistance. Notably, this structured transcriptional continuum was prominent in primary tumors but largely absent in matched PDX models, underscoring the importance of native tumor context for capturing clinically relevant tumor–TME interactions. We also developed a systematic surface-antigen discovery pipeline and identified ten novel putative tumor-associated surface target antigens (TAs), LRRC15, ATP2B3, CACNA1I, DCHS2, DSEL, LPAR4, PRRT4, TMEM229A, UNC5A, and UNC79, none of which have been previously described in EwS tumor biology. Characterization of these surface TAs revealed distinct expression patterns across EwS tumor cells, EwS CAF-like tumor cells, and classical CAFs. Moreover, some TAs expression differed between primary and metastatic tumors and between primary patient samples and matched PDX models, highlighting the critical importance of first validating therapeutic targets in primary tissues (PT). Together, these findings redefine the cellular architecture of EwS by revealing a dynamic tumor–CAF continuum that is uniquely preserved in primary tumors and establishes a framework for identifying clinically relevant tumor-associated surface TAs. These results provide a foundation for the rational development of next-generation immunotherapies and precision-targeted therapies for patients with EwS.

## Introduction

Ewing sarcoma (EwS) is an aggressive malignancy of bone and soft tissue with a high propensity for early metastatic spread (1, 2). Although rare, EwS is the second most common primary bone malignancy, accounting for approximately 10–15% of all bone sarcomas and more than 20% of pediatric solid tumors (3). Despite being described over a century ago, the current standard of care for EwS still relies largely on conventional multimodal regimens established in the 1970s, consisting of multiagent chemotherapy combined with surgery and/or radiotherapy (1, 4, 5). While advances in local control and dose-intensive chemotherapy have improved outcomes for some patients, prognosis remains strongly dictated by the presence of micrometastatic disease at diagnosis. Currently, 5-year survival exceeds 70% in patients with localized disease who respond to multimodal therapy, whereas survival remains below 30% for patients presenting with metastatic disease (20 to 25% at diagnosis) (2, 4, 6). Importantly, current therapeutic strategies are rarely curative, and patients with metastatic or relapsed disease continue to experience poor treatment responses and clinical outcomes. In EwS, approximately one in four patients experience relapse and have no effective alternative options. Additionally, those who relapse within two years face an estimated 5-year overall survival rate of approximately 7% (2, 5, 7). These limitations underscore the need for effective systemic therapies capable of improving survival beyond current clinical options.

Diagnosis of EwS hinges on the identification of a characteristic gene fusion, most commonly between the *EWSR1* gene and an ETS family member, typically *FLI1*. The resulting EWSR1::FLI1 fusion, generated through the t(11;22)(q24;q12) chromosomal translocation (8, 9), occurs in approximately 85% of cases and serves as the principal diagnostic marker of the disease (10, 11). The fusion protein contains the potent N-terminal transcriptional activation domain of EWSR1 and the ETS-family DNA-binding domain of FLI1 (12–15), creating a powerful aberrant transcription factor capable of extensively rewiring the transcriptional landscape of EwS cells through direct or indirect activation or suppression of known target genes (11). Although the precise cell of origin in EwS is not fully resolved and continues to be debated, accumulating evidence supports a mesenchymal and/or neural crest-related origin (16–18). Importantly, the cellular context in which EWSR1::FLI1 is expressed plays a critical role in determining phenotypic outcomes (13, 19–21), and in cell lines sustained fusion expression is required for oncogenic transformation (22, 23). While substantial advances have been made in understanding the molecular biology of EWSR1::FLI1, the complexity of EwS biology and the lack of robust *in vivo* systems have hindered comprehensive studies of EwS oncogenesis and therapeutic target development (24–26).

Therapeutic stagnation in EwS is primarily driven by the remarkable complexity and heterogeneity of the tumor cells and tumor microenvironment (TME) (10, 16, 27, 28), which has hindered the development of targeted and durable therapies. Antibody-based approaches, including those targeting IGF-1R, have produced meaningful and occasionally durable responses in only a subset of patients (29, 30), whereas immune checkpoint inhibitors have demonstrated limited clinical activity in EwS (31, 32). Recent studies in EwS cell lines and xenograft models have revealed complex transcriptional heterogeneity within these cells. The EWSR1::FLI1 expression level varies between individual cells, influencing the transcriptional landscape and creating diverse subpopulations with distinct phenotypic cell states that drive carcinogenesis and facilitate adaptation. High levels of fusion expression are associated with a proliferative and oncogenic cellular state, whereas low levels correspond to migratory and invasive subpopulations (33). The composition of these cell-to-cell differences in EWSR1::FLI1 expression varies across EwS cell lines (33). On this transcriptional landscape, a hybrid subpopulation, termed EwS CAF-like cells, exhibits high expression of fusion-activated genes while fusion-suppressed genes are derepressed. These cells display characteristics similar to cancer-associated fibroblasts (CAFs), including the secretion of extracellular matrix (ECM) proteins and the ability to remodel the TME (28, 34). Additionally, in EwS CAF-like cells, reciprocal crosstalk between EWSR1::FLI1 and Wnt/TGF-β signaling establishes a positive feedback loop that reinforces the CAF-like state, demonstrating how both tumor-intrinsic and TME-derived signals dynamically shape fusion activity and EwS cellular states (34, 35). This dynamic interplay between tumor-intrinsic programs and the TME cues suggests that EwS cellular states may differ across biological contexts. Consequently, it remains unclear how cellular heterogeneity varies between primary and metastatic tumors or how these transcriptional programs are represented in xenograft models, which lack a functional immune system.

The presence of EwS CAF-like subpopulations is particularly relevant because stromal remodeling and subsequent immune suppression are increasingly recognized as important contributors to tumor progression (28). CAFs have been extensively studied in epithelial malignancies, where they promote extracellular matrix deposition, immune suppression, metastasis, and therapeutic resistance (36). In sarcomas, however, characterization of CAFs is more challenging because tumor and stromal cells share mesenchymal features and overlapping transcriptional programs (37, 38). Consequently, it remains unclear whether non-tumor (classical) CAFs are present within the EwS TME, and if so, how they transcriptionally differ from EwS CAF-like cells. Emerging evidence suggests that EwS CAF-like cells possess enhanced invasive and metastatic potential and may actively participate in remodeling the local TME (28). Therefore, defining the presence and distribution of classical CAFs, distinguishing them from EwS CAF-like cells, and characterizing their respective transcriptional programs are critical for understanding the cellular organization of the EwS TME and factors that contribute to disease progression.

This heterogeneity also has important implications for the identification of tumor-associated surface antigens (TAs) and the development of targeted immunotherapies. Patients with relapsed or refractory EwS continue to have a poor prognosis, and despite considerable interest in immunotherapeutic strategies, clinical responses have remained limited (39, 40). One major challenge is the immunosuppressive TME, which can contain stromal barriers, CAFs, tumor-associated macrophages (TAMs), myeloid-derived suppressor cells (MDSCs), and other cell populations capable of limiting immune-cell infiltration and suppressing antitumor immune responses (39, 41). Although antigen-directed therapies, including monoclonal antibodies and chimeric antigen receptor (CAR) T-cell therapies, have transformed the treatment of several hematologic malignancies, their clinical efficacy in EwS has remained limited (39). Successful immunotherapeutic targeting requires antigens that are highly expressed on tumor cells, minimally expressed on normal tissues, and sufficiently represented across heterogeneous tumor populations to minimize antigen escape (42). The transcriptional diversity observed in EwS suggests that distinct cellular states, including the aggressive CAF-like populations involved in TME remodeling and tumor progression (28), may express unique antigens that could be therapeutically exploited.

Despite the growing recognition of EwS transcriptional heterogeneity, important questions remain regarding how these cellular states are represented in patient tumors, how they are maintained in matched patient-derived xenograft (PDX) models, and how they influence the landscape of therapeutically targetable surface TAs. In addition, the presence of classical CAFs and EwS CAF-like cells within primary tissue (PT) is not fully explored. To address these gaps, we performed extensive single cell transcriptome analysis of PT and matched PDX samples to characterize EwS cellular heterogeneity, define the composition of the TME, and identify novel tumor-associated surface antigens. Here, we further characterize the complexities of EwS TME in primary patient samples. Our findings indicate that there is a high level of heterogeneity among the EwS cells in multiple transcriptional directions. While EwS tumors exhibit a unique neuronal-like transcriptional signature, the transcriptional profiles of EwS clusters are distinct, with neuronal genes expressed both globally across tumors and discretely within specific clusters. In addition, CAF markers were selectively expressed in specific EwS clusters, further underscoring the transcriptional heterogeneity within these tumor cells. EwS TME does have few classical CAFs and proportionally is much fewer in comparison to EwS CAF-like cells. Classical CAFs have a different transcriptional signature than EwS CAF-like cells. Within the tumor, EwS cell subpopulations occupy distinct transcriptional states along a trajectory toward a CAF-like phenotype. We also observed transcriptional differences between PT and PDX models, which may be attributed to several factors, potentially including, as previously discussed, selective growth pressures applied to the heterogeneous landscape established by EWSR1::FLI1 fusion-driven transcriptional activity. Together, these results show extensive heterogeneity among EwS tumor cells, which may contribute to the clinical challenges associated with eradicating this disease, including a stepwise progression of EwS cells toward a more mesenchymal CAF-like state. Others have shown that these EwS CAF-like cells have increased invasive and metastatic potential (28), and we now highlight the potential for specific targeting of this aggressive subpopulation. Using a systematic surface TAs discovery framework, we identify 10 novel putative TAs for immunotherapeutic targeting in EwS. All 10 genes are reported here for the first time in EwS tumor biology, highlighting a set of largely unexplored targets with potential to expand the current landscape of immunotherapy and enable the development of more precise, tumor-restricted treatment strategies. Collectively, these findings provide a comprehensive view of EwS cellular heterogeneity in primary tumor samples and establish a framework for developing immunotherapeutic strategies capable of targeting biologically distinct and clinically relevant tumor cell populations.

## Methods

### Dataset Acquisition

All datasets were acquired from publicly available databases. The Ewing Sarcoma (EwS) single-nucleus RNA-sequencing (snRNA-seq) data were obtained from the single-cell Pediatric Cancer Atlas (ScPCA) Portal (Alex’s Lemonade Stand Foundation Childhood Cancer Data Lab) (43). We analyzed the EwS dataset (Project SCPCP000015), which contains 7 patient tumor samples and 6 matched orthotopic patient-derived xenograft (O-PDX) samples generated by investigators at St. Jude Children’s Research Hospital. Raw count matrices and accompanying metadata were downloaded directly from the ScPCA Portal. For the single cell RNA-sequencing (scRNA-seq) dataset, samples were obtained from the Gene Expression Omnibus (GEO) under accession number GSE243347. This dataset contains 18 primary EwS patient samples and was downloaded as raw count matrices with associated metadata. Within the single-cell dataset only 9 samples from 6 patients were used because they were not flow sorted before sequencing (44). Bulk RNA sequencing (RNA-seq) data were obtained from the St. Jude Cloud Pediatric Cancer Genome Project (PCGP) and included 283 tumor samples consisting of EwS, clear cell sarcoma (CCS), osteosarcoma (OS), round cell sarcoma (RCS), and synovial sarcoma (SS) samples (45). RNA expression data for cell lines were obtained from the DepMap portal (Cancer Dependency Map; CCLE dataset (Cancer Cell Line Encyclopedia); https://depmap.org) (46, 47). Protein expression for cell lines was also obtained from DepMap portal using both harmonized CCLE mass spectrometry dataset (harmonized_MS_CCLE_Gygi.csv) and harmonized CCLE reverse-phase protein array (RPPA) dataset (harmonized_RPPA_CCLE.csv) from the DepMap Public 26Q1 release (46, 47). All downstream analyses were performed in R (v4.2.0) using established bioinformatics packages and custom scripts.

### snRNA- and scRNA-seq data processing and quality control

Individual samples were imported and converted to Seurat (v5.4.0) objects while preserving cell- and gene-level metadata. Gene symbols were standardized prior to downstream analyses by replacing unsupported characters in feature (gene) names and establishing a 1-to-1 mapping between gene symbols and Ensembl IDs. Individual samples were subsequently merged into a single Seurat object for each dataset using unique cell prefixes to preserve sample identity. An initial quality control step was performed on the merged datasets by calculating the percentage of mitochondrial transcripts for each nucleus or cell using genes with the “MT-” prefix. Cells or nuclei were filtered based on the number of detected genes, total UMI counts, and mitochondrial transcript percentage to remove low-quality profiles while minimizing the retention of dying cells and doublets. Because scRNA-seq and snRNA-seq exhibit distinct transcriptional characteristics, quality control thresholds were optimized independently for each sequencing modality; however, identical filtering criteria were applied to all samples within each snRNA-seq or scRNA-seq modality to ensure consistent preprocessing across datasets. The actual filtering thresholds are found within the supplemental data section (Sup.Fig. 1 & 3). Following the initial filtering, additional cell-level quality control was performed using SingleCellTK (v2.8.0). Potential doublets were identified using scDblFinder (v1.12.0), which assigns each nucleus/cell a doublet score and classifies each profile as either a singlet or doublet. Ambient RNA contamination was estimated using DecontX (implemented in celda v1.14.2), which models contamination arising from extracellular RNA and estimates a cell-specific contamination fraction. Nuclei or cells passing the initial quality control filters, classified as singlets by scDblFinder, and exhibiting acceptable ambient RNA contamination as determined by DecontX were retained for downstream analyses.

**Figure 1.**
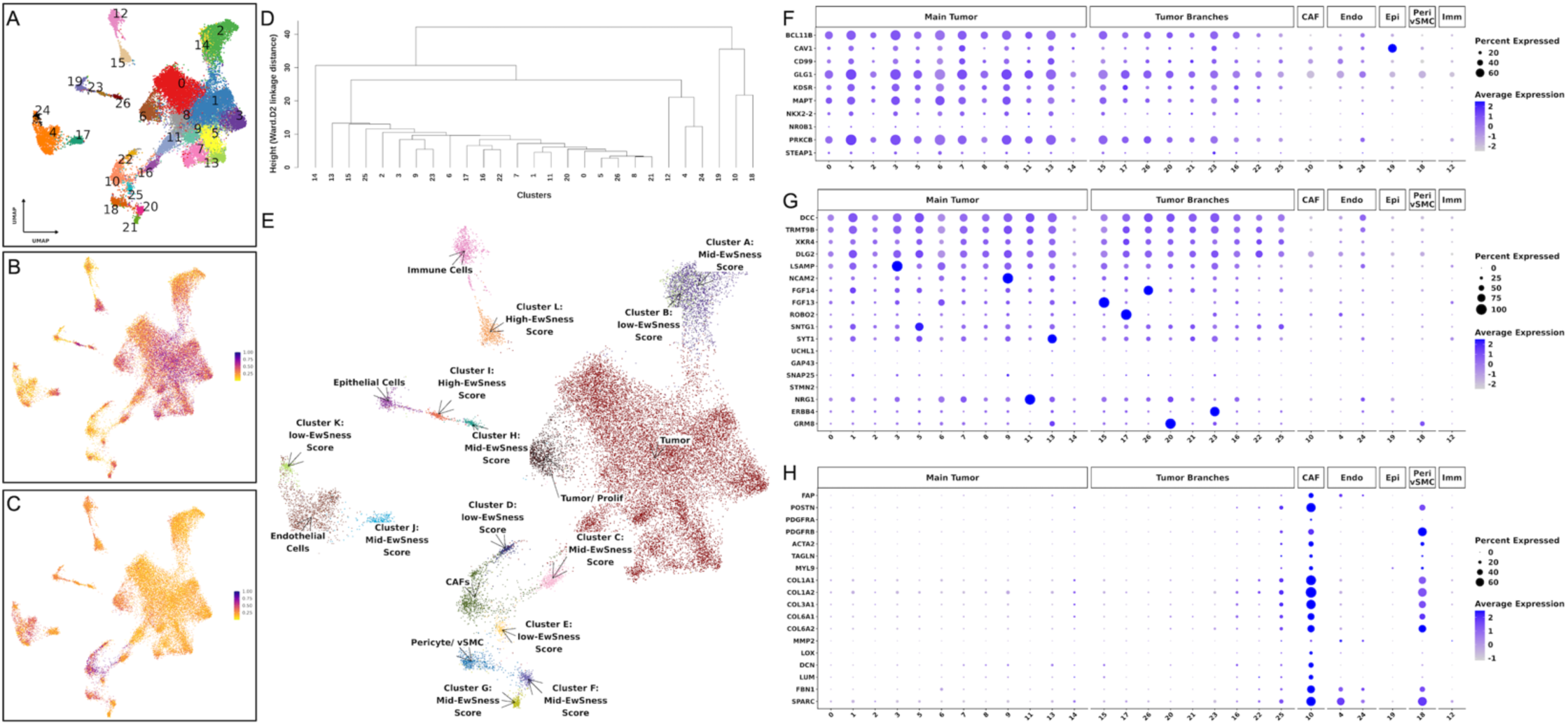
snRNA-seq transcriptomic profiling of EwS PT samples indicates a complex distribution of clusters and cell states. (A) UMAP of snRNA-Seq PT dataset showing unsupervised clustering of nuclei. 27 clusters were identified, and each point represents a single nucleus, colored by its cluster identity. (B) UMAP of normalized AUCell gene set (gene set 1; 32 genes upregulated by the fusion) enrichment scores. (C) UMAP of normalized AUCell gene set (gene set 3; 195 genes downregulated by the fusion) enrichment scores. High-scoring nuclei (purple) correspond to higher gene set expression, while low-scoring nuclei (yellow) correspond to lower gene set expression. The full list of the 11 gene sets and their information is in Sup.Table 3. (D) Hierarchical clustering of defined clusters based on PCA centroid distances. Dendrogram depicting relationships among 27 derived clusters (0–26). Cluster centroids were calculated from the first 20 principal components, and Euclidean distances were used for hierarchical clustering with Ward’s method (Ward.D2). Branch height reflects inter-cluster dissimilarity, with shorter branches indicating greater transcriptional similarity. (E) Annotated UMAP of ALSF snRNA-seq clusters highlighting major cell types, main tumor, and clusters that form the tumor branches. This annotation of the UMAP factors in various cell typing methods including using a reference database, CNV analysis, gene set analysis, hierarchical clustering, differential gene analysis (Final_All_Markers & Find_Marker), and top100 gene expression per cluster. Clusters that are considered as tumor branches are labeled with EwSness Scores. (F - H) Dot plot showing expression of selected (F) EwS, (G) neuronal, (H) CAF markers across clusters in different categories (Main Tumor, Tumor Branches, CAF, Endo, Epi, Peri/vSMC, Imm) of the UMAP. In each dot plot, the y-axis shows the list of genes and the x-axis shows the cluster numbers from the UMAP projections. Dot size represents the percentage of nuclei expressing each gene, and color intensity indicates the average expression levels.

### snRNA- and scRNA-seq data normalization, integration, dimensionality reduction, and clustering

Following quality control, expression data were normalized independently using two complementary approaches implemented in Seurat (v5.4.0). First, gene expression values were normalized using the LogNormalize method, and highly variable genes were identified for exploratory analyses. To improve normalization across multiple patient samples, we also performed SCTransform normalization, which applies regularized negative binomial regression while regressing out mitochondrial transcript content. Principal component (PC) analysis was subsequently performed using the identified variable features to reduce data dimensionality, and the optimal number of principal components retained for downstream analyses was determined by examination of elbow plots. To correct for batch effect driven by interpatient heterogeneity, while preserving biological variation, each patient sample was treated as an individual layer, and multiple integration strategies were evaluated using Seurat’s IntegrateLayers framework. Canonical correlation analysis (CCA), reciprocal principal component analysis (RPCA), and Harmony integration were performed using the SCTransform-normalized data, and the resulting integrated embeddings were compared by visual inspection of Uniform Manifold Approximation and Projection (UMAP) and t-distributed stochastic neighbor embedding (t-SNE) projections. Based on the degree of sample mixing and preservation of biologically meaningful cell populations, Harmony integration was selected for all subsequent analyses. Following Harmony integration, a shared nearest-neighbor graph was constructed using the integrated PC (dims = 20), and unsupervised graph-based clustering was performed to identify transcriptionally distinct cell populations. For all these analyses we used seed.use = 2012 and dims = 1:20 (PCs). UMAP was used as the primary method for visualization of the integrated datasets, while t-SNE embeddings were generated for comparison during optimization of the integration workflow. Cluster stability and sample representation were evaluated by examining the distribution of cells according to patient identity, sample origin, and clinical annotations across the low-dimensional embeddings. The final integrated Harmony reduction was used for all downstream analyses, including cluster annotation, differential gene expression, trajectory inference, and surface target antigens (TAs) Identification.

### CNV inference and malignant cell identification

To distinguish tumors from non-tumor nucleus/cell clusters, CNV analysis was performed using SCEVAN (v1.0.3). Following Harmony integration, RNA count layers were rejoined, and the raw gene expression count matrix was extracted from the RNA assay for downstream CNV analysis. Gene symbols were converted to contain only Ensembl IDs to ensure compatibility with SCEVAN while maintaining a 1-to-1 mapping with features and counts. SCEVAN was applied directly to the raw count matrix using the human reference genome to infer large-scale chromosomal copy number alterations and reconstruct tumor subclonal architecture. The pipeline classifies each cell as tumor, normal, or filtered based on its inferred CNV profile and identifies genetically distinct tumor subclones through unsupervised analysis of genome-wide copy number alterations. The resulting SCEVAN classifications were integrated with the Seurat metadata using dplyr (v1.2.1) and tidyr (v1.3.2) to assign malignant status to individual cells and evaluate the distribution of tumor and normal cells across transcriptionally defined clusters, patient IDs, and sample IDs. The inferred CNV profiles were subsequently used to distinguish tumor clusters from non-tumor populations and to support downstream cluster annotation and biological interpretation.

### SingleR-based cell type annotation

Cell type annotation of the integrated EwS snRNA-seq and scRNA-seq datasets were performed using SingleR (v2.0.0) with the Human Primary Cell Atlas reference dataset from celldex (v1.8.0). The final Harmony-integrated Seurat object was loaded, and cluster identities were set using the Harmony-derived cluster assignments. Prior to annotation, the RNA assay layer was set to the default assay. SingleR was applied to the RNA expression matrix using the “HumanPrimaryCellAtlasData” reference. Per-nucleus/cell annotation scores, predicted labels, pruned labels, & maximum prediction scores were incorporated into the Seurat metadata. SingleR score matrices were then aligned to Seurat cell barcodes, and cluster-level annotations were summarized by averaging SingleR scores across Harmony clusters. Heatmaps of mean SingleR reference scores were generated using pheatmap (v1.0.13), with scores scaled either by row or by column to visualize reference-cell-type enrichment across transcriptionally defined clusters.

### AUCell-based gene-set scoring for cell-state annotation

Cell-level gene-set activity was quantified using AUCell (v1.20.2). The log-normalized expression matrix was extracted from the Seurat object, and feature names were reformatted to retain gene symbols by removing appended Ensembl IDs. Gene-set collections were loaded from GMT files using GSEABase (v1.60.0); Sup.Table 3 provides more information on the gene sets. For each cell, genes were ranked according to expression using ‘AUCell_buildRankings’, and enrichment of each gene set within the top-ranked genes was calculated using ‘AUCell_calcAUC’. AUCell thresholds were examined using ‘AUCell_exploreThresholds’, and cells with AUC scores above the default threshold for each gene set were classified as active for that signature. Raw AUC scores and binary activity call labels were aligned to the Seurat cell barcodes and added to the Seurat metadata. AUC scores were also min–max scaled from 0 to 1 for UMAP visualization. To summarize gene-set activity across transcriptionally defined populations, mean scaled AUC scores and the percentage of AUCell-active cells were calculated for each cluster.

### Marker gene-based cell type annotation

Cell identities were also assigned using a marker gene-based annotation strategy. Canonical marker genes representing various cell types and cellular programs, including EwS tumor cell-specific genes, were curated from the literature and evaluated across each cluster. Additionally, cluster-specific marker genes identified by FindAllMarkers were ranked by average log2 fold-change, and the top 10 differentially expressed genes from each cluster were examined alongside canonical lineage markers, AUCell gene-signature enrichment, SingleR cell-type predictions, top 100 highest-expressed genes per cluster, and inferred CNV profiles generated by SCEVAN.

### Trajectory inference, pseudotime analysis, and functional enrichment

To investigate dynamic transcriptional changes and cellular state transitions, trajectory inference was performed using Monocle 3 (v1.14.26). The Harmony-integrated Seurat object was converted to a Monocle 3 cell_data_set object while preserving cell-level metadata and low-dimensional embeddings. The Harmony-derived UMAP coordinates were used as the input dimensional reduction, and Harmony cluster assignments generated in Seurat were transferred to the Monocle object to maintain consistent cluster identities throughout the analysis. Trajectory graphs were constructed using the learn_graph function with graph learning performed independently within each partition (use_partition = TRUE). Pseudotime was calculated using the ‘order_cells’ function, with a harmony cluster designated as the root population based on its inferred biological position within the trajectory. Cells were subsequently ordered along the learned principal graph to construct a continuous transcriptional progression.

### Candidate TAs identification pipeline

Candidate tumor-associated TAs were identified using a stepwise filtering pipeline designed to prioritize plasma membrane-associated proteins enriched in EwS and CAF cell populations while exhibiting limited expression in all other non-tumor clusters. First, an initial list of 2,467 genes annotated as plasma membrane-associated in the Human Protein Atlas (HPA) Subcellular resource was obtained and filtered to retain genes encoding predicted membrane proteins (48). Second, genes exhibiting limited or undetectable expression across HPA scRNA-sequencing datasets (classified by HPA as having RNA single-cell-type detection in “some,” “single,” or “none”) were retained, yielding 322 candidate genes. Third, these candidates were then matched to the genes detected in the EwS snRNA-seq and scRNA-seq datasets. Fourth, within each dataset, the 322 candidate genes were assessed to determine whether they ranked among the top 30 differentially expressed genes for each transcriptionally defined cluster. Duplicate genes identified across multiple clusters were consolidated to generate a unique list of candidates. Fifth, genes highly ranked in non-tumor nucleus/cell clusters were removed. Sixth, genes exhibiting substantial background expression across normal nuclei/cells were removed. Finally, the remaining genes were visualized across all clusters, and candidates with low expression in EwS tumor populations were excluded. This pipeline identified 40 candidate TAs in the snRNA-seq dataset and 25 in the scRNA-seq dataset, of which 14 were shared between the two datasets.

### Bulk RNA sequencing data processing and expression analysis

Bulk RNA sequencing (RNA-seq) data from the St. Jude Cloud Pediatric Cancer Genome Project (PCGP) were processed from raw feature counts (gene-level count) files. Sample metadata and feature counts output files were imported and individual count files were merged into a single gene-by-sample count matrix after verifying identical gene ordering across all samples. Sample metadata were manually curated to remove duplicated records, retain relevant clinical annotations, and harmonized disease classifications. Samples were grouped into five sarcoma subtypes, including EWS, osteosarcoma (OS), synovial sarcoma (SS), clear cell sarcoma (CCS), and round cell sarcoma (RCS), with PDX (patient-derived xenograft) samples additionally annotated separately from PT (primary tissues). Count matrices and sample metadata were combined into a DESeq2 (v1.50.2) object. Prior to normalization, lowly expressed genes were removed by retaining genes with at least 10 raw counts in a minimum of 5 samples. Expression counts were subsequently normalized using the DESeq2 median-of-ratios normalization method, which corrects for differences in sequencing depth and RNA composition between samples through estimation of sample-specific size factors. Variance stabilizing transformation (VST) was applied to normalized counts for downstream visualization and multivariate analyses. Quality control was performed by examining density distributions of log-transformed raw count data before and after filtering, allowing assessment of count distribution consistency across samples. Normalized expression values were subsequently used to evaluate expression of the selected putative TAs across all sarcoma subtypes and between PT and PDX samples.

## Results

### Single-Cell Profiling Reveals the Cellular Architecture of Primary EwS TME

We analyzed the single-nucleus RNA-seq (snRNA-seq) dataset from the Alex’s Lemonade Stand Foundation for Childhood Cancer (ALSF) Single-cell Pediatric Cancer Atlas (ScPCA) Portal (43). This dataset contains 13 EwS samples comprising 7 PT samples (from 5 patients) with 6 matched orthotopic patient-derived xenografts (O-PDXs) samples (Sup.Table 1). This dataset enabled us to comprehensively interrogate tumor cells and TME at single-nucleus resolution, allowing for a more careful and detailed characterization of PT cell heterogeneity while dissecting transcriptionally distinct EwS cell subpopulations. Also, it allowed us to assess shifts in the transcriptional landscape of tumor cells due to selective growth conditions through comparisons of sequenced PT and matched PDX samples. After standard Seurat preprocessing, we performed quality control (QC) to remove low-quality nuclei (Sup.Fig. 1) and then applied SCTransform for normalization and PCA (Principal Component Analysis) for initial dimensionality reduction. Following QC filtering, the PT dataset comprised of 27,765 nuclei expressing 58,751,215 UMI counts, whereas the PDX dataset comprised of 32,568 nuclei expressing 43,250,215 UMI counts (Sup.Fig. 1E & F). While not shown, the original dataset exhibited a strong batch effect driven by interpatient heterogeneity. To integrate samples, we compared several integration methods, see Methods section for more detail. We selected Harmony integration because it produced the most coherent and biologically relevant embedding across patient samples (Sup.Fig. 2A – D). Next, we performed an unsupervised SNN (Shared Nearest Neighbor) clustering in the integrated space and identified clusters. We visualized the data using both UMAP and t-SNE (not shown) with matched resolutions. UMAP provided improved separation of transcriptionally distinct cell populations and was therefore used for all subsequent analyses. Within the PT samples we generated 27 (0–26) clusters (Fig. 1A) that are present in both primary and metastatic conditions to varying degree (Sup.Fig. 2D). Unlike metastatic samples, clusters in primary tumors appeared more connected in UMAP space, suggesting a more continuous transcriptional landscape. It is common to observe a continuum of differentiation profiles in tumors, as cancer cells may exist in more dynamic, less discrete differentiation states than in normal cells (49, 50). Additionally, the differences observed between the primary and metastatic states may be influenced by multiple factors, including tumor location, intrinsic tumor heterogeneity, TME pressures, and clonal selection during metastatic dissemination, which together can lead to the emergence of more transcriptionally distinct and specialized cell populations in metastatic lesions.

**Figure 2.**
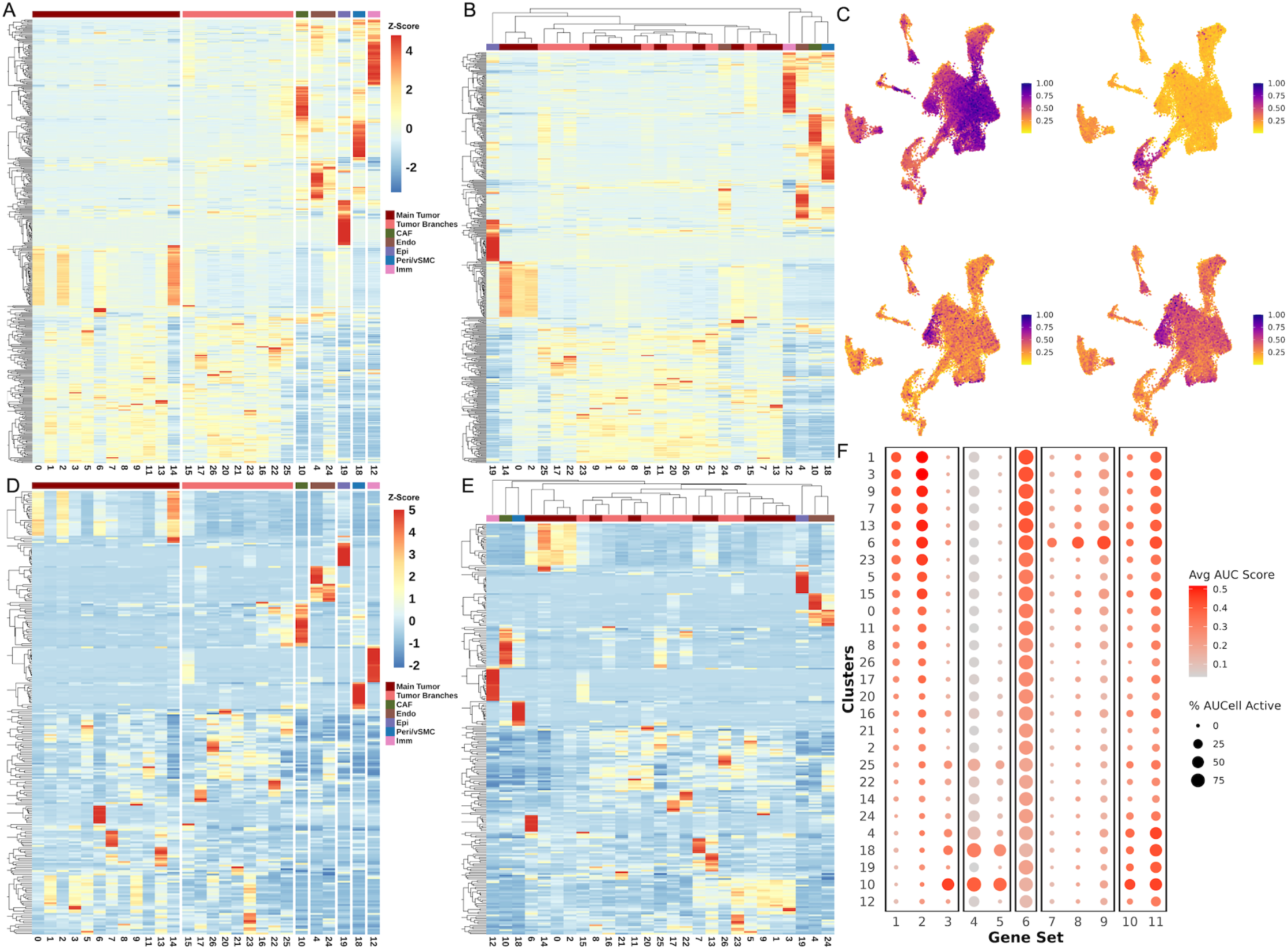
snRNA-seq cluster-specific gene expression and AUCell gene-set activity elucidates transcriptional heterogeneity in the EwS TME. (A & B) Heatmap showing the top 100 highest-expressed genes per defined cluster based on average normalized RNA expression sorted by (A) defined categories or (B) unbiased hierarchical clustering using top 100 genes. (C) UMAP of normalized AUCell gene set enrichment scores. Upper-left (gene set 2; 439 genes upregulated by the fusion), upper-right (gene set 5; 200 hallmark epithelial mesenchymal transition genes), lower-left (gene set 7; 37 gene set activated by ATR response to replication stress), lower-right (gene set 8; 184 gene set involved in DNA replication). High-scoring nuclei (purple) correspond to higher gene set expression, while low-scoring nuclei (yellow) correspond to lower gene set expression. (D & E) Heatmap showing the top 10 differentially expressed markers per cluster, ranked by average log2 fold change sorted by (D) defined categories or (E) unbiased hierarchical clustering using top 10 genes. Expression values are scaled per gene (row-wise Z-score) across clusters. Both genes and clusters are hierarchically clustered and ordered by similarity in their scaled expression profiles, grouping genes and clusters with shared patterns across clusters. Distinct blocks of high expression identify cluster-specific transcriptional signatures and reinforce higher-order relationships among related clusters. (F) Dot-plot summarizing normalized average AUCell scores (red) for each gene set (columns) per cluster (rows). Dot size represents the percentage of nuclei classified as “Active” for each gene set within each cluster (nuclei with gene-set enrichment based on AUCell AUC >= default threshold). Gene sets had minimal overlap, and Sup.Table 3 lists the 11 gene sets and the number of genes in each.

**Table 1.**
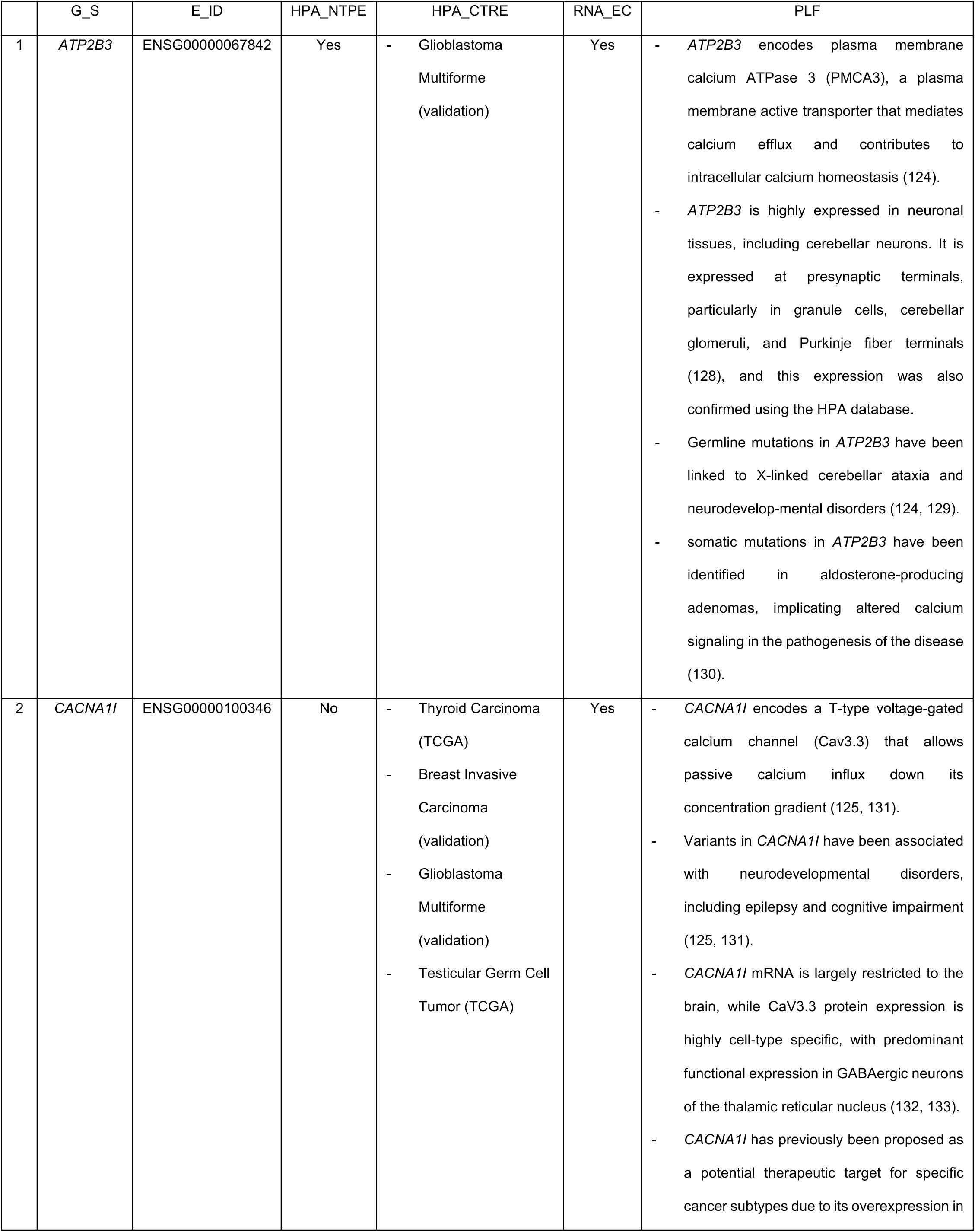
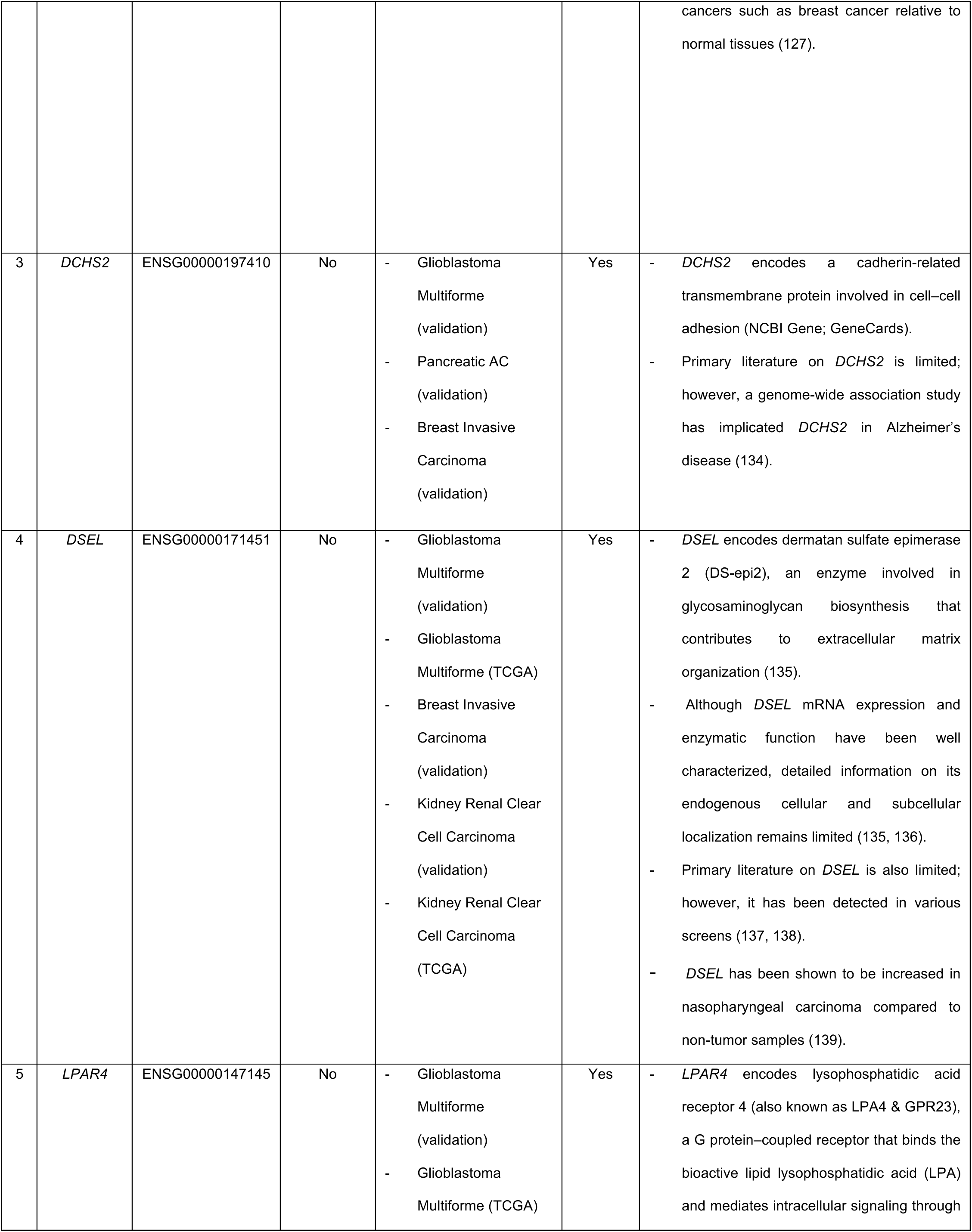
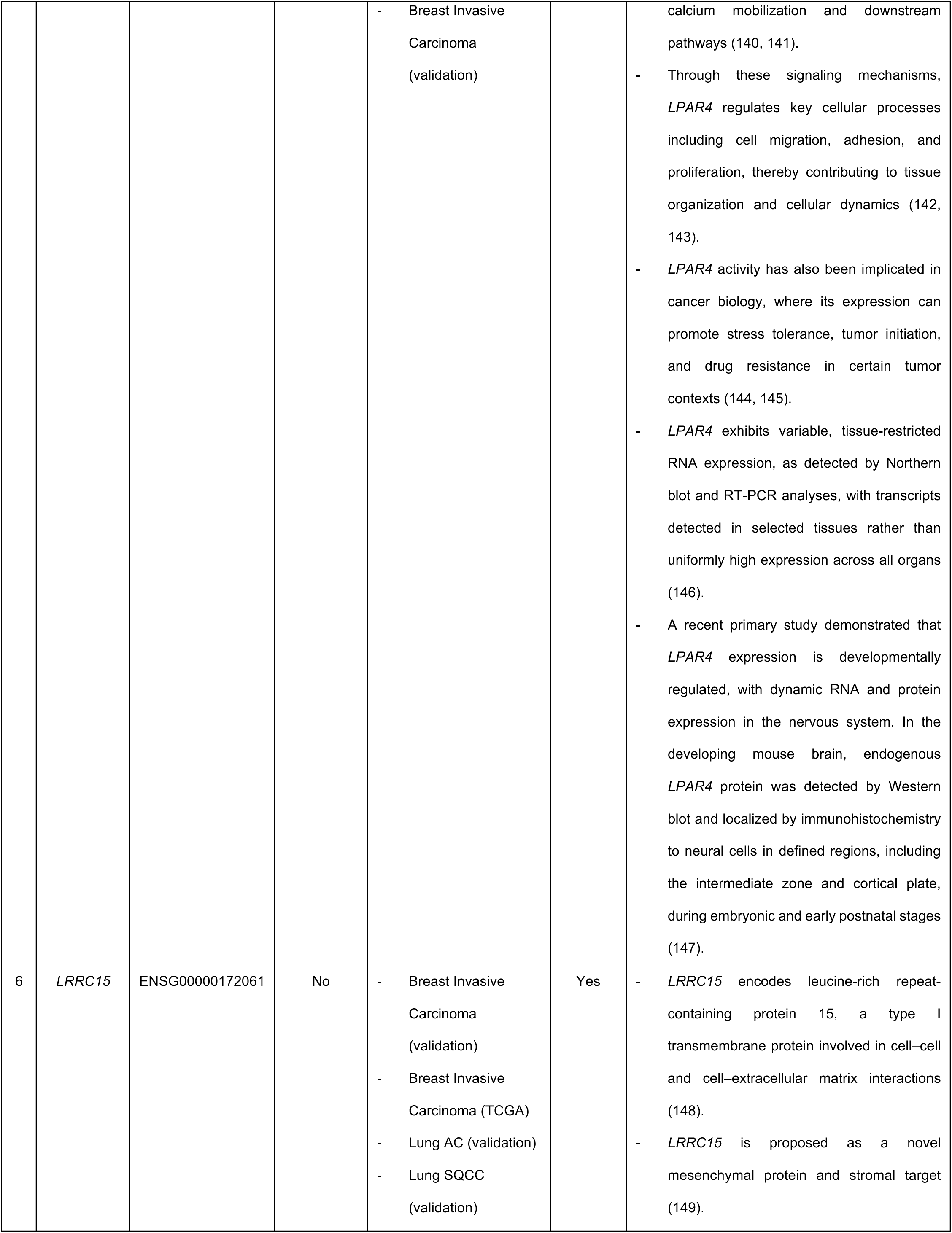
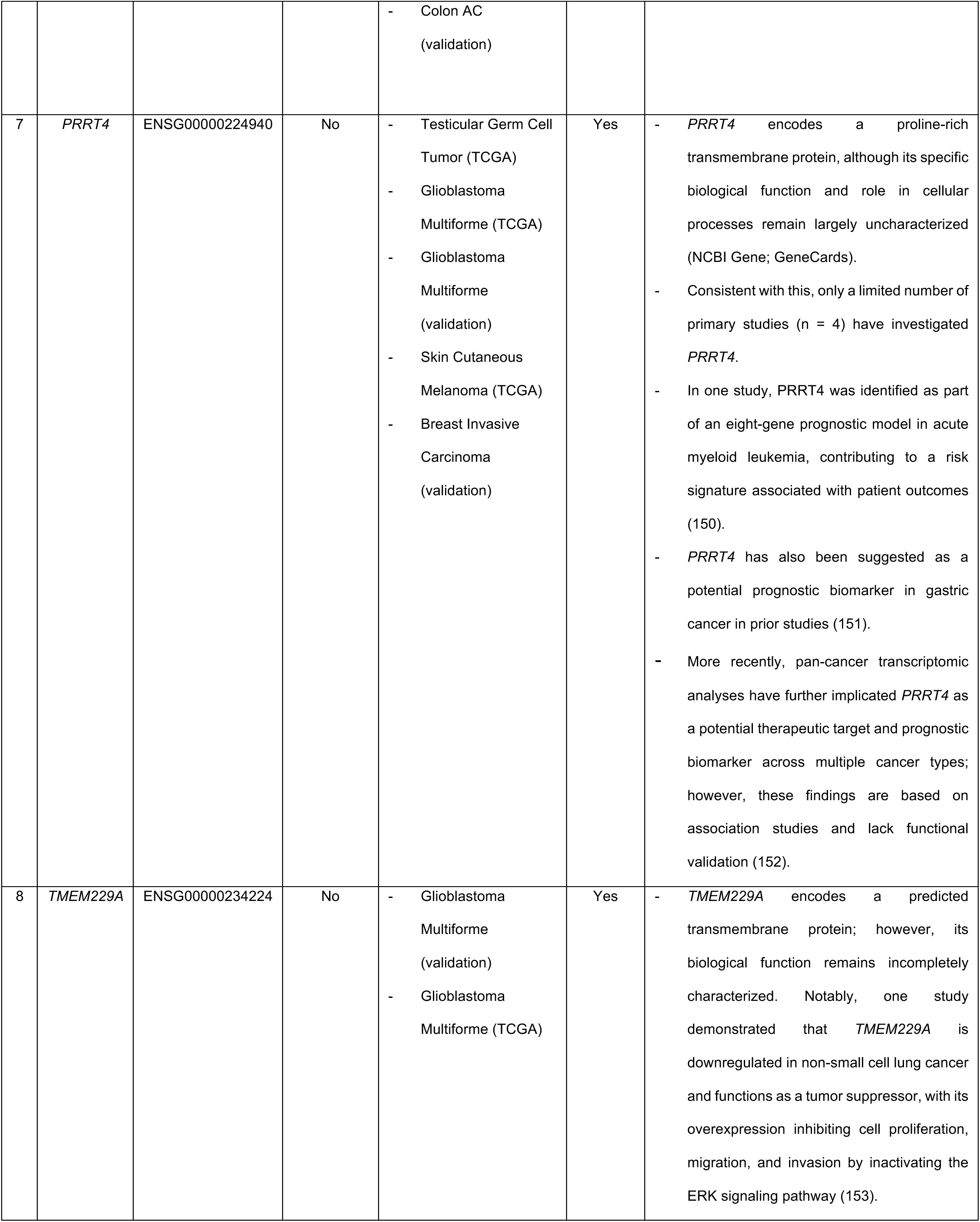
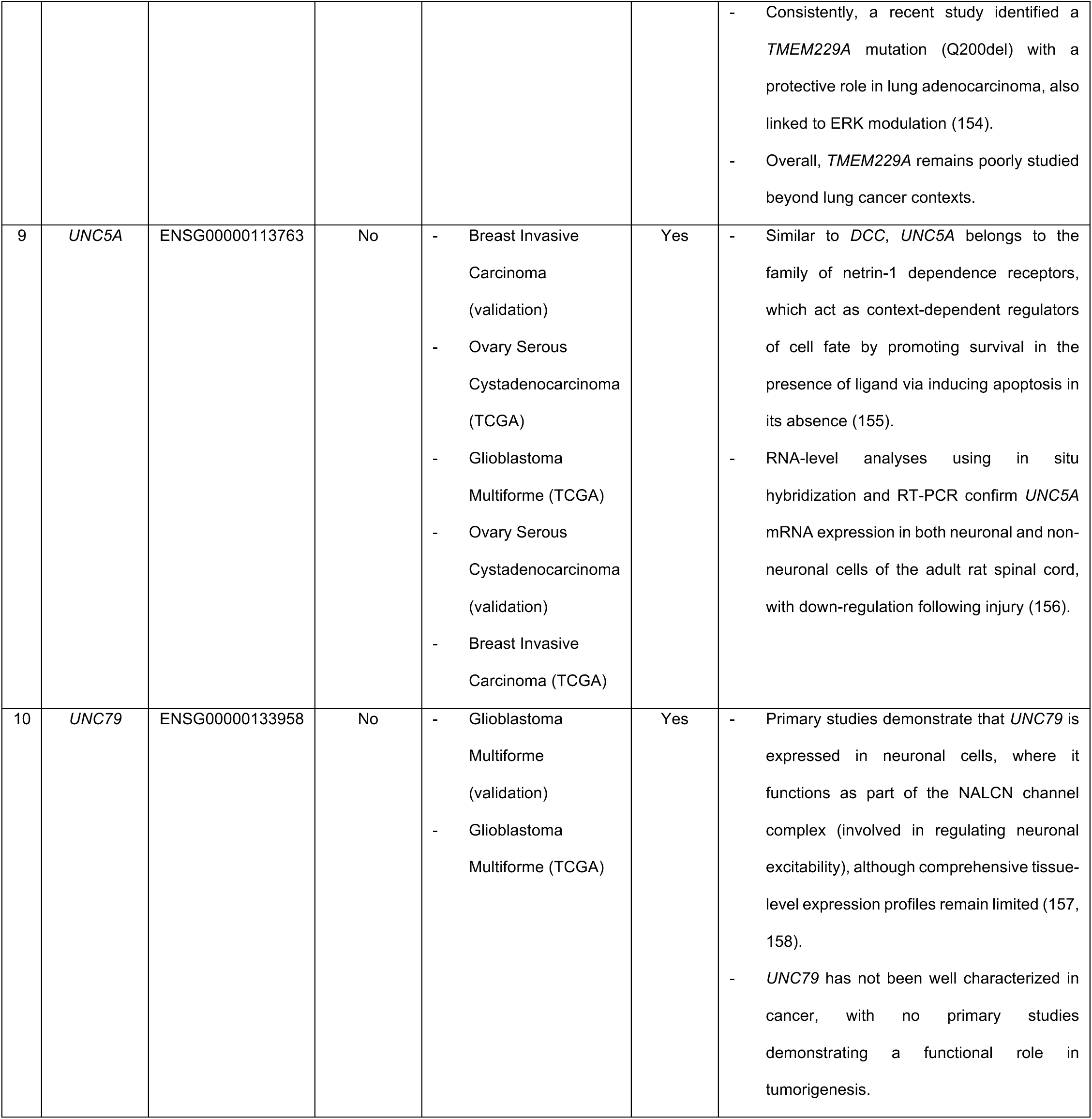
Overview of the top 10 putative EwS surface TAs. This table summarizes candidate target antigens identified in EwS datasets: *ATP2B3*, *CACNA1I*, *DCHS2*, *DSEL*, *LPAR4*, *LRRC15*, *PRRT4*, *TMEM229A*, *UNC5A*, and *UNC79*, along with their corresponding Ensembl gene identifiers (E_ID). Gene symbols are denoted as G_S. Human Protein Atlas normal tissue protein expression (HPA_NTPE) indicates whether a gene is known to be expressed at the protein level across non-tumor tissues within the database. While the Human Protein Atlas cancer tissue RNA expression (HPA_CTRE) indicates RNA expression levels across tumor types, only the top five cancers exhibiting median expression levels ≥0.5 pTPM are listed. RNA expression in EwS cell lines (RNA_EC) was obtained from the DepMap portal (Cancer Cell Line Encyclopedia, CCLE dataset). Primary literature findings (PLF) were curated from PubMed to highlight prior information on each gene.

|  | G_S | E_ID | HPA_NTPE | HPA_CTRE | RNA_EC | PLF |
| --- | --- | --- | --- | --- | --- | --- |
| 1 | *ATP2B3* | ENSG00000067842 | Yes | - Glioblastoma Multiforme (validation) | Yes | - *ATP2B3* encodes plasma membrane calcium ATPase 3 (PMCA3), a plasma membrane active transporter that mediates calcium efflux and contributes to intracellular calcium homeostasis (124).<br>- *ATP2B3* is highly expressed in neuronal tissues, including cerebellar neurons. It is expressed at presynaptic terminals, particularly in granule cells, cerebellar glomeruli, and Purkinje fiber terminals (128), and this expression was also confirmed using the HPA database.<br>- Germline mutations in *ATP2B3* have been linked to X-linked cerebellar ataxia and neurodevelopmental disorders (124, 129).<br>- somatic mutations in *ATP2B3* have been identified in aldosterone-producing adenomas, implicating altered calcium signaling in the pathogenesis of the disease (130). |
| 2 | *CACNA1I* | ENSG00000100346 | No | - Thyroid Carcinoma (TCGA)<br>- Breast Invasive Carcinoma (validation)<br>- Glioblastoma Multiforme (validation)<br>- Testicular Germ Cell Tumor (TCGA) | Yes | - *CACNA1I* encodes a T-type voltage-gated calcium channel (Cav3.3) that allows passive calcium influx down its concentration gradient (125, 131).<br>- Variants in *CACNA1I* have been associated with neurodevelopmental disorders, including epilepsy and cognitive impairment (125, 131).<br>- *CACNA1I* mRNA is largely restricted to the brain, while CaV3.3 protein expression is highly cell-type specific, with predominant functional expression in GABAergic neurons of the thalamic reticular nucleus (132, 133).<br>- *CACNA1I* has previously been proposed as a potential therapeutic target for specific cancer subtypes due to its overexpression in<br>cancers such as breast cancer relative to normal tissues (127). |
| 3 | *DCHS2* | ENSG00000197410 | No | - Glioblastoma Multiforme (validation)<br>- Pancreatic AC (validation)<br>- Breast Invasive Carcinoma (validation) | Yes | - *DCHS2* encodes a cadherin-related transmembrane protein involved in cell–cell adhesion (NCBI Gene; GeneCards).<br>- Primary literature on *DCHS2* is limited; however, a genome-wide association study has implicated *DCHS2* in Alzheimer's disease (134). |
| 4 | *DSEL* | ENSG00000171451 | No | - Glioblastoma Multiforme (validation)<br>- Glioblastoma Multiforme (TCGA)<br>- Breast Invasive Carcinoma (validation)<br>- Kidney Renal Clear Cell Carcinoma (validation)<br>- Kidney Renal Clear Cell Carcinoma (TCGA) | Yes | - *DSEL* encodes dermatan sulfate epimerase 2 (DS-epi2), an enzyme involved in glycosaminoglycan biosynthesis that contributes to extracellular matrix organization (135).<br>- Although *DSEL* mRNA expression and enzymatic function have been well characterized, detailed information on its endogenous cellular and subcellular localization remains limited (135, 136).<br>- Primary literature on *DSEL* is also limited; however, it has been detected in various screens (137, 138).<br>- *DSEL* has been shown to be increased in nasopharyngeal carcinoma compared to non-tumor samples (139). |
| 5 | *LPAR4* | ENSG00000147145 | No | - Glioblastoma Multiforme (validation)<br>- Glioblastoma Multiforme (TCGA) | Yes | - *LPAR4* encodes lysophosphatidic acid receptor 4 (also known as LPA4 & GPR23), a G protein–coupled receptor that binds the bioactive lipid lysophosphatidic acid (LPA) and mediates intracellular signaling through |
|  |  |  |  | - Breast Invasive Carcinoma (validation) |  | calcium mobilization and downstream pathways (140, 141).<br>- Through these signaling mechanisms, *LPAR4* regulates key cellular processes including cell migration, adhesion, and proliferation, thereby contributing to tissue organization and cellular dynamics (142, 143).<br>- *LPAR4* activity has also been implicated in cancer biology, where its expression can promote stress tolerance, tumor initiation, and drug resistance in certain tumor contexts (144, 145).<br>- *LPAR4* exhibits variable, tissue-restricted RNA expression, as detected by Northern blot and RT-PCR analyses, with transcripts detected in selected tissues rather than uniformly high expression across all organs (146).<br>- A recent primary study demonstrated that *LPAR4* expression is developmentally regulated, with dynamic RNA and protein expression in the nervous system. In the developing mouse brain, endogenous *LPAR4* protein was detected by Western blot and localized by immunohistochemistry to neural cells in defined regions, including the intermediate zone and cortical plate, during embryonic and early postnatal stages (147). |
| 6 | *LRRC15* | ENSG00000172061 | No | - Breast Invasive Carcinoma (validation)<br>- Breast Invasive Carcinoma (TCGA)<br>- Lung AC (validation)<br>- Lung SQCC (validation) | Yes | - *LRRC15* encodes leucine-rich repeat-containing protein 15, a type I transmembrane protein involved in cell–cell and cell–extracellular matrix interactions (148).<br>- *LRRC15* is proposed as a novel mesenchymal protein and stromal target (149). |
|  |  |  |  | - Colon AC<br>(validation) |  |  |
| 7 | *PRRT4* | ENSG00000224940 | No | - Testicular Germ Cell Tumor (TCGA)<br>- Glioblastoma Multiforme (TCGA)<br>- Glioblastoma Multiforme (validation)<br>- Skin Cutaneous Melanoma (TCGA)<br>- Breast Invasive Carcinoma (validation) | Yes | - *PRRT4* encodes a proline-rich transmembrane protein, although its specific biological function and role in cellular processes remain largely uncharacterized (NCBI Gene; GeneCards).<br>- Consistent with this, only a limited number of primary studies (n = 4) have investigated *PRRT4*.<br>- In one study, *PRRT4* was identified as part of an eight-gene prognostic model in acute myeloid leukemia, contributing to a risk signature associated with patient outcomes (150).<br>- *PRRT4* has also been suggested as a potential prognostic biomarker in gastric cancer in prior studies (151).<br>- More recently, pan-cancer transcriptomic analyses have further implicated *PRRT4* as a potential therapeutic target and prognostic biomarker across multiple cancer types; however, these findings are based on association studies and lack functional validation (152). |
| 8 | *TMEM229A* | ENSG00000234224 | No | - Glioblastoma Multiforme (validation)<br>- Glioblastoma Multiforme (TCGA) | Yes | - *TMEM229A* encodes a predicted transmembrane protein; however, its biological function remains incompletely characterized. Notably, one study demonstrated that *TMEM229A* is downregulated in non-small cell lung cancer and functions as a tumor suppressor, with its overexpression inhibiting cell proliferation, migration, and invasion by inactivating the ERK signaling pathway (153). |
|  |  |  |  |  |  | - Consistently, a recent study identified a *TMEM229A* mutation (Q200del) with a protective role in lung adenocarcinoma, also linked to ERK modulation (154).<br>- Overall, *TMEM229A* remains poorly studied beyond lung cancer contexts. |
| 9 | *UNC5A* | ENSG00000113763 | No | - Breast Invasive Carcinoma (validation)<br>- Ovary Serous Cystadenocarcinoma (TCGA)<br>- Glioblastoma Multiforme (TCGA)<br>- Ovary Serous Cystadenocarcinoma (validation)<br>- Breast Invasive Carcinoma (TCGA) | Yes | - Similar to *DCC*, *UNC5A* belongs to the family of netrin-1 dependence receptors, which act as context-dependent regulators of cell fate by promoting survival in the presence of ligand via inducing apoptosis in its absence (155).<br>- RNA-level analyses using in situ hybridization and RT-PCR confirm *UNC5A* mRNA expression in both neuronal and non-neuronal cells of the adult rat spinal cord, with down-regulation following injury (156). |
| 10 | *UNC79* | ENSG00000133958 | No | - Glioblastoma Multiforme (validation)<br>- Glioblastoma Multiforme (TCGA) | Yes | - Primary studies demonstrate that *UNC79* is expressed in neuronal cells, where it functions as part of the NALCN channel complex (involved in regulating neuronal excitability), although comprehensive tissue-level expression profiles remain limited (157, 158).<br>- *UNC79* has not been well characterized in cancer, with no primary studies demonstrating a functional role in tumorigenesis. |

While publicly available single-cell data from PT in EwS are extremely limited, with most work conducted in cell lines and in PDX/CDX (cell line-derived xenograft) models, we wanted to validate our findings using an independent dataset. We were fortunate to identify single-cell RNA-sequencing (scRNA-seq) data of 18 primary EwS tissue samples from 11 patients (GEO accession GSE243347). Out of the 18 samples, only 9 EwS samples from 6 patients were not flow sorted using CD45 before sequencing (44). Nearly all 9 PT samples originate from primary bone sites, apart from 1 sample annotated as “rib/kidney” (Sup.Table 2). Using these 9 EwS samples, we performed analysis using the same Seurat-based pipeline and, for QC, used different thresholds, as these samples were single cell rather than single nucleus (Sup.Fig. 3A). Additionally, because of differences in sequencing strategies, we have used this 9 EwS PT scRNA-seq dataset as an independent reference alongside the 13 EwS snRNA-seq dataset to further confirm and validate findings. Post-QC, the scRNA-seq dataset had 2205 cells expressing 20,535,744 UMI counts (Sup.Fig. 3B). Harmony integration was required (Sup.Fig. 3C & D), and we also performed unbiased clustering resulting in 18 (0–17) clusters (Sup.Fig. 4A).

**Figure 3.**
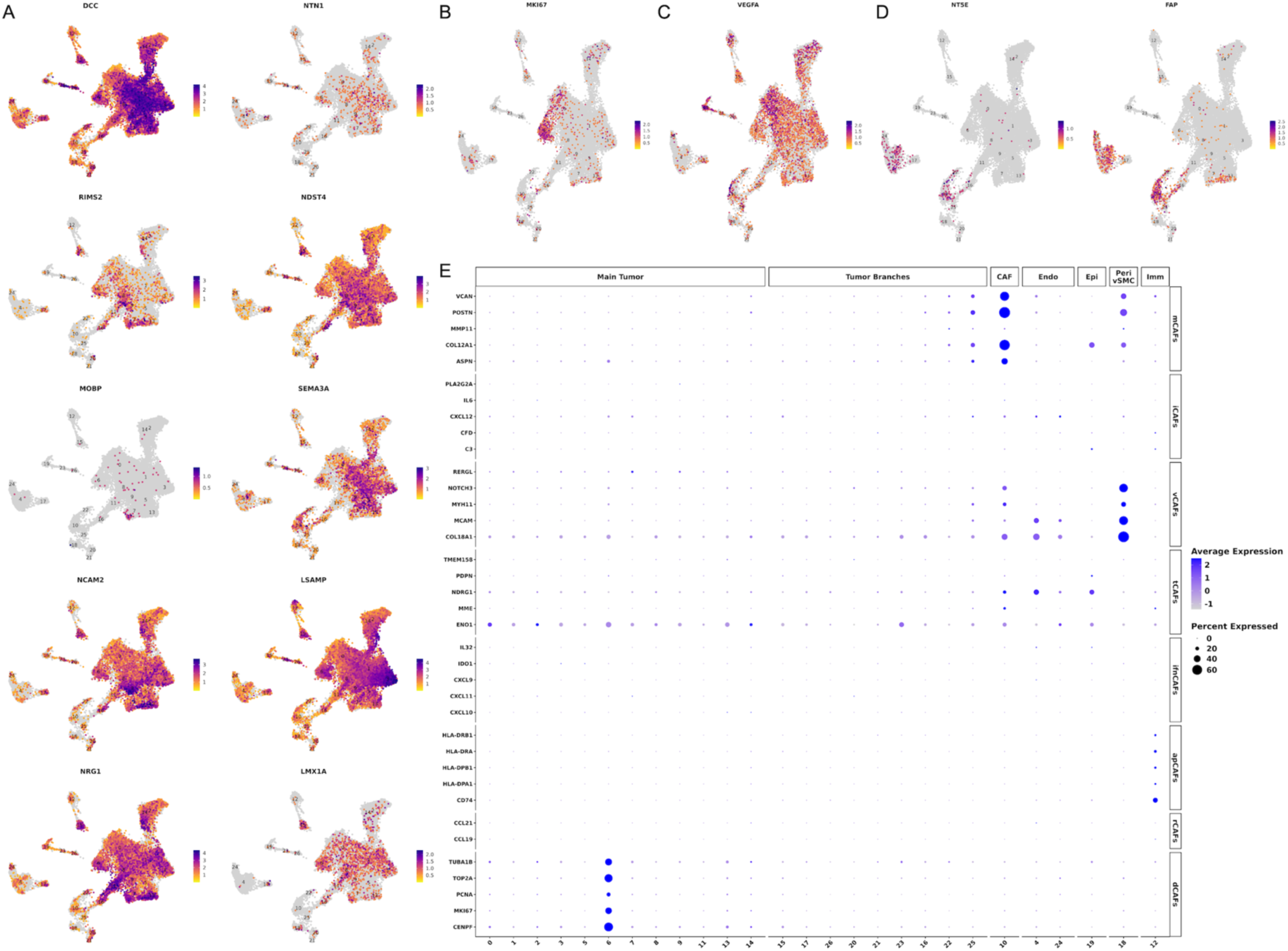
snRNA-seq gene expression profiles highlight dynamically neuronal-like, proliferative, angiogenic, and CAF-like cellular programs. (A - D) UMAP feature plots showing the distribution of selected gene expression per nucleus across clusters. High-scoring nuclei (purple) correspond to higher gene expression, while low-scoring nuclei (yellow) correspond to lower gene expression. (A) List of genes (*DCC*, *NTN1*, *RIMS2*, *NDST4*, *MOBP*, *SEMA3A*, *NCAM2*, *LSAMP*, *NRG1*, and *LMX1A*) expressed by the EwS tumor clusters, supporting a pervasive yet non-uniform neuronal-like transcriptional state. (B) Expression of *MKI67*, a marker for cell proliferation. (C) Expression of *VEGFA*, a marker of angiogenesis. (D) Expression of *NT5E* (CD73), a marker of EwS CAF-like cells, and *FAP*, a canonical marker of classical CAFs. (E) Dot plot showing expression of selected genes for the classification of CAF subpopulations (mCAFs = matrix; iCAFs = inflammatory; vCAFs = vascular; tCAFs = tumor-like; ifnCAFs = interferon-response; apCAFs = antigen-presenting; rCAFs = reticular-like; dCAFs = dividing) across clusters in different categories (Main Tumor, Tumor Branches, CAF, Endo, Epi, Peri/vSMC, and Imm) of the UMAP. In the dot plot, the y-axis right-hand side shows the list of genes, and the left-hand side shows the CAF subpopulation categories. The x-axis shows the cluster numbers from the UMAP projections. Dot size represents the percentage of nuclei expressing each gene, and color intensity indicates the average expression levels.

**Figure 4.**
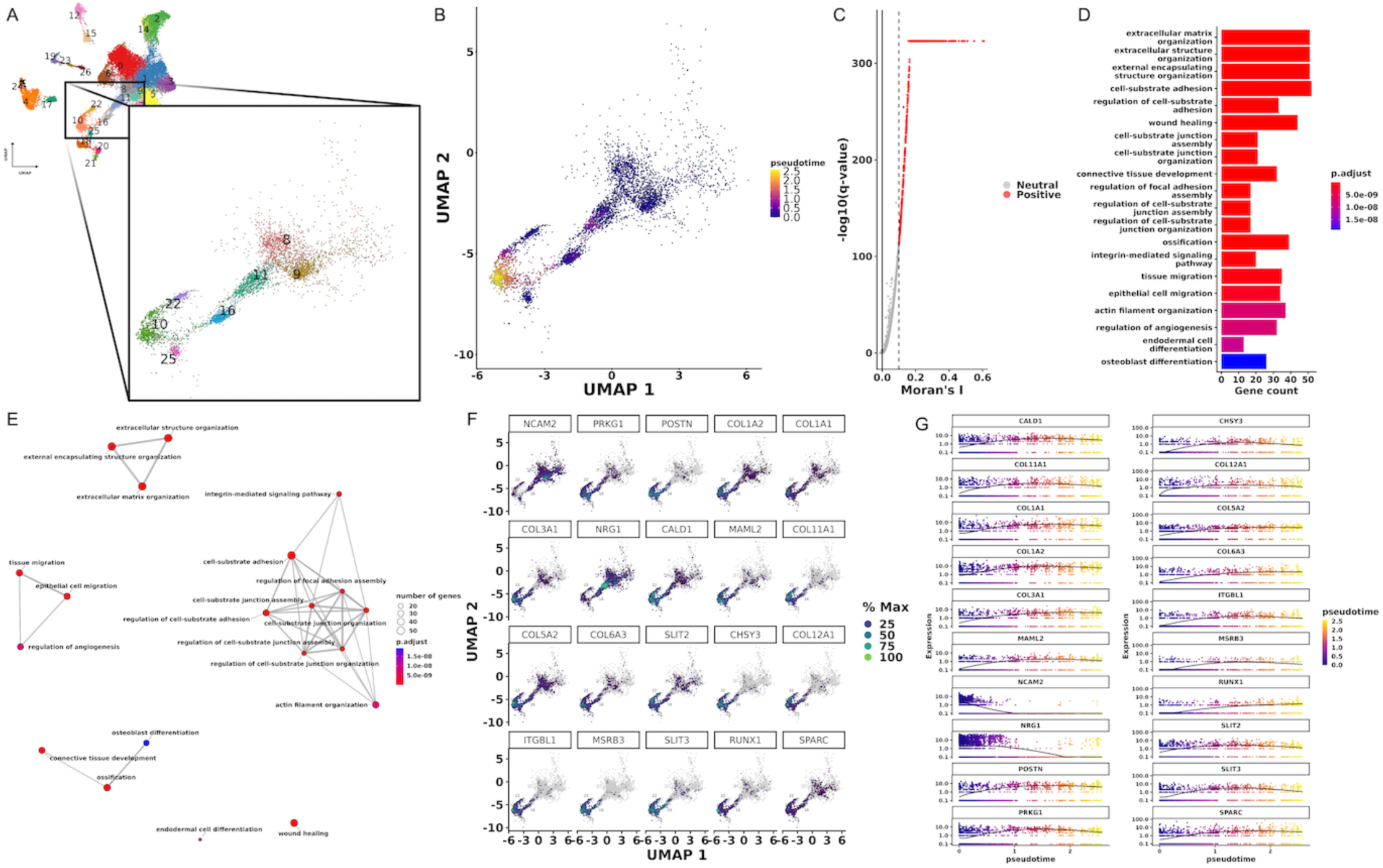
snRNA-seq trajectory analysis reveals ECM remodeling program along EwS–CAF Continuum. (A) UMAP illustration of the selected clusters for downstream trajectory analysis. (B) Monocle 3 pseudotime trajectory projected onto the UMAP embedding. Nuclei are colored by pseudotime, with early nuclei (purple) concentrated in the upper-right region and progressively increasing pseudotime values extending toward the central and lower-left branches (yellow). Trajectory inference was performed using reversed graph embedding in UMAP space, and nuclei were ordered using cluster 8 as the root. The continuous color gradient illustrates the inferred progression of cellular states along the branching trajectory structure in low-dimensional space. (C) Volcano-style scatter plot summarizing identified trajectory-associated genes on a learned principal graph after library-size normalization. Genes that showed significant variation along the trajectory are plotted by Moran’s I (x-axis), which measures how strongly expression follows the trajectory, and by statistical significance (−log10(q-value). Q-values were FDR-corrected, and zeros were replaced with a very small value to prevent infinite values during log transformation. The solid vertical line marks Moran’s I = 0, and the dashed vertical line marks Moran’s I = 0.1 cutoff, separating trajectory-structured genes (Positive; red) from neutral/weakly autocorrelated genes (Neutral; gray). (D) GO Biological Process enrichment analysis of trajectory-associated genes identified by Monocle 3 (Moran’s I > 0.1). Redundant GO terms were reduced using semantic similarity filtering (cutoff = 0.7). The top 20 enriched processes are shown on the y-axis, while the x-axis indicates the number of genes mapped to each term (gene count). The color scale represents the adjusted p-values (FDR). (E) Enrichment map network of the top 20 GO Biological Process terms derived from trajectory-associated genes (Moran’s I > 0.1). Pairwise semantic similarity between GO terms was calculated and terms were visualized as a network using a force-directed layout. Each node represents an enriched GO term; node size reflects the number of genes associated with the term, and node color indicates adjusted p-value (FDR). Edges connect terms with high semantic similarity, highlighting functional relationships and grouping related biological processes into clusters. (F) Feature plots showing expression of the top trajectory-associated genes (ranked by Moran’s I values) projected onto the UMAP embedding. Each panel displays one gene, with nuclei colored by % Max (scaled expression). On the Max % scale label, dark purple indicates low expression, while light green indicates high expression. (G) Gene expression dynamics across pseudotime for the top 20 trajectory-associated genes. Each panel shows normalized gene expression plotted against pseudotime, with nuclei colored by pseudotime (purple = early, yellow = late). The black line represents a smoothed trend curve summarizing the overall expression pattern across pseudotime. This visualization highlights genes that increase, decrease, or peak at specific stages along the inferred trajectory.

In both datasets, clusters were annotated using a multipronged strategy, beginning with reference-based cell-type prediction using the Human Primary Cell Atlas Data (a prebuilt single-cell reference dataset) (51). The prediction scores per nucleus/cell were averaged and visualized using heatmaps (row- and column-scaled), allowing systematic comparison of cluster-to-reference similarity patterns (Sup.Fig. 5A & B; Sup.Fig. 6A & B). This enabled the initial classification of some normal cell types based on their cell of origin. There is strong evidence that EwS tumors originate from mesenchymal stem cells (MSCs) (18), and based on our initial findings, it was much harder to distinguish tumor clusters from normal clusters with a mesenchymal origin (CAFs & vSMC) using only this approach. In contrast, immune, endothelial, and epithelial clusters were easier to identify because they exhibited stronger and more specific signatures for a particular reference cell type. To further classify clusters, we employed single-cell variational aneuploidy analysis (SCEVAN), a probabilistic framework that leverages gene expression profiles to infer copy-number alterations to identify tumor versus normal nuclei/cells (52). While genomically EwS is stable with very few secondary mutations driving tumorigenesis, copy number alterations (CNVs) have been reported in a subset of patients and can be used as a complementary approach to further define tumor and normal clusters (53, 54). Within the ALSF snRNA-seq dataset, CNVs were detected in 1 out of the 5 patients (Sup.Fig. 5C). In contrast, the GSE243347 scRNA-seq dataset showed CNVs in 4 out of the 6 patients, with varying degrees of alteration (Sup.Fig. 6C). Quantifying the distribution of CNV-positive cells within each cluster enabled the identification of clusters with elevated copy-number changes, consistent with regions in the tumor clusters (Sup.Fig. 5D; Sup.Fig. 6D). The scRNA dataset showed higher sensitivity for detecting CNVs than the snRNA dataset, which had more nuclei filtered out and not classified as normal or tumor. This was expected as SCEVAN is specifically designed for and validated on scRNA-seq data. Together, these findings highlight the challenge of distinguishing EwS from other mesenchymal cell types, and our initial methodology establishes a framework for elucidating tumor versus normal clusters.

**Figure 5.**
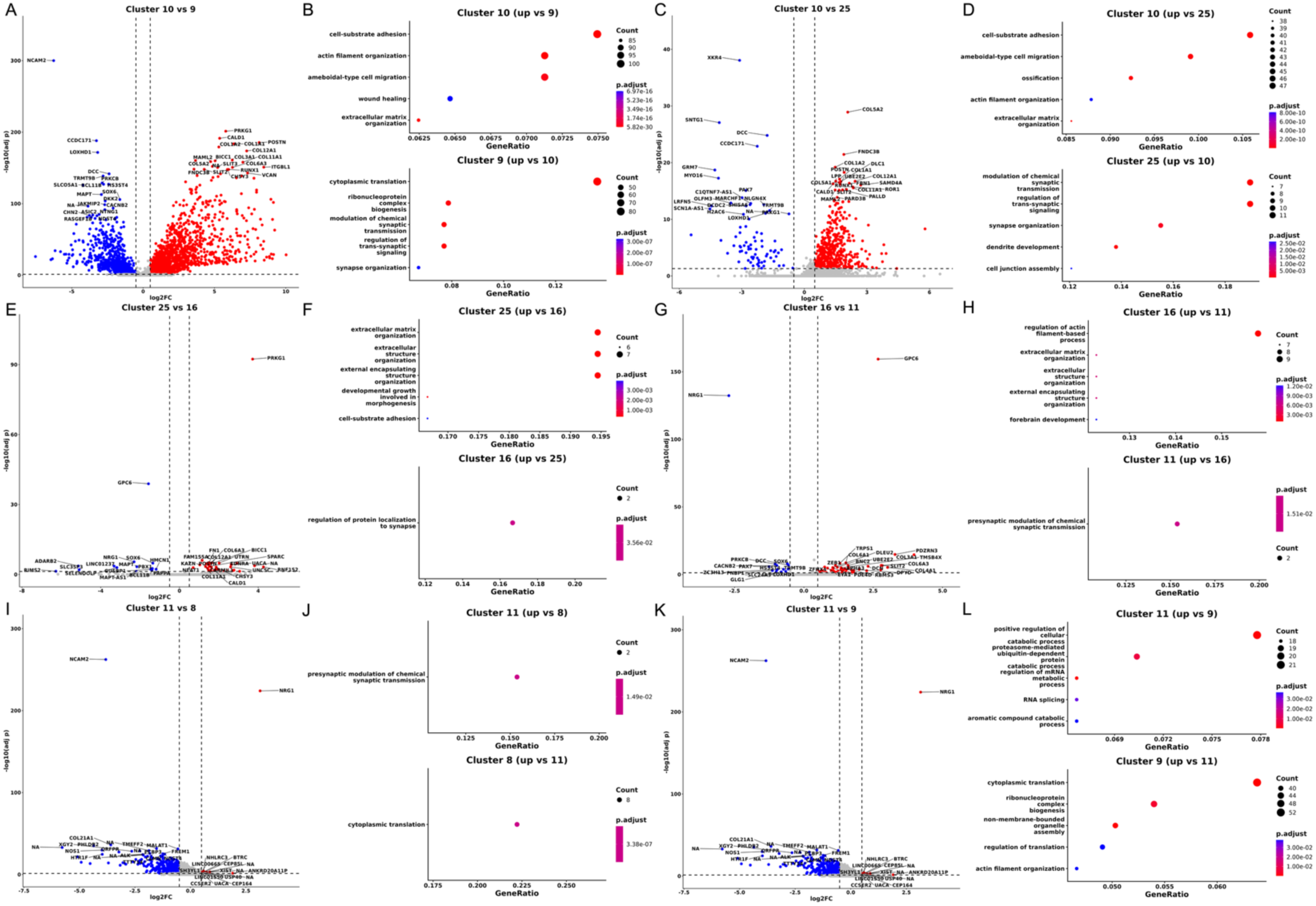
snRNA-seq Pairwise differential expression and GO enrichment analysis reveal functional differences between CAF-related clusters. Comparing transcriptional profiles of (A & B) 10 vs 9, (C & D) 10 vs 25, (E & F) 25 vs 16, (G & H) 16 vs 11, (I & J) 11 vs 8, (K & L) 11 vs 9 clusters. (A, C, E, G, I, and K) Volcano plot showing differential gene expression between selected 1-vs-1 clusters. Genes are plotted by log2 fold change (x-axis) and −log10 adjusted p-value (y-axis). Significantly upregulated genes in the initial cluster are shown in red, downregulated genes in blue, and non-significant genes in gray. Dashed vertical lines indicate log2FC cutoffs (±0.5), and the dashed horizontal line marks the adjusted p-value cutoff (FDR < 0.05). (B, D, F, H, J, and L) GO Biological Process enrichment analysis for genes upregulated in the initial (top) and second (bottom) clusters. The x-axis shows GeneRatio, dot size represents gene count, and color indicates adjusted p-value (FDR). Across all comparisons, differential expression analysis was performed using the Wilcoxon rank-sum test in Seurat, followed by GO Biological Process enrichment using clusterProfiler with Benjamini–Hochberg FDR correction. GO terms were ranked by GeneRatio, and the top enriched categories are shown for each comparison.

**Figure 6.**
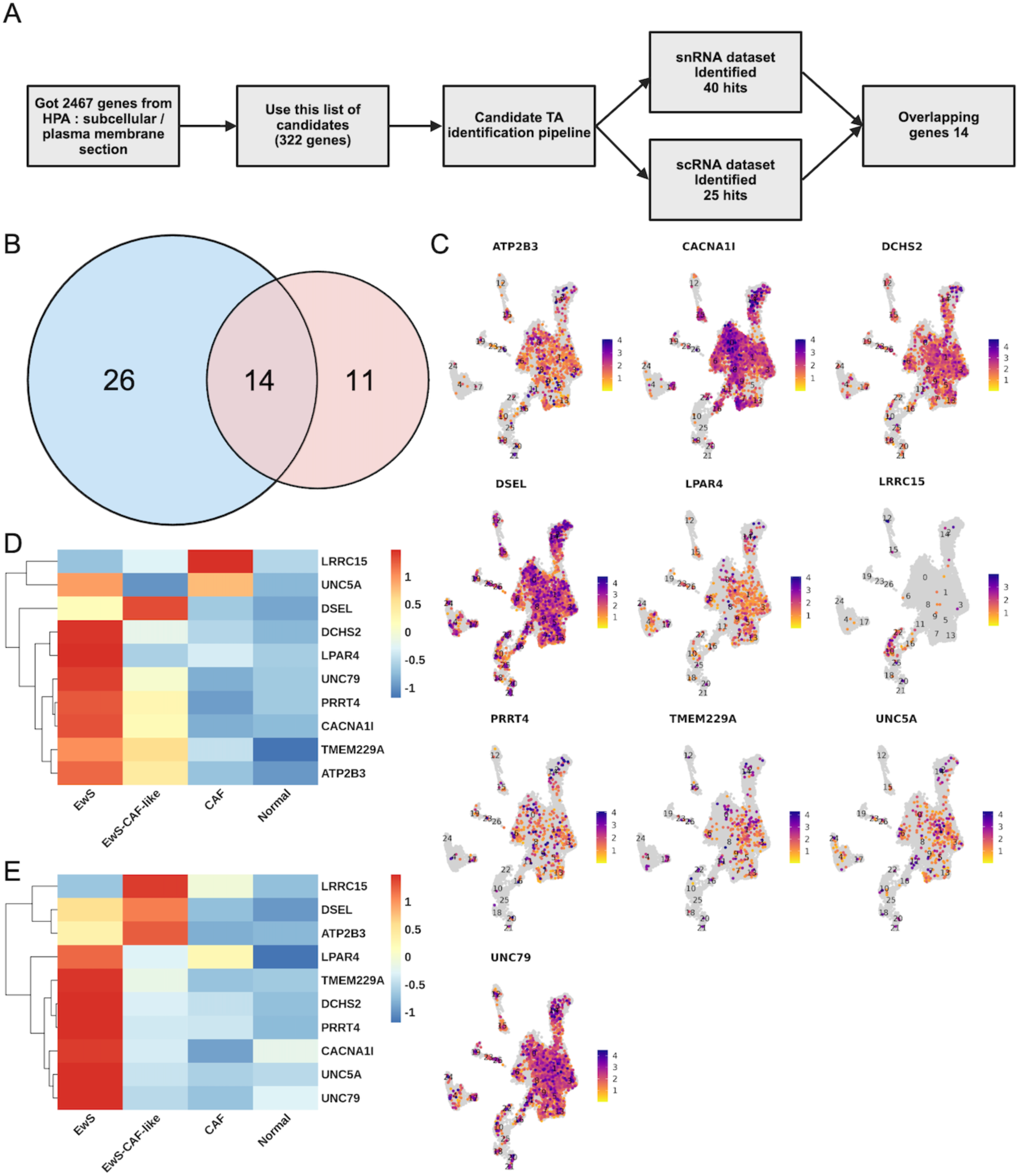
Integration of HPA, scRNA-seq, and snRNA-seq identifies top 10 putative surface EwS TAs in PT samples. (A) Schematic illustration of the TAs candidate identification pipeline. A total of 2,467 genes annotated in the Human Protein Atlas (HPA) as plasma-membrane or subcellular proteins were used as input. The pipeline identified 40 candidate hits in the snRNA-seq dataset and 25 candidate hits in the scRNA-seq dataset. (B) Venn diagram showing the overlap between candidate surface TAs identified in the snRNA-seq (blue) and scRNA-seq datasets (pink). A total of 14 genes were shared between the two datasets, while 26 genes were unique to snRNA-seq and 11 genes were unique to scRNA-seq. (C) UMAP feature plots showing the expression patterns of the top 10 putative EwS TAs across PT samples in the integrated snRNA dataset. Each panel shows normalized RNA expression for the selected gene across nuclei clusters, with purple color intensity indicating relative expression levels. (D & E) Heatmap showing the average expression of the top 10 putative EwS TAs across major cell categories (EwS, EwS CAF-like, CAF, and Normal) in the (D) snRNA-seq or (E) scRNA-seq integrated dataset. Average values were calculated from normalized RNA expression and scaled as row-wise Z-scores. The heatmap highlights the relative enrichment of the candidate TA, where red indicates higher enrichment within a particular category.

In addition to reference-based annotation and SCEVAN analyses, we further examined each cluster in both datasets independently using gene set enrichment analysis, unsupervised hierarchical clustering, differential gene expression (DEG) analysis, and evaluation of the top-expressed genes per cluster. For the gene set enrichment analysis, we used 11 gene sets: 9 published in MSigDB and 2 (gene sets 4 and 6) curated in-house via publicly available literature reviews for CAF and EwS diagnostic markers, respectively (Sup.Table 3). Gene sets 1 (37 genes (55)) and 2 (439 genes (56)) represent targets of the EWSR1::FLI1 fusion that are upregulated by the fusion, which overlapped 43.57% and 3.19% respectively (Sup.Table 3). Both gene sets 1 (Fig. 1B/2F; Sup.Fig. 4B/7F) and 2 (Fig. 2C/2F; Sup.Fig. 7C/7F) show similar expression patterns across clusters and reinforce our initial classifications of tumor vs. normal clusters. Tumor clusters within the main tumor body showed higher scores for both gene sets 1 and 2 (Fig. 2F; Sup.Fig. 7F), consistent with activation of EWSR1::FLI1 target genes. We collectively termed this measure EwSness scores, reflecting the degree to which nuclei/cells exhibit EwS-associated gene signatures in either gene set 1 or 2. Since EwS cells exist along a transcriptional continuum characterized by cell-to-cell variation in EWSR1::FLI1 expression and downstream activity (33), all clusters were ranked according to their gene set 1 EwSness scores and divided into three equally sized groups representing High, Middle, and Low. Gene set 1 was selected for this classification because it represents a more discrete EwS-associated gene signature, whereas gene set 2 comprises a broader collection of genes. Within the snRNA dataset, cluster 14 exhibited the lowest EwSness score among all tumor clusters (Fig. 2F). In the scRNA dataset, cluster 5 had the lowest EwSness score among tumor clusters, followed by cluster 7 (a B cell cluster), which showed substantially lower scores across both gene sets 1 and 2 (Fig. 7F). Gene set 6, comprising of a curated panel of 7 EwS diagnostic markers, further supported our classification of tumor versus normal clusters (Fig. 2F; Sup.Fig. 7F). Comparing results from snRNA and scRNA revealed that EwS diagnostic markers (gene set 6) more effectively supported the identification of tumor clusters in the scRNA dataset. Notably, cluster 5 in scRNA exhibited a substantially higher score than the B cell cluster 17 and exceeded the score for cluster 8, which is definitively classified as an EwS tumor cluster (Sup.Fig. 7F). The individual expression levels of these 7 diagnostic markers (*CD99*, *NKX2-2*, *PRKCB*, *GLG1*, *BCL11B*, *CAV1*, and *NR0B1* (57–63)), along with other EwS-associated genes *KDSR* (64), *MAPT* (65), and *STEAP1* (66), further highlighted the distinction between tumor and normal clusters in both datasets (Fig. 1F; Sup.Fig. 4F).

**Figure 7.**
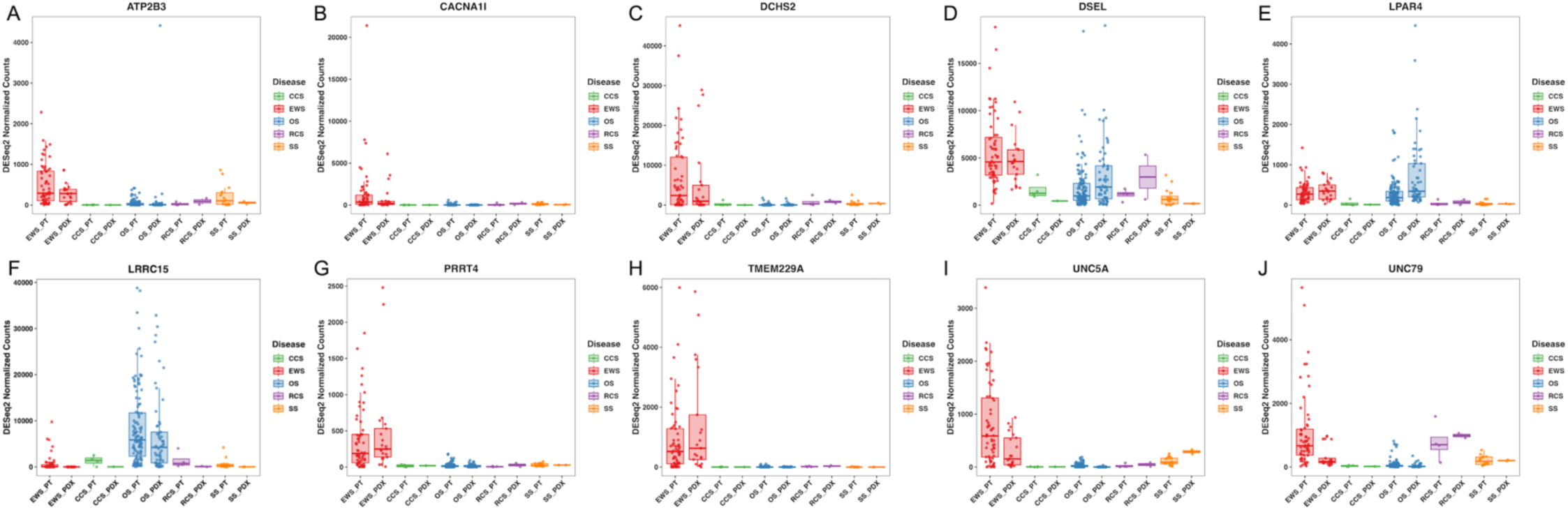
Expression of 10 top putative EwS surface TAs in St. Jude Bulk RNA 283 samples. (A - J) DESeq2-normalized bulk RNA-seq expression of (A) *ATP2B3*, (B) *CACNA1I*, (C) *DCHS2*, (D) *DSEL*, (E) *LPAR4*, (F) *LRRC15*, (G) *PRRT4*, (H) *TMEM229A*, (I) *UNC5A*, and (J) *UNC79* across cancer types; EwS (red), Clear Cell Sarcoma (CCS; Green), OS (blue), Round Cell Sarcoma (RCS; purple), and Synovial Sarcoma (SS; orange). Each disease type was further split by PT or PDX status, see sup Sup.Table 3 for full metadata. Boxplots summarize the distribution of expression values across samples and demonstrate variable enrichment of these candidate TAs among sarcoma subtypes, including EwS PT (60) and EwS PDX (21)samples.

Unsupervised hierarchical clustering using the first 20 principal components (PCs) was performed to further distinguish tumor from normal cells and to provide additional insight for cluster annotations. In the snRNA dataset, tumor and normal clusters separated clearly (Fig. 1D). In contrast, in the scRNA dataset, cluster 5, characterized by a Middle-EwSness and the presence of CNVs, was transcriptionally similar to cluster 15, which represents the normal CAF (classical CAF) cluster (Sup.Fig. 4D). This was not unexpected, as EwS cells are known to have the ability to undergo transcriptional shifts toward CAF-like subpopulations (28), and recent scRNA-seq analyses have shown that EwS tumor clusters can transcriptionally separate from the main tumor body and cluster adjacent to CAFs (67). By examining the top 100 genes in each cluster (Fig. 2A & B; Sup.Fig. 7A & B) and the top 10 (DEGs) when comparing all clusters to one another (Fig. 2D & E; Sup.Fig. 7D & E), we again identified transcriptional patterns that distinguish tumor from normal clusters. Here, we observe that in the scRNA dataset, when unsupervised hierarchical clustering is performed on the top 100 genes, cluster 5 groups with cluster 8, a tumor cluster with almost a similar EwSness score, and is positioned further from cluster 15, which represents classical CAFs (Fig. 4B). Furthermore, examination of the top 10 DEG per-cluster heatmap demonstrates that cluster 5 exhibits a distinct transcriptional profile compared to clusters 15 and 8, suggesting that it represents a transitional population bridging the tumor to classical CAFs (Fig. 4D & E). This observation is consistent with cluster 25 in the snRNA-seq dataset, which similarly exhibited both tumor and CAF-like transcriptional signatures (Fig. 2A–D).

Using these analytical approaches in combination, we successfully annotated both datasets and distinguished tumor from normal clusters, including classical CAF populations (snRNA = 10; scRNA = 15) and vascular smooth muscle (vSMC; snRNA = 18; scRNA = 7) clusters, which EwS is believed to share a mesenchymal origin with (Fig. 1E; Sup.Fig. 4E). Epithelial cells were identified only in the snRNA-seq dataset, and cluster 19 was primarily derived from metastatic lung samples (Fig. 1E). Based on close inspection and the expression of surfactant-associated markers such as SFTPB and SFTA3, these cells were annotated as alveolar epithelial cells (68). In both datasets, immune cell populations were successfully annotated; however, in the snRNA, all immune cells clustered into a single cluster (cluster 12), whereas in the scRNA we had clusters 2 (T cells), 3 (NK cells), 4 (monocytes/dendritic cells), 13 (monocytes), 16 (mast cells), and 17 (B cells). Increasing the cluster resolution within the scRNA dataset further subdivided the immune populations but did not affect the composition or identity of cluster 5 (Fig. 1E; Sup.Fig. 4E), suggesting insufficient transcriptional divergence to robustly resolve this cluster into two distinct subpopulations. Cluster 5, in scRNA, is similar to cluster 25, in snRNA, as both tumor-associated clusters exhibit transcriptional profiles closely resembling those of classical CAF populations. Overall, we show that both EwS datasets exhibit substantial heterogeneity across tumor clusters and highlight the complexity of distinguishing tumor from normal clusters with similar cell of origin.

### Diverse and Multidirectional Transcriptional Heterogeneity in EwS

Examining the EwS intratumoral landscape revealed 12 clusters comprising of the main tumor (0, 1, 2, 3, 5, 6, 7, 8, 9, 11, 13, and 14) and 9 clusters termed tumor branches (15, 17, 26, 20, 21, 23, 16, 22, and 25) at the snRNA level (Fig. 1E). Similarly, at the scRNA level we observed both main tumor clusters (0, 1,9, 10, and 11) and tumor branches (14, 12, 8, 5) (Sup.Fig. 4E). The reduced separation of tumor clusters in snRNA compared to scRNA is most likely a reflection of biological differences in transcript capture. scRNA-seq profiles whole-cell RNA, including cytoplasmic transcripts, and thus often provides greater resolution of dynamic cell states. While snRNA-seq only captures nuclear RNA, which contains less total RNA, thereby reducing transcriptional complexity and obscuring distinctions between closely associated tumor subpopulations (69). Nonetheless, given the limited availability of high-resolution EwS transcriptomic datasets, integrating scRNA-seq and snRNA-seq provides a complementary framework for cross-validation, enabling a more robust characterization of tumor heterogeneity, TME, and the identification of candidate TAs.

In both datasets, EwS tumor clusters consistently exhibited a pronounced neuronal-like transcriptional signature compared with normal cell populations, although the expression of neuronal markers did vary across tumor clusters (Fig. 1G; Sup.Fig. 4G). Notably, tumor cells expressed genes involved in axon guidance and neuronal signaling, including *DCC*, a netrin-1 receptor that can function as a dependence receptor inducing apoptosis in the absence of its ligand (NTN1), but promoting tumor survival when netrin-1 is present (70, 71). Within the TME transcriptional landscape both *DCC* and *NTN1* are expressed, possibly promoting tumor cell survival and TME-dependent growth (Fig. 3A; Sup.Fig. 8A). Neuronal signaling, including axon-guidance programs, is associated with immune suppression, tumor progression, and the acquisition of stem-like states (72, 73). *SEMA3A* was enriched in tumor clusters across both datasets (Fig. 3A; Sup.Fig. 8A), is a secreted axon-guidance molecule that suppresses anti-tumor immunity by impairing CD8⁺ T-cell migration and function through cytoskeletal disruption (74). Additional neuronal adhesion and migration-associated genes such as *LSAMP* (75, 76), *NCAM2* (77), *DLG2* (78), and *ROBO2* (79) were detected, suggesting roles in cell adhesion, synaptic organization, and invasive behavior (Fig. 1G; Sup.Fig. 4G). Tumor clusters also displayed enrichment of synaptic and neurotransmission-related genes, including *GRM8*, *SNAP25*, *SYT1*, and the *NRG1–ERBB4* signaling axis, which is implicated in neuronal differentiation, survival, and tumor–TME interactions (Fig. 1G; Sup.Fig. 4G) (80–83). Genes associated with cytoskeletal dynamics and neurite outgrowth, such as the axonal growth regulators *STMN2* and *GAP43* (84–86), and intracellular fibroblast growth factors *FGF13* and *FGF14* (87–89), further supported a neuronal-like cellular state in PT samples. Additional markers, including UCHL1 (90–92), *SNTG1* (93), *XKR4* (94), and *TRMT9B* (95) (listed only in snRNA dataset), reinforced the presence of neuronal maintenance and differentiation programs within tumor cells (Fig. 1G; Sup.Fig. 4G). Neuronal-associated markers such as the *MOBP* gene, a classical marker of mature myelinating oligodendrocytes, were detected at low levels (96, 97). In contrast, genes including *RIMS2* (98), *NDST4* (99), and *NRG1* (100) were expressed at substantially higher levels and exhibited distinct distribution patterns across tumor clusters, suggesting their contribution to the pronounced intratumoral heterogeneity observed in EwS tumor subpopulations (Fig. 3A; Sup.Fig. 8A). We also identified several neuronal identity genes, including master transcriptional regulators such as *LMX1A*, which is involved in neuronal lineage specification and dopaminergic neuron differentiation (101, 102), further supporting the presence of a coordinated neuronal-like transcriptional program within EwS tumor clusters (Fig. 3A; Sup.Fig. 8A). There is substantial evidence that EwS tumors express neuronal-like transcriptional programs, including resemblance to neural crest stem cells (103–105). There is evidence linking the EwS fusion EWSR1::FLI1 to neuronal gene programs (65, 106); however, this relationship is complex, as the fusion expression varies from cell-to-cell and functions as both a transcriptional activator and repressor of various genes targets (107).

Beyond these neuronal-like signatures, EwS tumor clusters diverged into multiple transcriptional trajectories, reflecting distinct functional states. Previous studies have shown that variation in the intensity of EWSR1::FLI1 across individual cells represents a major source of heterogeneity in EwS tumors, and is associated with differences in proliferative, migratory, and metabolic cellular states (27, 67). In both datasets, a prominently proliferative cluster was identified (snRNA = 6; scRNA = 14), characterized by high expression of cell-cycle associated genes, including *MKI67* (Fig. 3B; Sup.Fig. 8B). Consistent with previous reports indicating elevated replication stress in EwS (108), this proliferative subpopulation exhibited transcriptional features indicative of replication stress, including activation of DNA damage response and cell cycle checkpoint genes (Fig. 2F; Sup.Fig. 7F). Also, *VEGFA*, a key marker of angiogenesis, was detected across multiple tumor clusters (Fig. 3C; Sup.Fig. 8C). We were also able to identify a distinct *VEGFA*-high cluster (snRNA = 0; scRNA = 8), indicative of a pro-angiogenic tumor subpopulations in PT samples. This observation is consistent with prior studies demonstrating that angiogenesis can be driven by specific tumor cell subsets, further highlighting the role of intratumoral heterogeneity in promoting tumor progression in EwS (35, 109). Additionally, some EwS subpopulations, such as CD73+ cells, exhibit features typically associated with CAFs (28). In particular, CD73+ EwS cells upregulate mesenchymal lineage genes normally suppressed by the EWS::FLI1 oncogenic program, thereby acquiring a hybrid, more stromal-like phenotype (28). These CAF-like tumor cells express ECM genes and actively contribute to TME remodeling by depositing structural proteins such as collagens. CD73+ EwS cells were also observed along the tumor border and invasive fronts, suggesting a potential role in tumor progression. Interestingly, the density of these CAF-like tumor cell subpopulations varies across cell lines (28), possibly reflecting intrinsic differences between EwS tumor cells as well as extrinsic cues that shape cellular plasticity and transcriptional heterogeneity. Here, we show that in PT samples, several tumor clusters shift transcriptionally toward a CAF-like state in both datasets (snRNA = 16, 22, 25; scRNA = 8, 5) (Fig. 1A; Sup.Fig. 4A). These clusters had higher scores for gene set 3, which comprises of 195 genes downregulated by EWSR1::FLI1 (Fig. 1C/2F; Sup.Fig. 4C/7F) and for gene set 5, which comprises of 200 hallmark epithelial mesenchymal transition (EMT) genes (Fig. 2C/2F; Sup.Fig. 7C/7F). Also, FAP, a canonical CAF marker (110, 111), was expressed at lower levels in these CAF-like clusters and even less in some tumor clusters within the main tumor mass, highlighting the tumors’ heterogeneity in expressing CAF-associated genes (Fig. 1H; Sup.Fig. 4H). Furthermore, not all CD73+ tumor cells expressed FAP, further increasing transcriptional variation in the expression of mesenchymal-associated genes within tumor cells (Fig. 3D; Sup.Fig. 8D). Taken together, these findings highlight that EwS PT cells are not defined by a single uniform transcriptional state but instead comprise a spectrum of dynamically interconverting cellular programs spanning neuronal-like, proliferative, angiogenic, and CAF-like phenotypes. This multidirectional heterogeneity reflects both intrinsic variability in EWSR1::FLI1 activity and extrinsic influences from the TME, collectively shaping tumor behavior and adaptability. More importantly, the coexistence of these diverse transcriptional states within individual tumors underscores the challenge of targeting EwS with single-agent therapies and emphasizes the need for therapeutic strategies that account for this cellular plasticity.

### Trajectory analysis Reveals ECM Remodeling Programs Along EwS-CAF continuum

In this study, we show that the EwS TME contains both classical CAFs (snRNA = 10; scRNA = 15) and CAF-like tumor cell populations (snRNA = 16, 22, 25; scRNA = 8, 5). In both datasets, analysis of EwSness scores revealed that EwS CAF-like clusters exhibited higher scores than classical CAFs and other nonmalignant cells but lower than those of main tumor clusters. Using gene sets 4 (CAF Markers; *FAP*, *PDPN*, *PDGFRB*, *ACTA2*, *COL1A1*, *THY1*, *TNC*, *SPARC*, *LUM* (112–115)) and 5 (Hallmark EMT genes), we observed upregulation of mesenchymal genes in EwS CAF-like clusters compared to the main tumor clusters (Fig. 2F; Sup.Fig. 7F). Additionally, classical CAF clusters exhibited higher CAF gene signatures than EwS CAF-like clusters (Fig. 1H; Sup.Fig. 4H), whereas EwS CAF-like clusters showed increased neuronal background signatures relative to classical CAFs but lower than those from the main tumor clusters (Fig. 1G; Sup.Fig. 4G). For example, *LSAMP*, a neuronal adhesion molecule whose loss has been implicated in EMT and enhanced cellular migration (116), was significantly enriched in main tumor clusters relative to EwS CAF-like clusters (Fig. 3A; Sup.Fig. 8A). To further characterize the CAF subpopulations present within the TME, we applied the CAF classification framework proposed by the Bodenmiller lab, which defines multiple transcriptionally and functionally distinct CAF phenotypes in a tumor-type–independent manner (117). Within the snRNA dataset, cluster 10 was identified as matrix CAFs (mCAFs) (Fig. 3E), whereas in the scRNA dataset, cluster 15 exhibited a mixed phenotype comprising of both mCAFs and inflammatory CAFs (iCAFs) (Sup.Fig. 8E). Notably, EwS CAF-like clusters across both datasets predominantly expressed mCAF-associated transcriptional programs, indicating a shift toward matrix-remodeling phenotypes. Collectively, these results support a model in which PT samples cells acquire stromal-like transcriptional programs without fully mirroring classical CAF identity, highlighting the plasticity of EwS tumor cells within the TME.

Interestingly, in both datasets, we observed multiple EwS CAF-like transcriptional states that progressively transition from main tumor clusters toward a CAF-like state, characterized by decreasing neuronal signatures and increasing mesenchymal signatures. For example, in the snRNA dataset, we observe a bridge region connecting the tumor (clusters 8, 9, 11) to the classical CAFs (cluster 10) via EwS CAF-like subpopulations (clusters 16, 25, 22) (Fig. 1A). However, this transitional bridge region was not observed in the scRNA dataset (Sup.Fig. 4A), as previously mentioned, likely due to its higher resolution. To test whether there is a transitional shift from tumor nuclei/cells to classical CAFs, we performed trajectory analysis using Monocle 3, which models transcriptional trajectories and infers dynamic cell-state transitions (118, 119). In the snRNA dataset, we subset two main tumor clusters (8 and 9), the classical CAF cluster (10), and intermediate clusters transcriptionally positioned between these states (11, 16, 22, and 25) (Fig. 4A). Trajectory analysis was performed using Monocle 3, which learns a principal graph to model the underlying structure of transcriptional space and reconstructs nuclei state transitions independent of predefined clusters. Cluster 8 was designated as the root for trajectory inference (Sup.Fig. 11A), and pseudotime was assigned by ordering nuclei along the learned principal graph according to their distance from the root (Fig. 4B). Testing all the genes, 513 showed significant variation along the trajectory (Moran’s I > 0.1), indicating that these genes are dynamically regulated along the inferred transition from tumor to CAF-like, to classical CAF states (Fig. 4C). Gene Ontology (GO) enrichment analysis of trajectory-associated genes (513 genes) revealed significant enrichment of extracellular matrix organization, cell–substrate adhesion, and integrin-mediated signaling pathways, indicating a shift toward matrix remodeling and stromal-like functions (Fig. 4D). Additional enriched processes included epithelial cell migration, actin filament organization, and tissue migration, further supporting the acquisition of migratory and invasive properties along the trajectory. While only the top 20 GO terms are reported, at number 29, response to TGF-β was also significantly (adjusted p-value of 8 × 10⁻⁷) enriched along the trajectory, with more than 30 genes assigned to this category, further implicating TGF-β signaling in the acquisition of CAF-like and matrix-remodeling features. Consistent with this, recent work demonstrated that EwS CAF-like cells occupy a hybrid transcriptional state in which derepression of *TGFBR2* and increased expression and secretion of TGFB2 sustain TGF-β pathway activation and extracellular matrix deposition, thereby stabilizing the EwS CAF-like phenotype (34). Enrichment map analysis demonstrated that these GO terms form highly interconnected networks, with extracellular matrix organization and cell adhesion processes clustering together, highlighting coordinated activation of mesenchymal and stromal gene programs (Fig. 4E). Notably, angiogenesis-related pathways also emerged within this network, suggesting that cells progressing along the trajectory may contribute to pro-tumorigenic remodeling of the TME. Feature plots of the top 20 trajectory-associated genes projected onto the UMAP embedding revealed distinct spatial expression patterns along the inferred trajectory, with neuronal-associated genes such as *NCAM2* (77) and *NRG1* (100) enriched in early tumor states (cluster 8 to 9), while extracellular matrix and mesenchymal genes, including *POSTN*, *COL1A1*, and *COL1A2* (117), were progressively upregulated in EwS CAF-like and classical CAF clusters (Fig. 4F). Consistent with these observations, analysis of gene expression dynamics across pseudotime demonstrated a clear shift in transcriptional programs, with neuronal-associated genes decreasing and mesenchymal and ECM-related genes increasing as cells progressed along the trajectory (Fig. 4G).

To validate the trajectory analysis findings, we performed cluster-to-cluster differential gene expression (DEG) analyses followed by GO enrichment analysis (Fig. 5). Comparing cluster 10 (classical CAFs) with cluster 9 (main tumor cluster) revealed significant transcriptional differences, with cluster 10 enriched for CAF-associated genes and cluster 9 characterized by neuronal-like programs (Fig. 5A & B). Comparison of cluster 10 with cluster 25 (a CAF-like cluster proximal to the classical CAFs) showed a marked reduction in neuronal programs in cluster 25, alongside increased enrichment of CAF-associated signatures (Fig. 5C & D). In contrast, comparison between cluster 25 and cluster 16 revealed relatively modest transcriptional differences; however, cluster 25 was more strongly characterized by CAF-like programs, whereas cluster 16 retained higher expression of neuronal genes (Fig. 5E & F). Progressing backward along the trajectory, cluster 16 showed enrichment for CAF-like programs, while cluster 11 was dominated by neuronal gene expression (Fig. 5G & H). Comparisons between cluster 11 and clusters 8 or 9 reveal limited transcriptional divergence, with clusters 8 and 9 exhibiting a more pronounced neuronal-like phenotype (Fig. 5I - L). Collectively, these findings support a model in which EwS tumor cells undergo coordinated transcriptional reprogramming toward stromal-like states, marked by progressive loss of neuronal identity and activation of CAF-associated programs.

To further assess this transcriptional shift from tumor to CAF-like to cells which closely resemble classical CAFs, we performed a trajectory analysis on the scRNA dataset, which yielded similar findings. We subset two main tumor clusters (10 and 1), the classical CAF cluster (15), and intermediate clusters transcriptionally positioned between these states (5, 14, and 8) (Sup.Fig. 9A). Trajectory analysis was performed using similar methods, and cluster 10 was designated as the root for trajectory inference. Interestingly, and likely due to the higher resolution of the scRNA dataset, we observed transcriptionally distinct EwS subpopulations that give rise to multiple branching trajectories. From cluster 10, the primary trajectory bifurcates into two sub-branches: one progressing through cluster 14 toward to cluster 8, and the other extending through cluster 5 toward the classical CAF cluster 15 (Sup.Fig. 11B). Notably, this branching pattern further highlights the transcriptional similarity between clusters 5 and 10 and the differences between clusters 5 and 8, despite both clusters 5 and 8 being classified as EwS CAF-like subpopulations. With pseudotime overlaid, cluster 15 occupied the most distal position from the root, underscoring the clear differences between tumor and the non-tumor classical CAFs (Sup.Fig. 9B). Consistent with the presence of multiple trajectories, including a trajectory from cluster 10 to the highly proliferative cluster 14, we identified a larger set of genes (n = 2,656) that dynamically change along the inferred trajectories (Sup.Fig. 9C). Several genes exhibited trajectory-specific expression patterns, including transient peaks or gradual activation, highlighting the complexity of interpreting gene expression changes along a single trajectory in the presence of multiple branching paths. Nonetheless, GO enrichment analysis of trajectory-associated genes (2,656 genes) further highlighted significant enrichment for extracellular matrix organization and cell–substrate adhesion (Sup.Fig. 9D & E). Feature plots of the top 20 trajectory-associated genes showed that most were CAF/ECM-related (Sup.Fig. 9F). These CAF/ECM-related genes showed progressive upregulation along the trajectory, with low expression in early tumor states and increasing expression toward CAF-like and classical CAF populations (Sup.Fig. 9G). DEG analysis also revalidated current findings and showed that both clusters 5 and 8 are EwS CAF-like clusters (Sup.Fig. 10). Collectively, using both snRNA and scRNA datasets, these findings support a model in which EwS tumor cells undergo coordinated transcriptional reprogramming toward multiple stromal-like states, marked by progressive loss of neuronal identity and activation of mesenchymal, extracellular matrix, and CAF-associated programs.

### Transcriptional Comparison of PT and Corresponding PDX Models

Gene expression in EwS tumors is highly context-dependent, with the EWSR1::FLI1 fusion governing the transcriptional landscape and responding dynamically to both the intracellular states and extracellular cues (34, 35). This raises important questions about how transcriptional programs differ between PT and matched O-PDX samples. To investigate this, we used the snRNA dataset, which, as previously described, includes 13 EwS samples from 5 patients, comprising of 7 PT and 6 matched O-PDX samples (Sup.Table 1). The 6 O-PDX samples were processed similarly but independently from the PT samples, and after initial processing of the dataset, we also performed unbiased clustering, which resulted in 14 (0–13) clusters (Sup.Fig. 12A). Gene set analysis revealed that gene sets 1 and 2 (EWSR1::FLI1-upregulated targets) showed diminished cluster-level consistency in EwSness scores between the two gene sets in the O-PDX samples compared with the PT samples, suggesting increased transcriptional variability in fusion-driven programs (Sup.Fig. 12B). In addition to gene set 3 (EWSR1::FLI1-downregulated targets), all gene sets demonstrated reduced activity, as evidenced by a decrease in the proportion of AUCell-active cells per cluster when comparing matched O-PDX to PT samples. Within the O-PDX tumor, cluster 13 was the most proliferative, followed by clusters 5 and 1 (gene set 7). These clusters similarly showed elevated ATR activation in response to replication stress (gene set 7) relative to all other tumor clusters within O-PDX samples (Sup.Fig. 12B). EwS, Neuronal, angiogenic, and CAF markers were detected within the O-PDX tumor clusters (Sup.Fig. 12C - I). However, the structured model in which EwS tumor cells undergo coordinated transcriptional reprogramming toward stromal-like states, marked by progressive loss of neuronal identity and activation of CAF-associated programs, was not observed. The expression patterns of transcriptional programs at the cluster level were more structured in PT than in matched O-PDX samples (Sup.Fig. 12H & I). A clear example is the expression pattern of *FAP* within tumor clusters in PT vs O-PDX samples (Sup.Fig. 12F). Additionally, analysis of the average expression of CAF-associated genes across tumor clusters revealed heterogeneous expression, with multiple clusters exhibiting varying levels of these genes (Sup.Fig. 12I). Together, these findings suggest that while core tumor programs are retained, O-PDX models reduced coordination of fusion-driven and CAF-associated gene expression compared to PT tumor cells.

Gene expression profiles also differ between PT and matched O-PDX samples. For example, the expression of *LMX1A*, a transcriptional factor involved in neuronal lineage specification (101, 102), was reduced in matched O-PDX samples compared to PT samples (Sup.Fig.12C). Additionally, *LRRC15*, an adhesion molecule associated with migration and invasion (120), was only detected in PT samples, further underscoring differences between the two conditions (Sup.Fig. 13A). To further investigate this, we analyzed a bulk RNA-seq dataset from St. Jude Cloud (45) comprising 283 samples (Sup.Table 4), including 81 EwS samples (PT = 60 & PDX = 21). Analysis of *LRRC15* expression in the EwS samples further confirmed differential *LRRC15* expression between PT and PDX models (Sup.Fig. 13B). In EwS PT-TME, LRRC15 is expressed by both classical CAFs (snRNA = 10; scRNA = 15) and EwS CAF-like snRNA = 16, 22, 25; scRNA = 8, 5) subpopulations (Sup.Fig. 13A & E). LRRC15 has attracted considerable interest as a therapeutic target in cancer, largely because of its expression on CAFs, which contribute to remodeling the TME and facilitating immune evasion (121–123), and we are actively developing LRRC15 CAR T cells for other tumor types. However, in EwS TME classical CAFs are limited and not all CAF-like subpopulations express *LRRC15*. Also, *LRRC15* expression appears to be largely confined to primary PT samples (Sup.Fig. 13C & D). Although not shown, *LRRC15* expression in EwS tumor cell lines (A673, TC32, and CHLA10) was also barely detectable, indicating a need to identify additional unique EwS TAs.

### Integrated TAs Discovery Pipeline Identifies Novel Targets in EwS TME

Identifying putative chimeric TAs in EwS requires understanding tumor and TME heterogeneity, as well as better characterization of stromal versus tumor transcriptional states. Having established a framework to characterize EwS bulk tumor cells, CAF-like subpopulations, and classical CAFs within the TME, we next sought to identify novel surface TAs for potential CAR T-cell development. Using the Human Protein Atlas (HPA), we initially obtained a list of 2,467 genes annotated as plasma membrane or subcellular proteins (48). This list was subsequently curated and refined to include only surface plasma membrane genes with some, a single, or no RNA expression across normal tissue within the HPA scRNA-seq data (Fig. 6A). These 322 genes were used as initial TAs candidates and were independently detected in both the snRNA-seq and scRNA-seq datasets. All 322 genes were then fed into our candidate TAs identification pipeline, where candidates where then prioritized using a stepwise filtering pipeline designed to identify plasma membrane-associated proteins enriched in EwS tumor and CAFs while exhibiting limited expression in all other non-tumor cell populations. The workflow incorporated sequential filtering criteria based on cellular localization, tumor enrichment, and expression across tumor and TME cell states, and is described in detail within the Methods section. This approach yielded 40 hits in the snRNA dataset and 25 in the scRNA dataset. Notably, 14 candidates were shared between the two datasets, and the top 10 candidates were prioritized for downstream analysis (Fig. 6B & C). These 10 top candidates were selected based on their tissue expression profiles in the HPA database, with the 4 excluded candidates exhibiting protein expression across multiple normal tissues. Except for ATP2B3, with a distinct neuronal expression profile, no detectable protein-level expression was observed for the identified 10 putative TAs within the database. Additionally, RNA expression of these candidates was very discrete, and plasma membrane localization was confirmed (Table 1). Collectively, these characteristics support the identified putative surface TAs as promising candidates for CAR T-cell therapy and underscore the need for further investigation.

Interestingly, *LRRC15* was identified in this screen as one of the candidate surface TAs. However, as noted previously, its expression was very low across tumor clusters. The expression of *LRRC15* also differed between snRNA and scRNA datasets, likely due to multiple factors, including the predominance of primary tumor site samples in the scRNA cohort (Fig. 6D & E). As previously mentioned, *LRRC15* expression in EwS CAF-like cells was predominantly observed in primary PT samples (Sup.Fig. 13C & D). Additionally, the scRNA dataset contained a relatively small number of cells, resulting in limited representation of classical CAFs and a mixed population of mCAFs and iCAFs (Sup.Fig. 8D). Overall, *LRRC15* is expressed in the PT EwS TME, but at lower levels than in other sarcomas, such as osteosarcoma (OS), and is much more difficult to study in PDX and *in vitro* conditions (Fig. 7F). The other nine putative TAs (*ATP2B3*, *CACNA1I*, *DCHS2*, *DSEL*, *LPAR4*, *PRRT4*, *TMEM229A*, *UNC5A*, and *UNC79*) exhibited substantially higher RNA expression than *LRRC15* across the EwS TME and tumor clusters (Fig. 6C).

While all the putative surface TAs identified in the screen were detected independently in both datasets (Fig. 6C; Sup.Fig. 14), we also assessed each TAs expression profile across grouped EwS, EwS CAF-like, CAF, and normal clusters (Fig. 6D & E). Both *ATP2B3* and *DSEL* were expressed in both bulk tumor and EwS CAF-like clusters (Fig. 6D & E). In the snRNA dataset, *ATP2B3* expression was higher in EwS tumor clusters (Fig. 6D), whereas in the scRNA dataset it was higher in EwS CAF-like clusters (Fig. 6E), further highlighting the inherent differences between the two datasets. While *DSEL*, across both snRNA-seq and scRNA-seq datasets, uniquely showed high expression in EwS CAF-like clusters while remaining expressed in EwS tumor clusters (Fig. 6D & E). *DCHS2* and *LPAR4*, in both datasets, were expressed in EwS tumor clusters (Fig. 6D & E). To evaluate whether the expression of the 10 putative TAs is maintained in both primary and metastatic EwS tumors, thereby assessing their potential as broadly applicable therapeutic targets, we analyzed RNA expression profiles using the snRNA dataset. *ATP2B3*, *CACNA1I, and PRRT4* expression were higher in metastatic tumor samples than in primary tumor clusters (Sup.Fig. 15). While *LRRC15* and *TMEM229A* expression was higher in primary tumor samples than in metastatic tumors (Sup.Fig. 15). For *DCHS2*, *DSEL*, *LPAR4*, *UNC5A*, and *UNC79* expression between primary and metastatic tumors was similar (Sup.Fig. 15). The expression of all putative TAs was also confirmed using the St. Jude bulk RNA-seq dataset (Fig. 7). Additionally, with the exception of *LRRC15*, all other putative surface TAs were detected in both PT and PDX EwS samples. *DSEL*, *LPAR4*, and *LRRC15* also had strong expression profiles in OS samples (Fig. 7D, E, & F). Moreover, differences in expression patterns between PT and PDX across putative TAs reveal that LRAR4 is more highly expressed in PDX than in PT samples in OS (Fig. 7E), further highlighting the transcriptional differences between PT and PDX samples. To further characterize the expression profiles of the 10 putative TAs, RNA expression was evaluated using the Cancer Cell Line Encyclopedia (CCLE) dataset available through the DepMap portal (DepMap Public 26Q1) (46, 47). Consistent with our previous findings, LRRC15 exhibited minimal to no RNA expression in EwS cell lines, whereas the remaining 9 putative TAs showed substantially higher transcript expression (Sup.Fig. 16). Expression levels varied among EwS cell lines and across the candidate TAs, further highlighting the heterogeneity of EwS cell lines. Furthermore, analysis of the broader DepMap cancer cell line collection revealed that several of these candidate TAs were also highly expressed in cell lines representing other cancer types, further indicating that their expression is not restricted to EwS (Sup.Fig. 16).

To determine whether the candidate TAs were also expressed at the protein level, we analyzed the harmonized CCLE mass spectrometry (MS) dataset available through the DepMap portal (46, 47). Compared with the CCLE RNA-sequencing dataset, the CCLE MS dataset contained substantially fewer profiled cell lines, including only 4 EwS cell lines (A673, SK-N-MC, SK-ES-1, and TC-71). Furthermore, only 3 of the 10 putative TAs (ATP2B3, DCHS2, and LRRC15) were represented in the MS dataset following UniProt annotation and protein mapping, limiting protein-level evaluation of the 7 remaining candidates. Consistent with the RNA expression analysis, LRRC15 protein expression was largely absent in EwS cell lines, whereas ATP2B3 and DCHS2 exhibited higher normalized protein expression levels. Protein expression levels also varied among the four EwS cell lines, further supporting the heterogeneous expression of these candidate surface TAs within EwS cell lines (Sup.Fig. 17). We therefore examined the DepMap CCLE reverse-phase protein array (RPPA) dataset, an antibody-based proteomic platform, to determine whether these candidate TAs were represented in an independent protein expression dataset. However, the RPPA dataset included substantially fewer profiled cell lines and quantified only a limited panel of 144 proteins. Consequently, none of the 10 putative TAs were represented in the RPPA dataset. Although protein expression of these putative TAs remains to be experimentally validated in EwS cell lines, PDXs, and PT samples, evidence from the HPA supports their protein expression. All 10 candidate TAs have immunofluorescence assay (IFA) data within the HPA database generated in cancer cell lines demonstrating protein expression and localization to the plasma membrane.

## Discussion

In this study, we used complementary snRNA-seq (13 EwS samples from ALSF) (43) and scRNA-seq (GEO accession GSE243347) (44) datasets to characterize the cellular heterogeneity of PT EwS tumors and to identify putative tumor-associated surface TAs for the development of targeted immunotherapeutic strategies, including CAR T-cell therapy, T-cell engagers (TCEs), and antibody-drug conjugates (ADCs), and related immunotherapeutic approaches. Our analysis demonstrates that PT EwS tumor cells are not transcriptionally uniform but instead occupy multiple cellular states, including neuronal-like, proliferative, angiogenic, and CAF-like programs. This further supports original findings in EwS cell lines and other *in vivo* and *in vitro* models (28, 34, 35). These transcriptional states were observed across independent datasets and support a model in which EwS tumor cells diversify along multiple transcriptional axes, indicating substantial intratumoral heterogeneity in patient-derived EwS samples.

With the discovery of EwS CAF-like cells (28), significant questions have arisen about the presence of classical CAFs within the TME and the transcriptional differences between classical CAFs and CAF-like cells. Here, we show that in both datasets, EwS TME harbors classical CAFs and EwS CAF-like tumor subpopulations (Fig. 1; Sup.Fig. 4). These EwS CAF-like subpopulations transcriptionally resemble stromal cells while retaining tumor-associated features, exhibiting reduced neuronal-like features alongside increased mesenchymal-like signatures, extracellular matrix programs, and CAF-associated genes. Trajectory analysis further supported a progressive transition from main tumor states, with high neuronal-like signatures, toward CAF-like and stromal-like transcriptional programs (Fig. 4; Sup.Fig. 9). This suggests that EwS cells may undergo dynamic transcriptional reprogramming within the TME, potentially allowing tumor cells to remodel the extracellular matrix, alter immune accessibility, and promote invasive or metastatic behavior in a coordinated manner. These findings are consistent with previous reports describing EwS CAF-like cells and further extend these concepts to PT samples.

Our comparison of PT and matched PDX samples also highlights the importance of tumor context. Although core EwS transcriptional programs were retained in PDX models, the coordinated CAF-like and TME-associated programs observed in PT were less apparent. At the RNA level, additional transcriptional differences were evident between PT and PDX; for example, *LRRC15* expression was completely lost in PDX models (Sup.Fig. 13). This suggests that PDX models may not fully capture the stromal and immune pressures present in patient tumors. As a result, TAs discovery based only on PDX or cell line models may overlook antigens associated with specific tumor-TME states, particularly those enriched in EwS CAF-like subpopulations.

Using EwS PT samples, we identified a set of putative surface TAs expressed in bulk EwS clusters and CAF-like clusters. Several candidates, including *ATP2B3*, *CACNA1I*, *DCHS2*, *DSEL*, *LPAR4*, *PRRT4*, *TMEM229A*, *UNC5A*, and *UNC79* (Fig. 6), showed expression in EwS tumor clusters and were confirmed in a larger St. Jude bulk RNA dataset (Fig. 7). Notably, many of these candidates are associated with neuronal biology, reflecting the broader neuronal-like transcriptional state observed in EwS tumors. This is particularly evident for *ATP2B3*, *CACNA1I*, *UNC5A*, and *UNC79*, which have known or inferred roles in neuronal signaling, calcium transport, axon guidance, and/or ion channel regulation. Notably, none of the identified candidate surface TAs have been previously characterized in EwS, underscoring the novelty of our approach and the value of PT tumor–based analysis in identifying previously unknown therapeutic targets.

Among the identified candidates, several exhibited features that support their prioritization for further validation, while also highlighting important gaps in current biological and therapeutic potential. *CACNA1I* and *ATP2B3*, both involved in calcium transport (124, 125), were consistently detected across datasets and in both PT and PDX bulk RNA samples (Fig. 7), with both demonstrating enrichment in metastatic tumor populations within the snRNA dataset (Sup.Fig. 15). Notably, *ATP2B3* represents a more tractable candidate with emerging therapeutic interest, as preclinical inhibitors have been proposed (126), whereas *CACNA1I*, although previously suggested as a potential target in other cancers (127), remains largely unexplored. In contrast, *DCHS2* and *DSEL* represent less well-characterized candidates, with limited functional and therapeutic data available. Both were expressed in EwS tumor clusters across datasets, with *DCHS2* showing expression in EwS clusters, while *DSEL* displayed a distinct pattern with high expression in EwS CAF-like clusters in addition to the tumor clusters (Fig. 6). These findings suggest that some putative surface TAs may target not only the main tumor populations but also the transcriptionally distinct EwS CAF-like states that contribute to TME remodeling. Collectively, most of these putative TAs remain poorly characterized, particularly with respect to their RNA and protein expression levels. From a therapeutic perspective, optimal targets should exhibit discrete protein expression in tumor cells with minimal or absent expression in normal tissues, thereby maximizing specificity and minimizing potential toxicity. In general, while RNA expression is not a definitive indicator for protein abundance, transcriptomic profiles can provide an initial indication of the likelihood of protein expression. In particular, low or undetectable transcript levels across normal tissues may support a reduced likelihood of substantial corresponding protein expression, although this relationship is gene- and tissue-dependent and requires direct protein-level validation. Consistent with this framework, using the HPA database showed that the putative surface TAs exhibited low or undetectable RNA expression across most normal tissues while our analyses demonstrate strong RNA expression profile in EwS tumors. Additionally, the RNA expression patterns observed with these putative surface TAs suggest that these candidates may not only target EwS tumors but may also extend to an array of other tumors.

An important consideration is that RNA-based antigen discovery does not necessarily reflect protein-level expression. To address this, available protein-level data for the identified surface TAs was evaluated through literature review and interrogation of several publicly available datasets, including the HPA database, but supporting protein-level evidence remains limited. Using the HPA database, almost all putative surface TAs identified lacked detectable protein expression within normal tissue, despite evidence of plasma membrane localization based on IFA staining in a cancer cell line. This emphasizes the need for future studies to validate these candidate TAs experimentally at the protein level using antibody-based methods in EwS and other tumor types. In addition, several candidates remain poorly characterized in cancer biology, particularly *PRRT4*, *TMEM229A*, and *UNC79*. While this increases their novelty, it also underscores the need for functional studies to determine whether these targets are consistently expressed at the cell surface and whether targeting them would be safe and effective.

Overall, our findings support a model in which EwS tumors harbor transcriptionally diverse tumor cell states, including EwS CAF-like subpopulations. The EwS TME contains both classical CAFs and CAF-like tumor populations, both of which may contribute to matrix remodeling, immune evasion, and therapeutic resistance. By integrating single-nucleus/cell tumor-state analyses with surface TAs prioritization, this study identifies novel candidate surface TAs that may improve immunotherapy targeting strategies in EwS. Future work should focus on validating protein expression in-house, assessing normal tissue safety, and testing whether these candidates can be targeted individually or in combination to overcome antigen heterogeneity and improve CAR T-cell efficacy in EwS.

## List of Abbreviations

CAF: Cancer-Associated Fibroblast
CAR: Chimeric Antigen Receptor
CCLE: Cancer Cell Line Encyclopedia
CNV: Copy Number Variation
DEG: Differentially Expressed Gene
ECM: Extracellular Matrix
EMT: Epithelial-to-Mesenchymal Transition
EwS: Ewing Sarcoma
GO: Gene Ontology
HPA: Human Protein Atlas
iCAF: Inflammatory Cancer-Associated Fibroblast
mCAF: Matrix Cancer-Associated Fibroblast
MSC: Mesenchymal Stem Cell
NK: Natural Killer
O-PDX: Orthotopic Patient-Derived Xenograft
OS: Osteosarcoma
PCA: Principal Component Analysis
PC: Principal Component
PCGP: Pediatric Cancer Genome Project
PDX: Patient-Derived Xenograft
PT: Primary Tissue
QC: Quality Control
RCS: Round Cell Sarcoma
RNA-seq: RNA Sequencing
RPPA: Reverse-Phase Protein Array
scRNA-seq: Single-Cell RNA Sequencing
ScPCA: Single-Cell Pediatric Cancer Atlas
SCEVAN: Single-Cell Variational Aneuploidy Analysis
SNN: Shared Nearest Neighbor
snRNA-seq: Single-Nucleus RNA Sequencing
SS: Synovial Sarcoma
TAs: Tumor-Associated Antigens
TAM: Tumor-Associated Macrophage
TGF-β: Transforming Growth Factor Beta
TME: Tumor Microenvironment
t-SNE: t-Distributed Stochastic Neighbor Embedding
UMAP: Uniform Manifold Approximation and Projection
UMI: Unique Molecular Identifier
vSMC: Vascular Smooth Muscle Cell

**Supplemental Table 1.** Metadata from patient samples obtained through the Alex’s Lemonade Stand Foundation (ALSF) scPCA Portal. The dataset includes 13 EwS samples comprising 7 tissue samples from 5 EwS patients and 6 matched O-PDX samples. These samples came with unfiltered snRNA-sequenced raw counts which were used as the starting point for the analysis.

File Name: **ST1_ALSF_Metadata.csv**

**Supplemental Table 2.** Metadata from patient samples obtained through GSE243347 using the NCBI GEO portal. The dataset includes 9 EwS primary tissue samples from 6 patients that were directly sequenced using the CEL-Seq2 method. These samples came with unfiltered scRNA-sequenced raw counts which were used as the starting point for the analysis.

File Name: **ST2_ GSE243347_Metadata.csv**

**Supplemental Table 3.** Metadata for 11 gene sets, including the total number of genes in each gene set and the percentage of overlapping genes between gene sets.

File Name: **ST3_ Gene_Sets.csv**

**Supplemental Table 4.** Metadata of 283 Bulk RNA-seq samples obtained through St. Jude Cloud. The dataset includes 5 CCS (Clear Cell Sarcoma), 81 EwS (Ewing Sarcoma), 173 OS (Osteosarcoma), 6 RCS (Round Cell Sarcoma) and 18 SS samples (Synovial Sarcoma).

File Name: **ST4_Bulk_RNA_Metadata.csv**

**Supplementary Figure 1.**
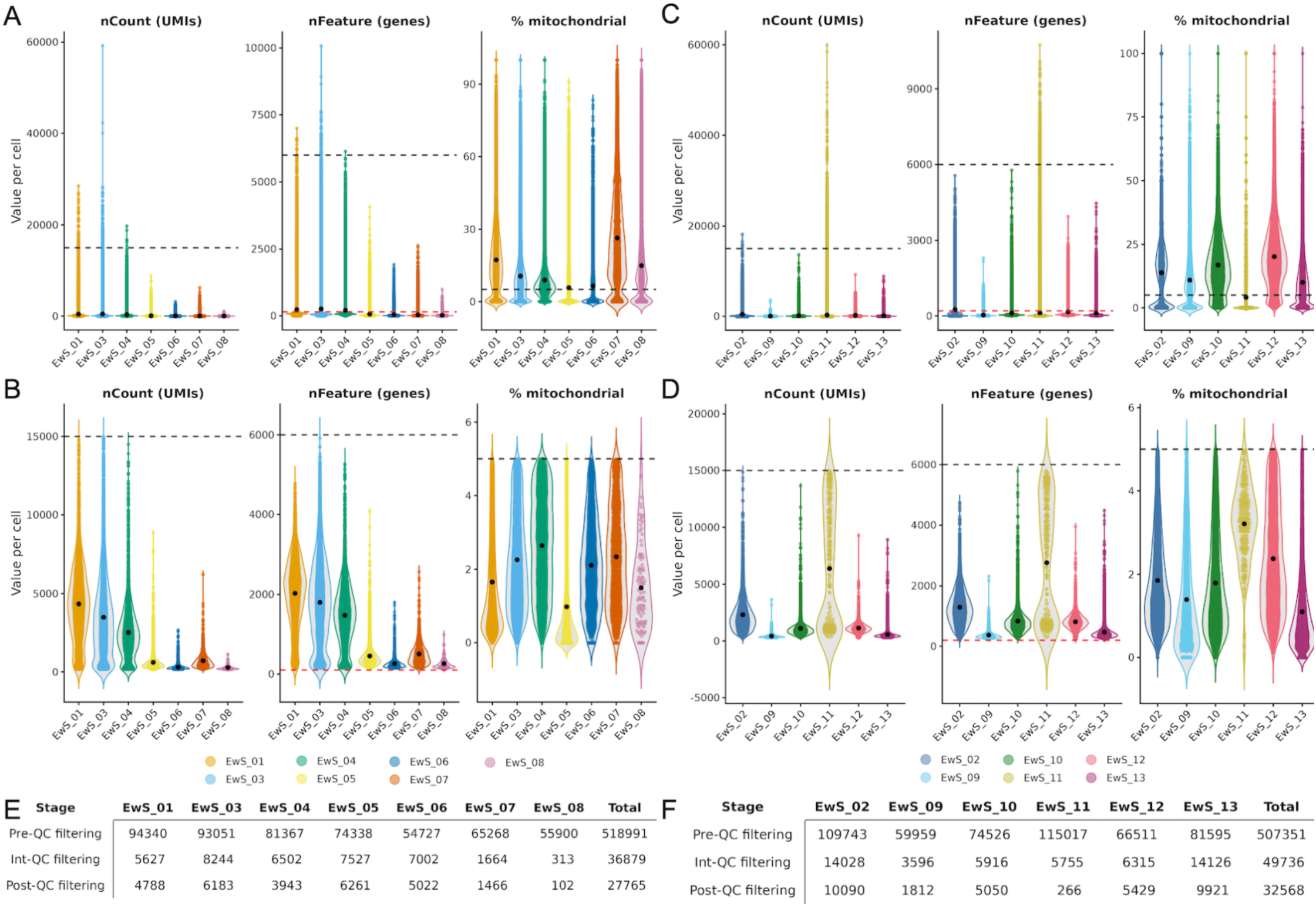
Quality Control of ALSF snRNA-Seq dataset. The 13 ALSF EwS snRNAseq samples’ raw counts were initially separated into groups: Primary Tumor (PT; A & B) and O-PDX (PDX; C & D) samples (color coded). (A – D) Plots correspond to per-nucleus quality metrics: nCount (UMIs), nFeature (genes), and % mitochondrial. Violin plots depict distribution within samples, while overlaid jittered points represent individual nuclei. Black circles indicate the mean value per sample. Horizontal dashed lines denote QC thresholds: nCount < 15,000 (upper, black); nFeature < 6,000 (upper, black) and > 200 (PT:150 and PDX: 200) (lower, red); percent.mt < 5% (upper, black). nCounts also was filtered with > 150 for PT and > 300 for PDX samples. The same QC pipeline was used for all ALSF snRNA-Seq datasets and thresholds are indicated on both (A & C) Pre-QC filtering plots and (B & D) Post-QC filtering plots. QC was performed in two steps: initial filtering using hard cutoffs, followed by refined filtering accounting for ambient RNA, doublets, and nuclear contamination. Nuclei counts per sample and total counts for (E) PT and (F) PDX are indicated in the tables.

**Supplementary Figure 2.**
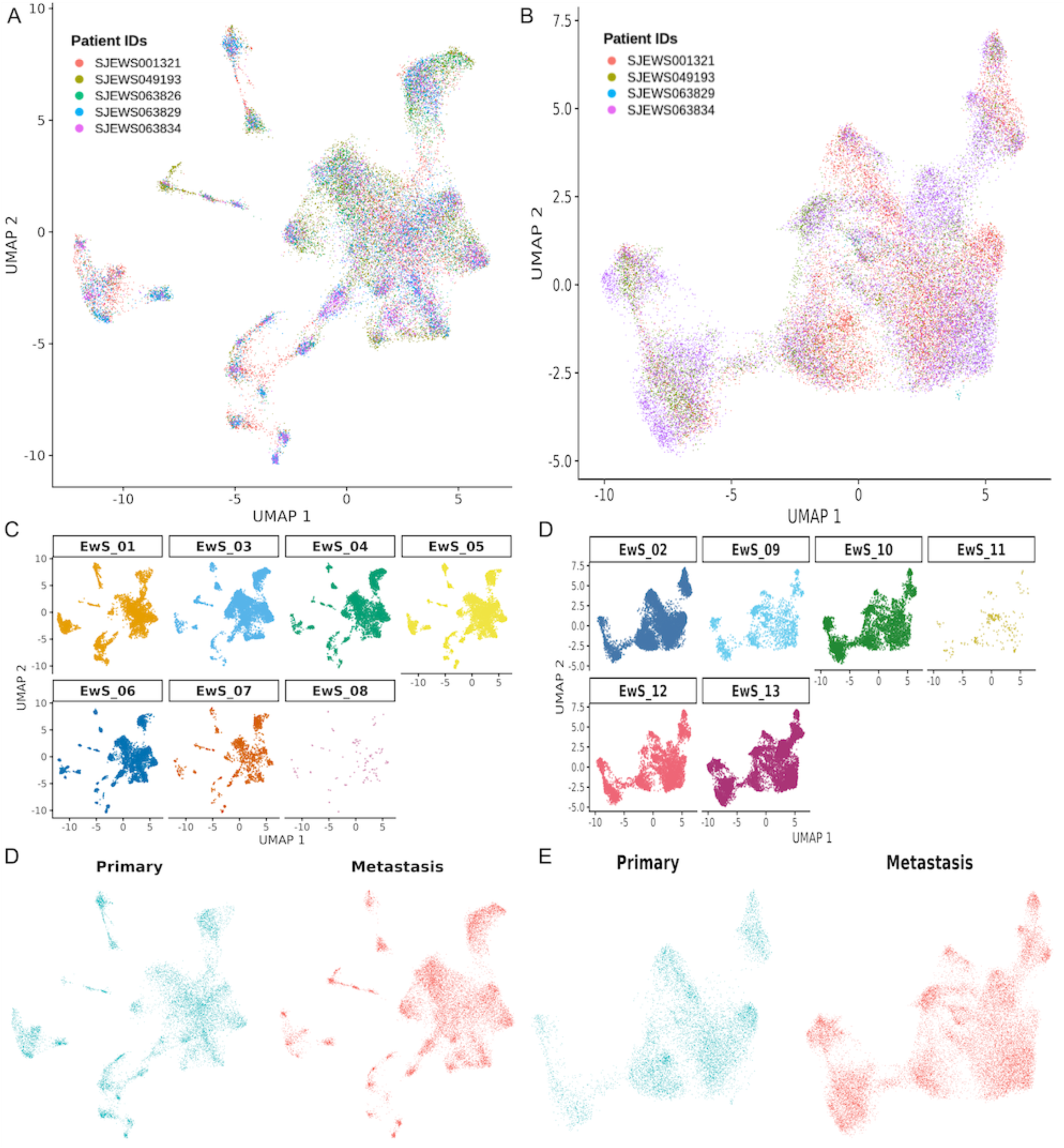
Harmony-integrated snRNA landscape. After SCTransform normalization and PCA, we used Harmony to integrate all samples into a shared low-dimensional space that emphasizes biological similarity across patients, enabling joint clustering and visualization. From this integrated space, we built a shared-nearest-neighbor graph using the first 20 PCAs, identified communities with the default Louvain algorithm (resolution = 0.8), and visualized the data with UMAP computed from the same PCs. UMAP of all nuclei colored by Patient IDs in (A) PT (5 patients) and (B) PDX (4 patients). (C) PT and PDX (D) UMAPs split by sample ID, with each panel displaying only that sample’s nuclei, clarifying each sample’s contribution to the global UMAP structure. (E) PT and PDX (F) UMAPs split primary (left; blue) and metastasis (right; red) samples.

**Supplementary Figure 3.**
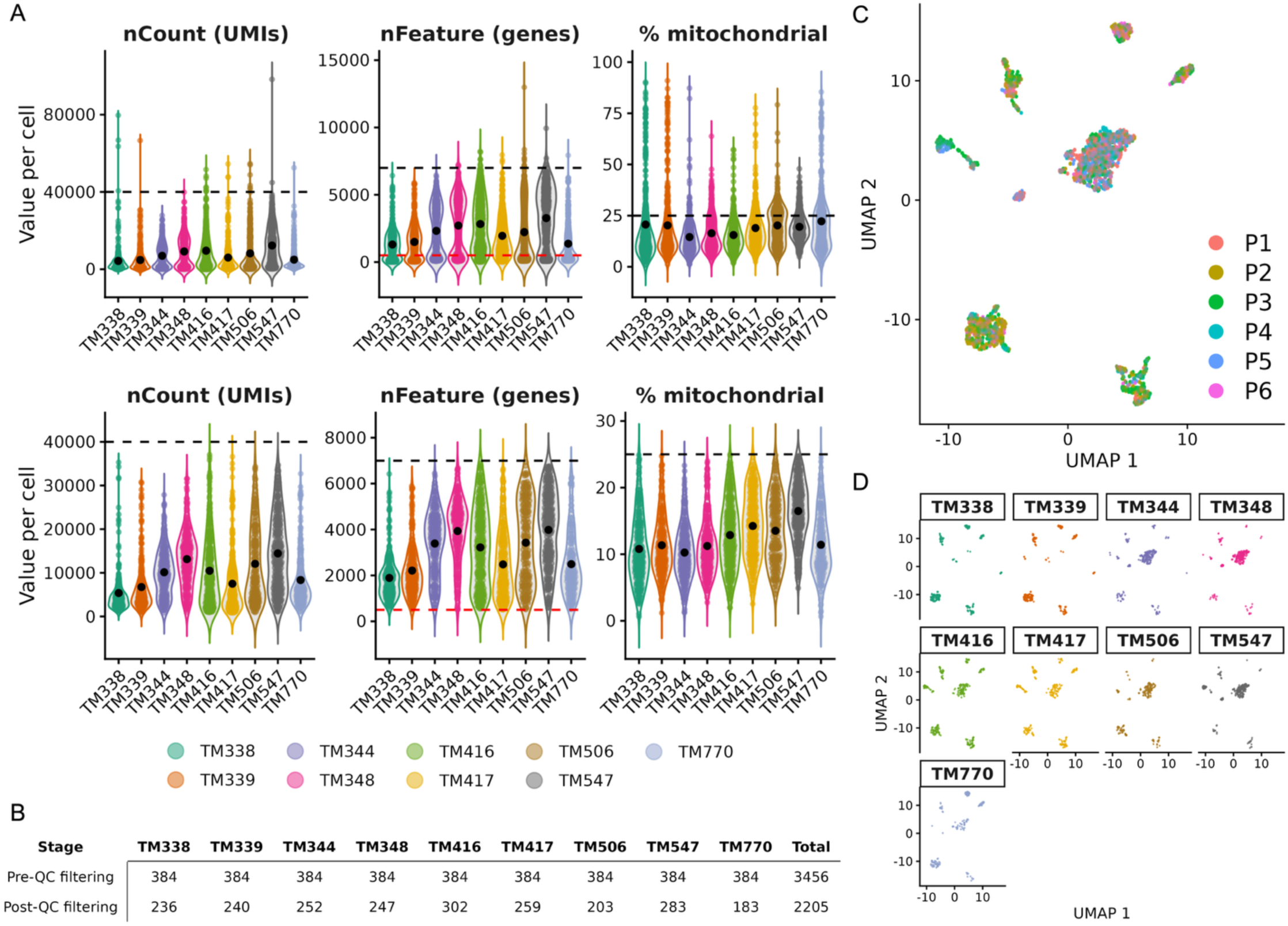
Quality Control and Harmony-integrated GSE243347 scRNA-Seq dataset. The 9 EwS scRNAseq samples’ raw counts (from 6 patient samples) were initially cleaned up and processed as Seurat objects. (A) Plots correspond to per-cell quality metrics: nCount (UMIs), nFeature (genes), and % mitochondrial. Violin plots depict distribution within samples, while overlaid jittered points represent individual cells. Black circles indicate the mean value per sample. Horizontal dashed lines denote QC thresholds: nCount = 40,000 (upper, black); nFeature = 7,000 (upper, black) and 500 (lower, red); percent.mt = 25% (upper, black). The same QC thresholds were used for all scRNA-Seq datasets, and these thresholds are indicated on both (upper) pre-QC filtering plots and (lower) post-QC filtering plots. (B) Cell counts per sample and total counts are indicated in the tables. After SCTransform normalization and PCA, we used Harmony to integrate all samples into a shared low-dimensional space that emphasizes biological similarity across patients, enabling joint clustering and visualization. From this integrated space, we built a shared-nearest-neighbor graph using the first 20 PCAs, identified communities with the default Louvain algorithm (resolution = 1.5), and visualized the data with UMAP computed from the same PCs. (C) UMAP of all cells colored by Patient IDs and (D) UMAPs split by sample ID, with each panel displaying only that sample’s cells, clarifying each sample’s contribution to the global UMAP structure.

**Supplementary Figure 4.**
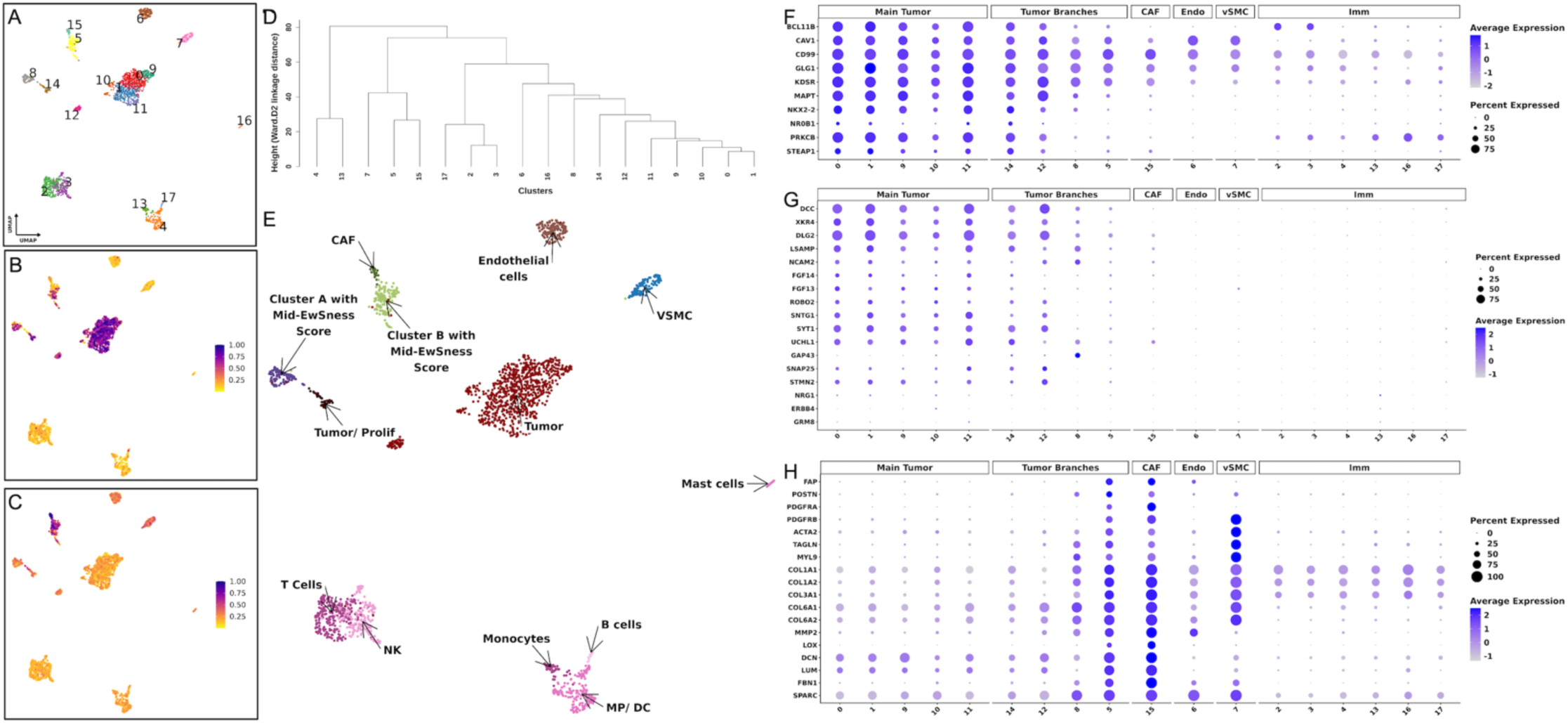
scRNA-seq transcriptomic profiling of EwS PT samples indicates a complex distribution of clusters and cell states. (A) UMAP of GSE243347 scRNA-Seq PT dataset showing unsupervised clustering of cells. 18 clusters were identified, and each point represents a single cell, colored by its cluster identity. (B) UMAP of normalized AUCell gene set (gene set 1; 32 genes upregulated by the fusion) enrichment scores. (C) UMAP of normalized AUCell gene set (gene set 3; 195 genes downregulated by the fusion) enrichment scores. High-scoring cells (purple) correspond to higher gene set expression, while low-scoring cells (yellow) correspond to lower gene set expression. (D) Hierarchical clustering of defined clusters based on PCA centroid distances. Dendrogram depicting relationships among 18 derived clusters (0–17). Cluster centroids were calculated from the first 20 principal components, and Euclidean distances were used for hierarchical clustering with Ward’s method (Ward.D2). Branch height reflects inter-cluster dissimilarity, with shorter branches indicating greater transcriptional similarity. (E) Annotated UMAP of GSE243347 snRNA-seq clusters highlighting major cell types, main tumor, and clusters that form the tumor branches. This annotation of the UMAP factors in various cell typing methods including using a reference database, CNV analysis, gene set analysis, hierarchical clustering, differential gene analysis (Final_All_Markers & Find_Marker), and top100 gene expression per cluster. Clusters that are considered as tumor branches are labeled with EwSness Scores. (F - H) Dot plot showing expression of selected (F) EwS, (G) neuronal, (H) CAF markers across clusters in different categories (Main Tumor, Tumor Branches, CAF, Endo, Epi, Peri/vSMC, Imm) of the UMAP. In each dot plot, the y-axis shows the list of genes and the x-axis shows the cluster numbers from the UMAP projections. Dot size represents the percentage of cells expressing each gene, and color intensity indicates the average expression levels.

**Supplementary Figure 5.**
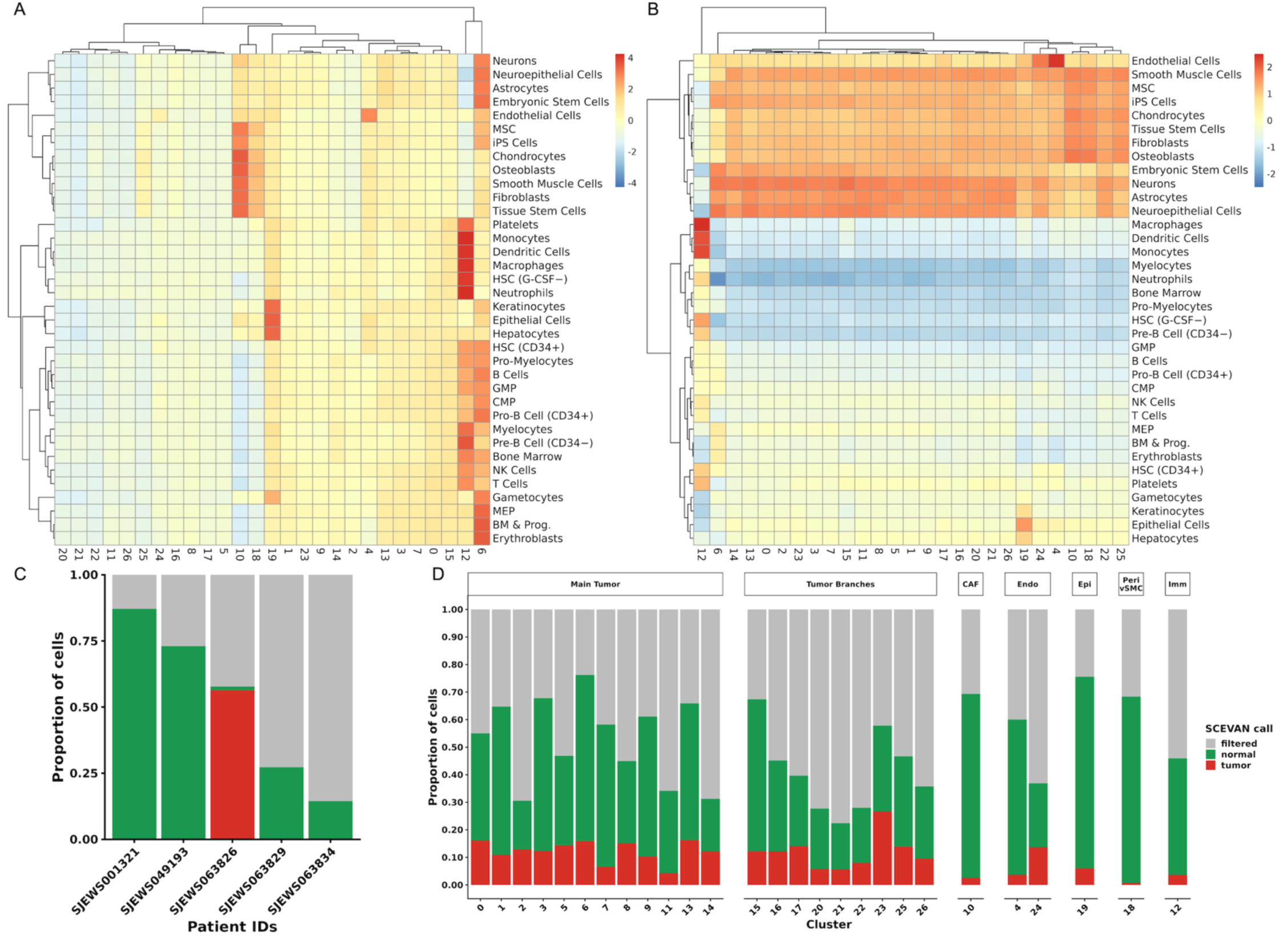
Cell Typing using reference database and SCEVAN analysis in EwS snRNA-seq dataset. (A - B) Heatmap of average SingleR annotation scores with reference cell types (rows) and cluster numbers (columns). Scores were averaged per cluster, and the heatmap was generated after (A) row scaling (z-scores across clusters within each reference cell type) to emphasize cluster-to-cluster variation for each reference cell type, or (B) column scaling (z-scores across reference cell types within each cluster) to highlight the most enriched reference cell identity per cluster. In both cases, hierarchical clustering was applied to rows and columns using Euclidean distance and complete linkage. (C - D) Stacked bar graph showing the proportion of nuclei per (C) patient or (D) cluster classified by SCEVAN as tumor (red), normal (green), or filtered (gray) based on CNV analysis. Clusters are grouped by the different categories (Main Tumor, Tumor Branches, CAF, Endo, Epi, Peri/vSMC, Imm) from UMAP embedding.

**Supplementary Figure 6.**
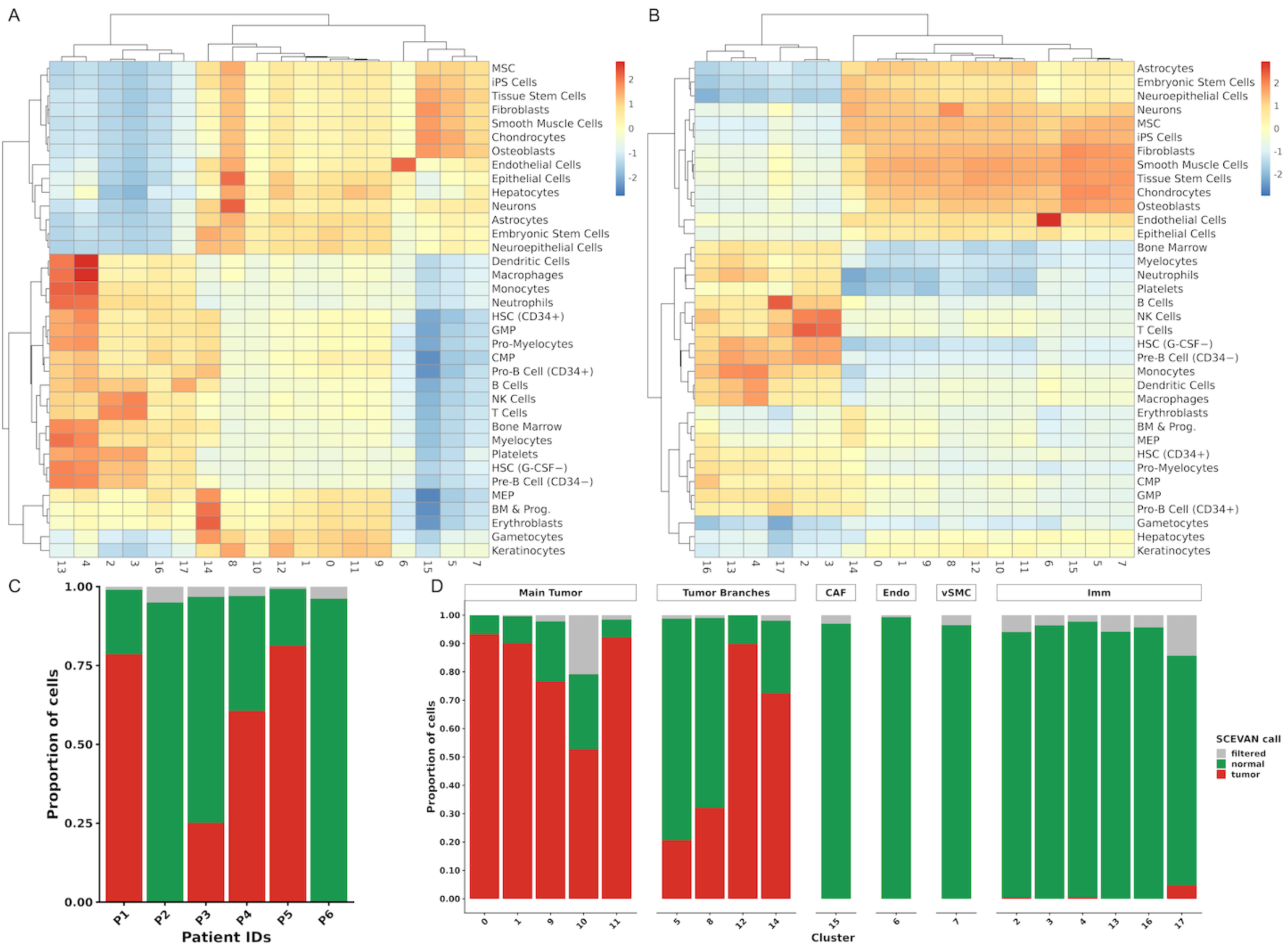
Cell Typing using reference database and SCEVAN analysis in EwS scRNA-seq dataset. (A - B) Heatmap of average SingleR annotation scores with reference cell types (rows) and cluster numbers (columns). Scores were averaged per cluster, and the heatmap was generated after (A) row scaling (z-scores across clusters within each reference cell type) to emphasize cluster-to-cluster variation for each reference cell type, or (B) column scaling (z-scores across reference cell types within each cluster) to highlight the most enriched reference cell identity per cluster. In both cases, hierarchical clustering was applied to rows and columns using Euclidean distance and complete linkage. (C - D) Stacked bar graph showing the proportion of cells per (C) patient or (D) cluster classified by SCEVAN as tumor (red), normal (green), or filtered (gray) based on CNV analysis. Clusters are grouped by the different categories (Main Tumor, Tumor Branches, CAF, Endo, vSMC, and Imm) from UMAP embedding.

**Supplementary Figure 7.**
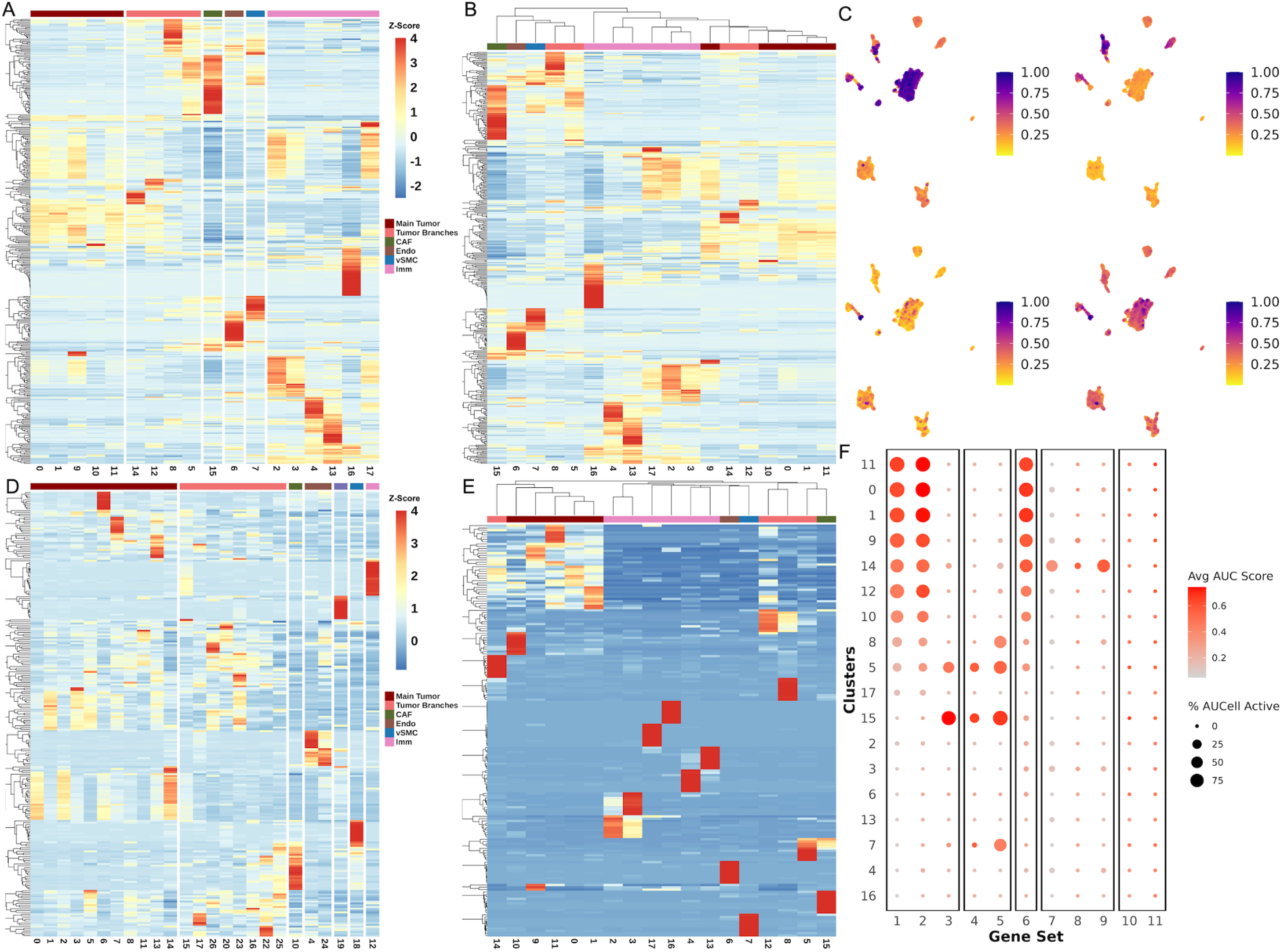
scRNA-seq cluster-specific gene expression and AUCell gene-set activity elucidates transcriptional heterogeneity in the EwS TME. (A & B) Heatmap showing the top 100 highest-expressed genes per defined cluster based on average normalized RNA expression sorted by (A) defined categories or (B) unbiased hierarchical clustering using top 100 genes. (C) UMAP of normalized AUCell gene set enrichment scores. Upper-left (gene set 2; 439 genes upregulated by the fusion), upper-right (gene set 5; 200 hallmark epithelial mesenchymal transition genes), lower-left (gene set 7; 37 gene set activated by ATR response to replication stress), lower-right (gene set 8; 184 gene set involved in DNA replication). High-scoring cells (purple) correspond to higher gene set expression, while low-scoring cells (yellow) correspond to lower gene set expression. (D & E) Heatmap showing the top 10 differentially expressed markers per cluster, ranked by average log2 fold change sorted by (D) defined categories or (E) unbiased hierarchical clustering using top 10 genes. Expression values are scaled per gene (row-wise Z-score) across clusters. Both genes and clusters are hierarchically clustered and ordered by similarity in their scaled expression profiles, grouping genes and clusters with shared patterns across clusters. Distinct blocks of high expression identify cluster-specific transcriptional signatures and reinforce higher-order relationships among related clusters. (F) Dot-plot summarizing normalized average AUCell scores (red) for each gene set (columns) per cluster (rows). Dot size represents the percentage of cells classified as “Active” for each gene set within each cluster (cells with gene-set enrichment based on AUCell AUC >= default threshold). Gene sets had minimal overlap, and Sup.Table 3 lists the 11 gene sets and the number of genes in each.

**Supplementary Figure 8.**
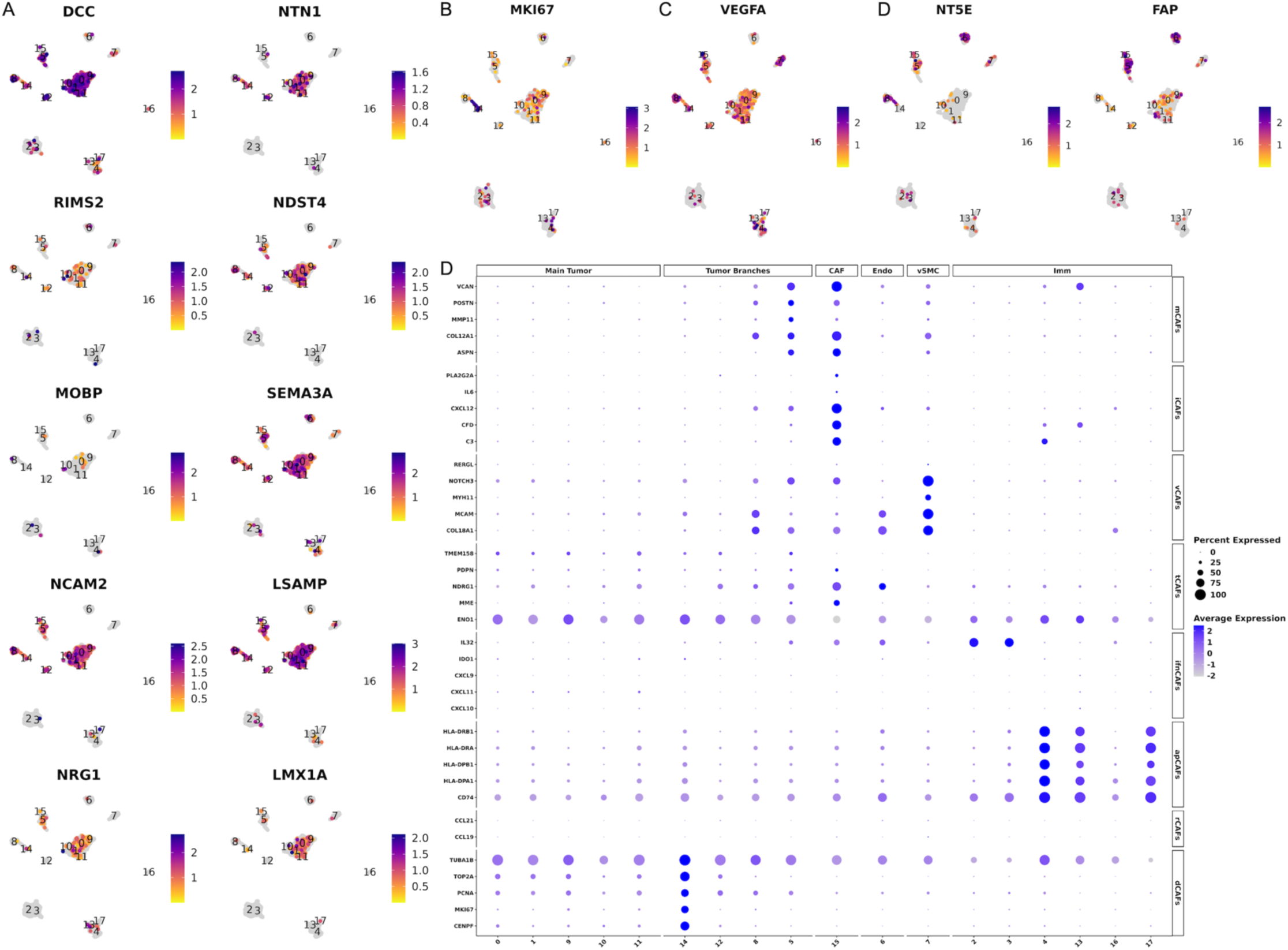
scRNA-seq gene expression profiles highlight dynamically neuronal-like, proliferative, angiogenic, and CAF-like cellular programs. (A - D) UMAP feature plots showing the distribution of selected gene expression per cell across clusters. High-scoring cells (purple) correspond to higher gene expression, while low-scoring cells (yellow) correspond to lower gene expression. (A) List of genes (*DCC*, *NTN1*, *RIMS2*, *NDST4*, *MOBP*, *SEMA3A*, *NCAM2*, *LSAMP*, *NRG1*, and *LMX1A*) expressed by the EwS tumor clusters, supporting a pervasive yet non-uniform neuronal-like transcriptional state. (B) Expression of *MKI67*, a marker for cell proliferation. (C) Expression of *VEGFA*, a marker of angiogenesis. (D) Expression of *NT5E* (CD73), a marker of EwS CAF-like cells, and *FAP*, a canonical marker of classical CAFs. (E) Dot plot showing expression of selected genes for the classification of CAF subpopulations (mCAFs = matrix; iCAFs = inflammatory; vCAFs = vascular; tCAFs = tumor-like; ifnCAFs = interferon-response; apCAFs = antigen-presenting; rCAFs = reticular-like; dCAFs = dividing) across clusters in different categories (Main Tumor, Tumor Branches, CAF, Endo, vSMC, and Imm) of the UMAP. In the dot plot, the y-axis right-hand side shows the list of genes, and the left-hand side shows the CAF subpopulation categories. The x-axis shows the cluster numbers from the UMAP projections. Dot size represents the percentage of cells expressing each gene, and color intensity indicates the average expression levels.

**Supplementary Figure 9.**
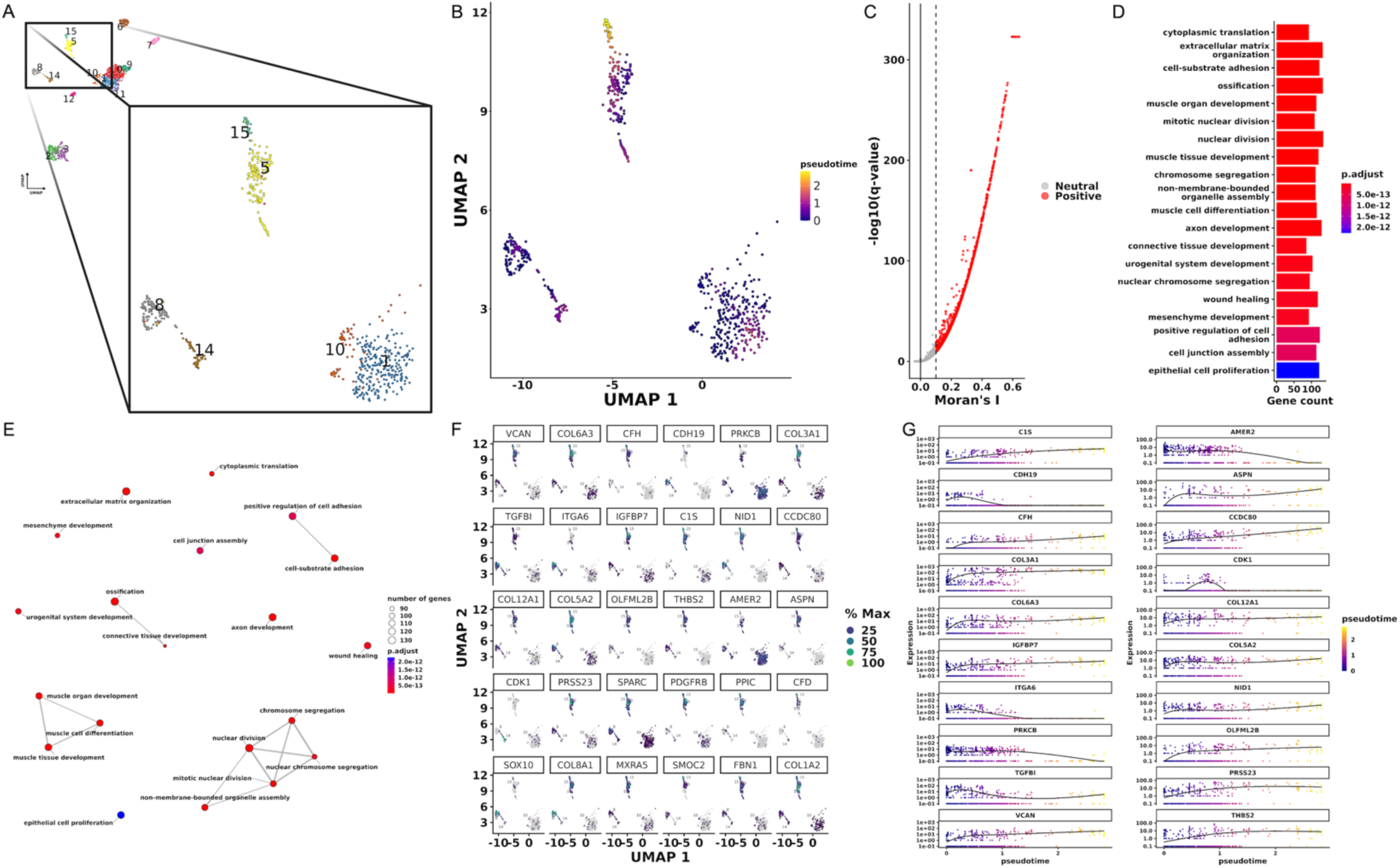
scRNA-seq Trajectory Analysis Reveals ECM Remodeling Program Along EwS–CAF Continuum. (A) UMAP illustration of the selected clusters for downstream trajectory analysis. (B) Monocle 3 pseudotime trajectory projected onto the UMAP embedding. Cells are colored by pseudotime, with early cells (purple) concentrated in the lower-right region and progressively increasing pseudotime values extending toward the central and upper-left branches (yellow). Trajectory inference was performed using reversed graph embedding in UMAP space, and cells were ordered using cluster 10 as the root. The continuous color gradient illustrates the inferred progression of cellular states along the branching trajectory structure in low-dimensional space. (C) Volcano-style scatter plot summarizing identified trajectory-associated genes on a learned principal graph after library-size normalization. Genes that showed significant variation along the trajectory are plotted by Moran’s I (x-axis), which measures how strongly expression follows the trajectory, and by statistical significance (−log10(q-value). Q-values were FDR-corrected, and zeros were replaced with a very small value to prevent infinite values during log transformation. The solid vertical line marks Moran’s I = 0, and the dashed vertical line marks Moran’s I = 0.1 cutoff, separating trajectory-structured genes (Positive; red) from neutral/weakly autocorrelated genes (Neutral; gray). (D) GO Biological Process enrichment analysis of trajectory-associated genes identified by Monocle 3 (Moran’s I > 0.1). Redundant GO terms were reduced using semantic similarity filtering (cutoff = 0.7). The top 20 enriched processes are shown on the y-axis, while the x-axis indicates the number of genes mapped to each term (gene count). The color scale represents the adjusted p-values (FDR). (E) Enrichment map network of the top 20 GO Biological Process terms derived from trajectory-associated genes (Moran’s I > 0.1). Pairwise semantic similarity between GO terms was calculated and terms were visualized as a network using a force-directed layout. Each node represents an enriched GO term; node size reflects the number of genes associated with the term, and node color indicates adjusted p-value (FDR). Edges connect terms with high semantic similarity, highlighting functional relationships and grouping related biological processes into clusters. (F) Feature plots showing expression of the top trajectory-associated genes (ranked by Moran’s I values) projected onto the UMAP embedding. Each panel displays one gene, with cells colored by % Max (scaled expression). On the Max % scale label, dark purple indicates low expression, while light green indicates high expression. (G) Gene expression dynamics across pseudotime for the top 20 trajectory-associated genes. Each panel shows normalized gene expression plotted against pseudotime, with cells colored by pseudotime (purple = early, yellow = late). The black line represents a smoothed trend curve summarizing the overall expression pattern across pseudotime. This visualization highlights genes that increase, decrease, or peak at specific stages along the inferred trajectory.

**Supplementary Figure 10.**
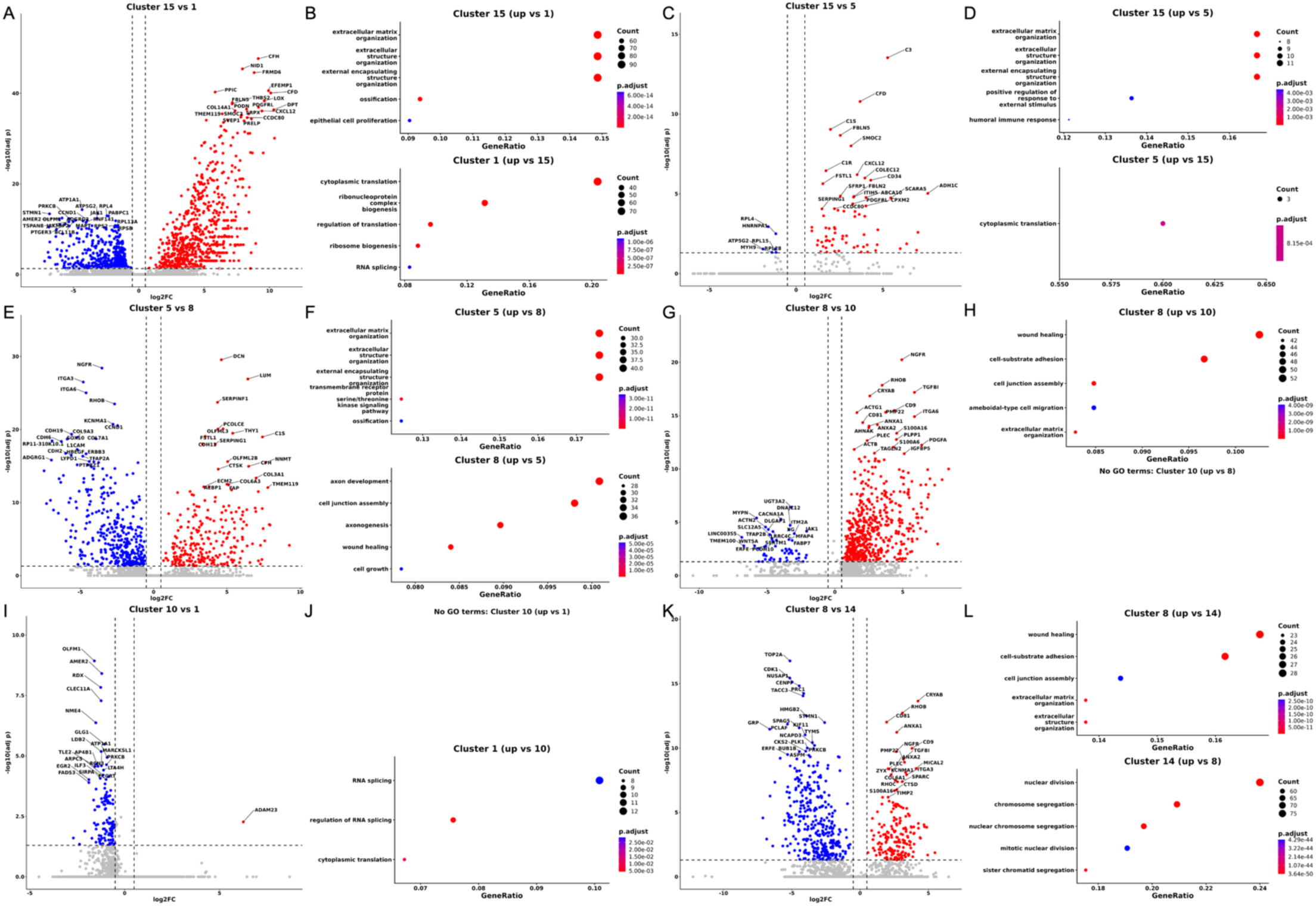
scRNA-seq Pairwise differential expression and GO enrichment analysis reveal functional differences between CAF-related clusters. Comparing transcriptional profiles of (A & B) 15 vs 1, (C & D) 15 vs 5, (E & F) 5 vs 8, (G & H) 8 vs 10, (I & J) 10 vs 1, (K & L) 8 vs 14 clusters. (A, C, E, G, I, and K) Volcano plot showing differential gene expression between selected 1-vs-1 clusters. Genes are plotted by log2 fold change (x-axis) and −log10 adjusted p-value (y-axis). Significantly upregulated genes in the initial cluster are shown in red, downregulated genes in blue, and non-significant genes in gray. Dashed vertical lines indicate log2FC cutoffs (±0.5), and the dashed horizontal line marks the adjusted p-value cutoff (FDR < 0.05). (B, D, F, H, J, and L) GO Biological Process enrichment analysis for genes upregulated in the initial (top) and second (bottom) clusters. The x-axis shows GeneRatio, dot size represents gene count, and color indicates adjusted p-value (FDR). Across all comparisons, differential expression analysis was performed using the Wilcoxon rank-sum test in Seurat, followed by GO Biological Process enrichment using clusterProfiler with Benjamini–Hochberg FDR correction. GO terms were ranked by GeneRatio, and the top enriched categories are shown for each comparison.

**Supplementary Figure 11.**
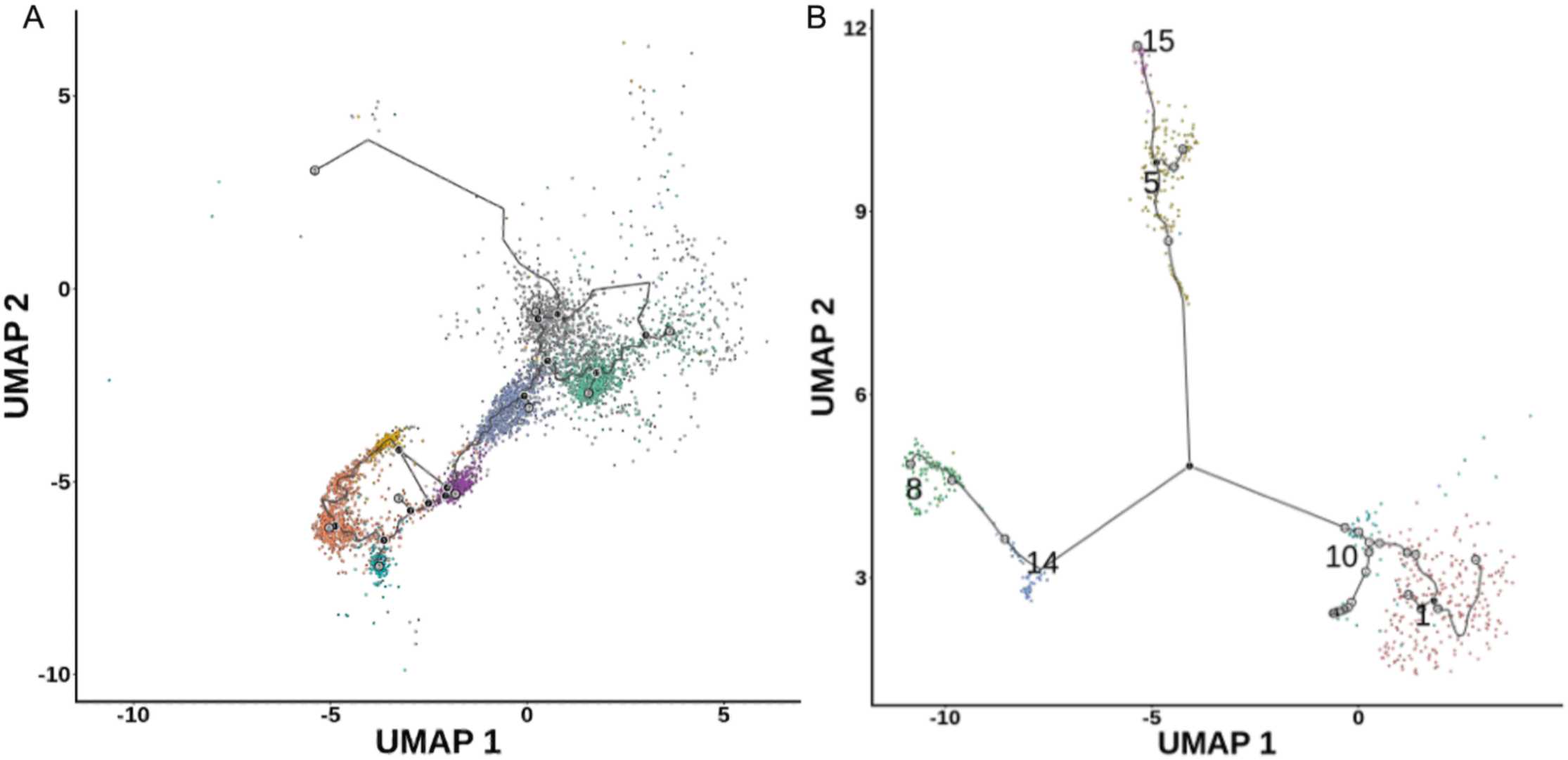
Trajectory analysis of EwS tumor nuclei and cell states using Monocle 3. (A) Monocle 3 analysis of the snRNA-seq dataset visualized in UMAP space. nuclei are colored by cluster identity, and the learned principal graph (black lines) represents inferred trajectories of transcriptional state transitions across EwS tumor nuclei. The trajectory reveals a continuum of nuclei states with multiple branching structures, consistent with transcriptional diversification. (B) Monocle 3 analysis of the scRNA-seq dataset, with cells colored by cluster identity and the learned principal graph overlaid in black. Distinct branch points and terminal states are observed, supporting the presence of multiple transcriptional trajectories within the tumor population.

**Supplementary Figure 12.**
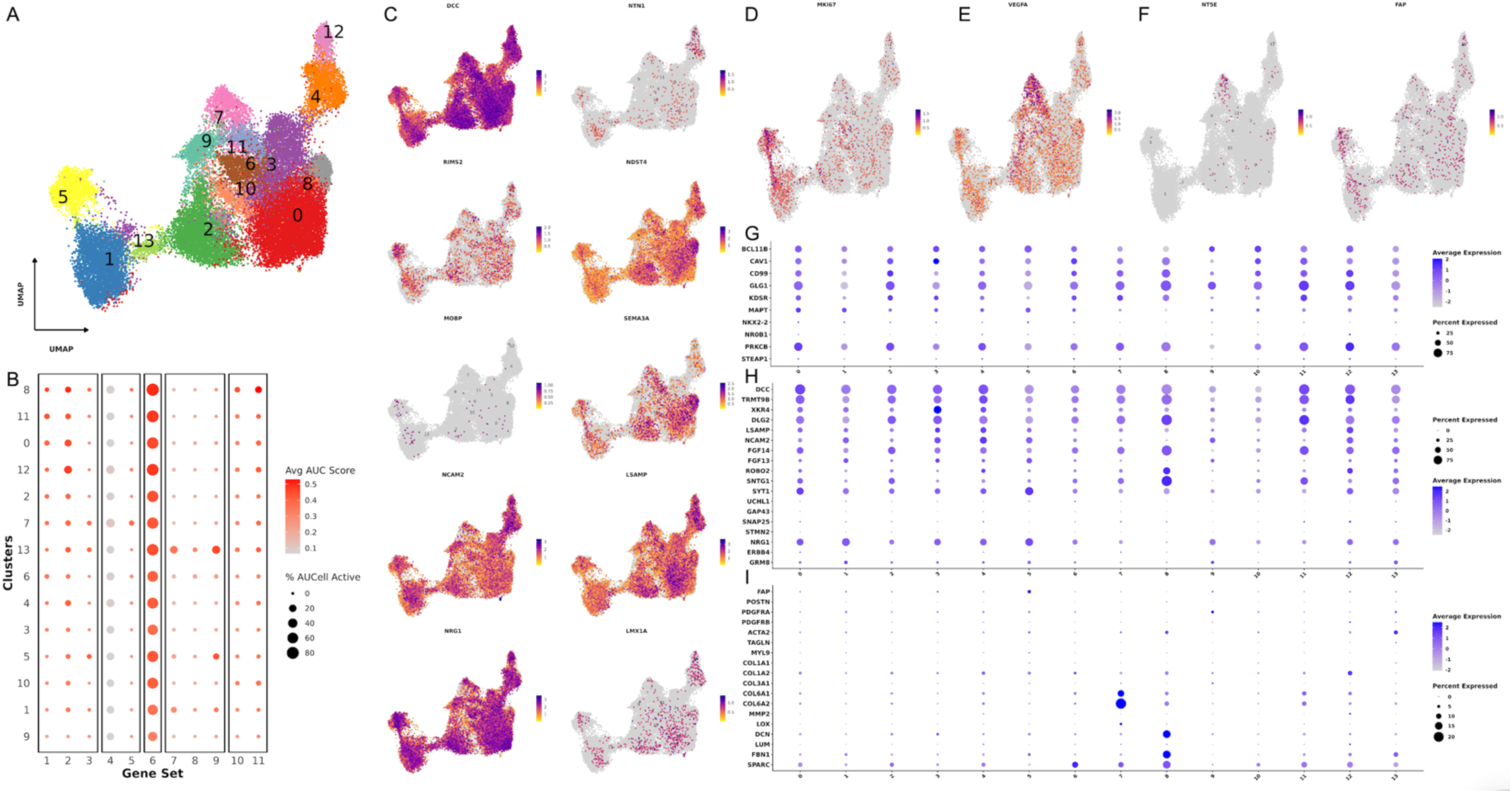
snRNA-seq gene expression profiles of matched O-PDX samples. (A) UMAP of snRNA-Seq O-PDX dataset showing unsupervised clustering of nuclei. 14 clusters were identified, and each point represents a single nucleus, colored by its cluster identity. (B) Dot-plot summarizing normalized average AUCell scores (red) for each gene set (columns) per cluster (rows). Dot size represents the percentage of nuclei classified as “Active” for each gene set within each cluster (nuclei with gene-set enrichment based on AUCell AUC >= default threshold). Gene sets had minimal overlap, and Sup.Table 3 lists the 11 gene sets and the number of genes in each. (C - F) UMAP feature plots showing the distribution of selected gene expression per nucleus across clusters. High-scoring nuclei (purple) correspond to higher gene expression, while low-scoring nuclei (yellow) correspond to lower gene expression. (C) List of genes (*DCC*, *NTN1*, *RIMS2*, *NDST4*, *MOBP*, *SEMA3A*, *NCAM2*, *LSAMP*, *NRG1*, and *LMX1A*) expressed by the EwS tumor clusters, supporting a pervasive yet non-uniform neuronal-like transcriptional state. (D) Expression of *MKI67*, a marker for cell proliferation. (E) Expression of *VEGFA*, a marker of angiogenesis. (F) Expression of *NT5E* (CD73), a marker of EwS CAF-like cells, and *FAP*, a canonical marker of classical CAFs. (G - I) Dot plot showing expression of selected (G) EwS, (H) neuronal, (I) CAF markers across clusters within the UMAP. In each dot plot, the y-axis shows the list of genes, and the x-axis shows the cluster numbers from the UMAP projections. Dot size represents the percentage of nuclei expressing each gene, and color intensity indicates the average expression levels.

**Supplementary Figure 13.**
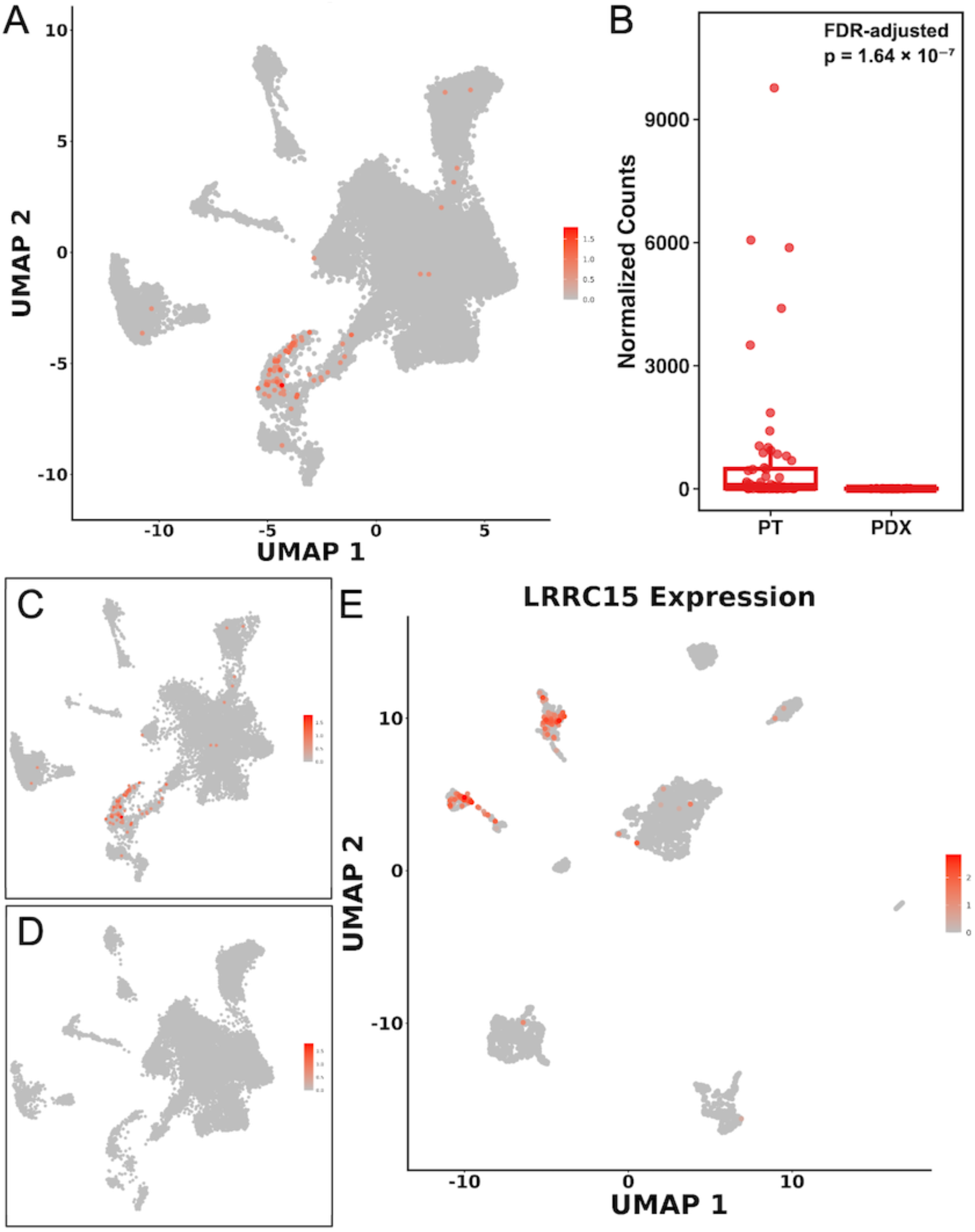
LRRC15 expression across EwS patient tumors and PDX models. (A) UMAP feature plot of snRNA-seq data from EwS PT samples, highlighting *LRRC15* expression-positive nuclei (red) and negative (gray). (B) boxplot comparison of *LRRC15* RNA expression levels in St. Jude bulk RNA-seq data of 81 EwS (PT = 60 & PDX = 21) samples. *LRRC15* expression is significantly reduced in PDX models compared to PT samples (FDR-adjusted p = 1.64 × 10⁻⁷), indicating loss of *LRRC15* expression in PDX samples. (C–D) UMAP feature plots of *LRRC15* expression in the snRNA dataset, split into (C) primary and (D) metastatic samples, demonstrate restricted *LRRC15* expression across different EwS TME conditions. (E) UMAP feature plots showing the expression patterns of LRRC15 within the scRNA dataset (GSE243347).

**Supplementary Figure 14.**
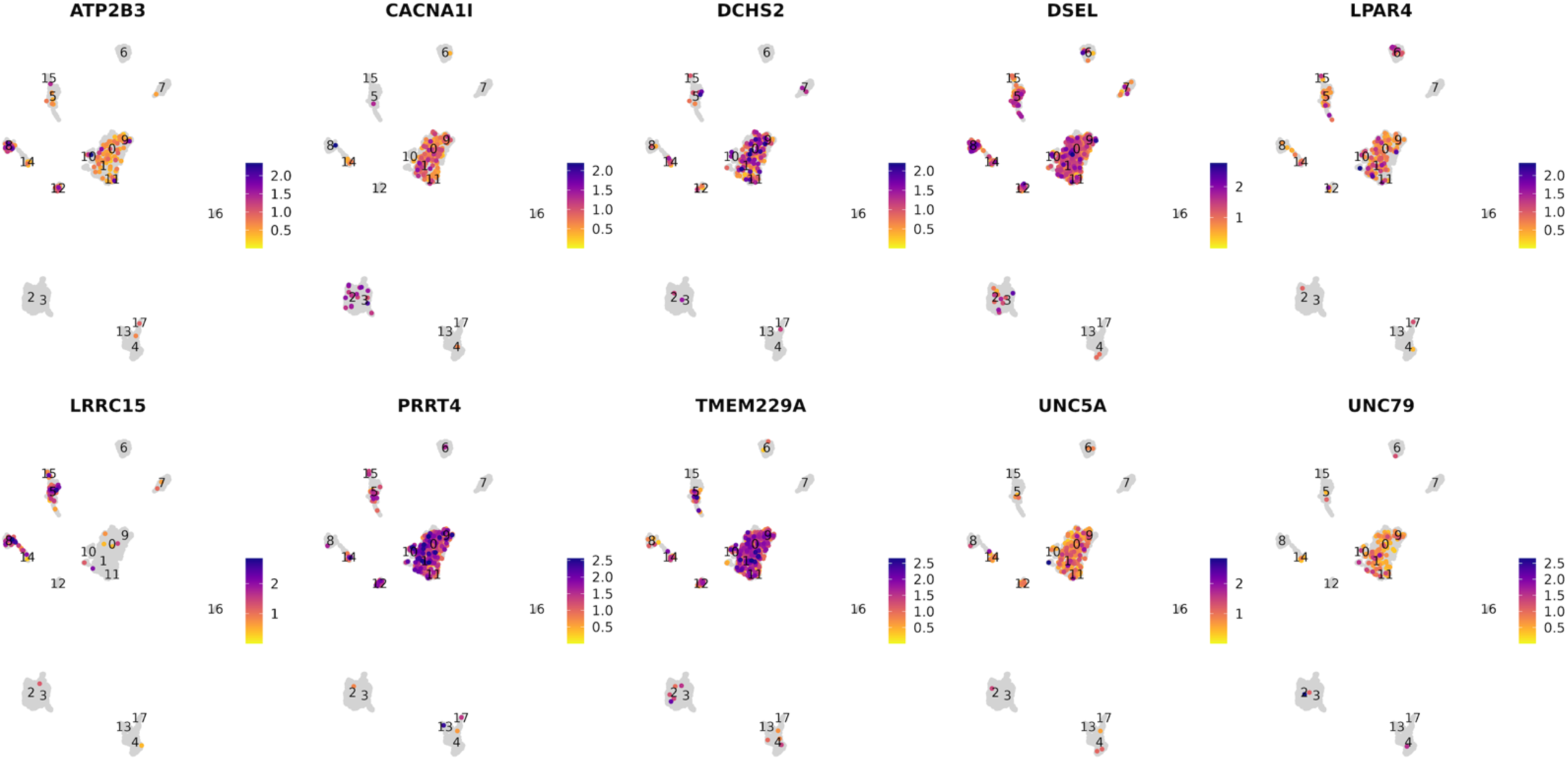
Top 10 surface putative EwS TAs expression in scRNA dataset. UMAP feature plots showing the expression patterns of the top 10 putative EwS TAs (*ATP2B3*, *CACNA1I*, *DCHS2*, *DSEL*, *LPAR4*, *LRRC15*, *PRRT4*, *TMEM229A*, *UNC5A*, and *UNC79*) across PT samples in the integrated scRNA dataset (GSE243347). Each panel shows normalized RNA expression for the selected gene across cell clusters, with purple color intensity indicating relative expression levels.

**Supplementary Figure 15.**
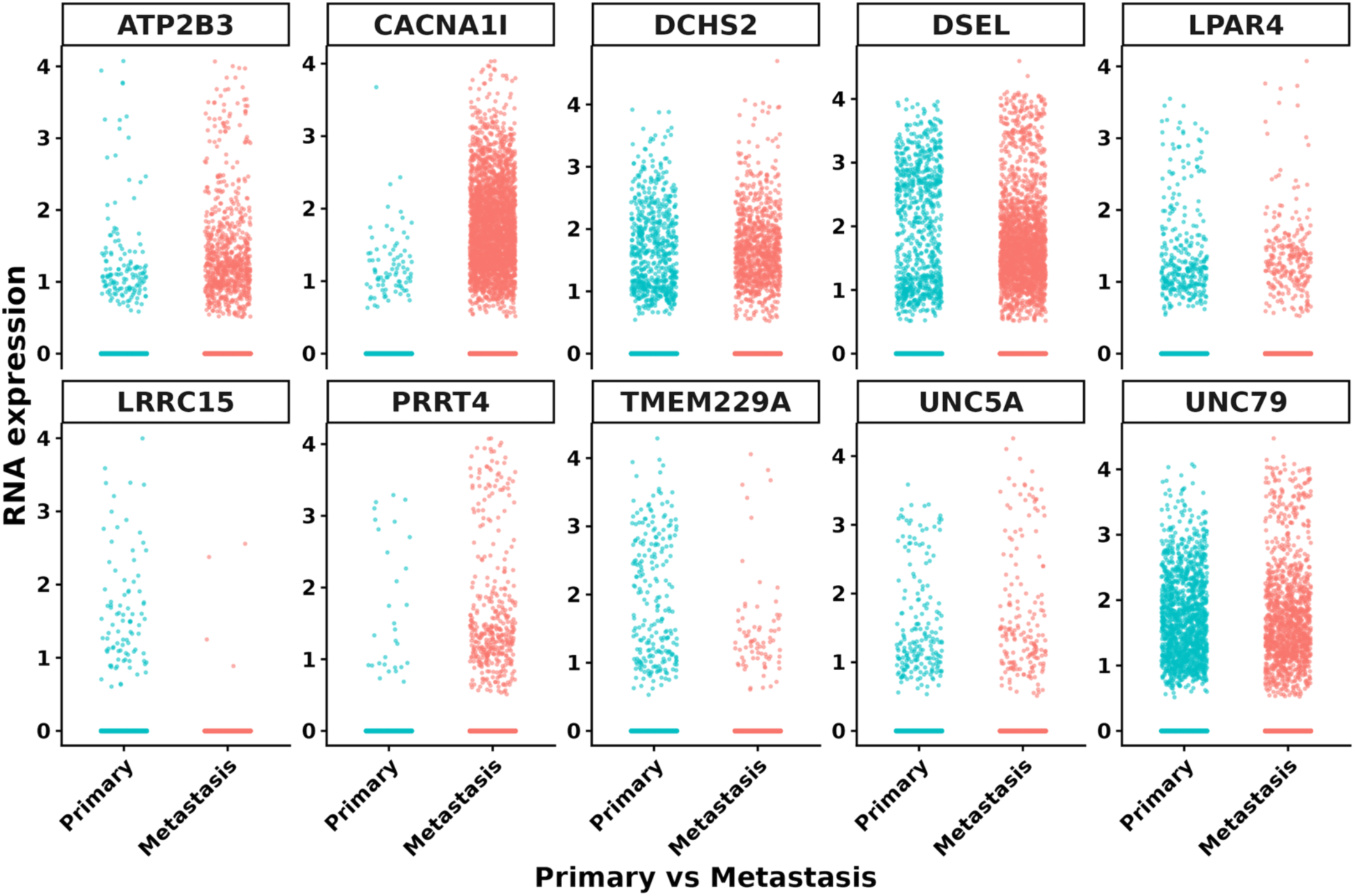
Top 10 putative EwS TAs expression across primary and metastatic EwS tumors in the snRNA dataset. Jitter plots showing RNA expression levels of putative tumor-associated antigens (*ATP2B3*, *CACNA1I*, *DCHS2*, *DSEL*, *LPAR4*, *LRRC15*, *PRRT4*, *TMEM229A*, *UNC5A*, and *UNC79*) across primary (blue) and metastatic (red) EwS tumor nuclei in the snRNA-seq dataset. Each point represents a single nucleus, colored by tumor origin. Expression values were derived from normalized RNA expression. Several candidates, including ATP2B3, CACNA1I, and PRRT4, show higher expression in metastatic tumor cells, whereas others (e.g., DCHS2, DSEL, UNC5A, and UNC79) exhibit comparable expression across primary and metastatic states. These data highlight conserved, primary-, and metastasis-associated transcriptional patterns among the putative TAs.

**Supplementary Figure 16.**
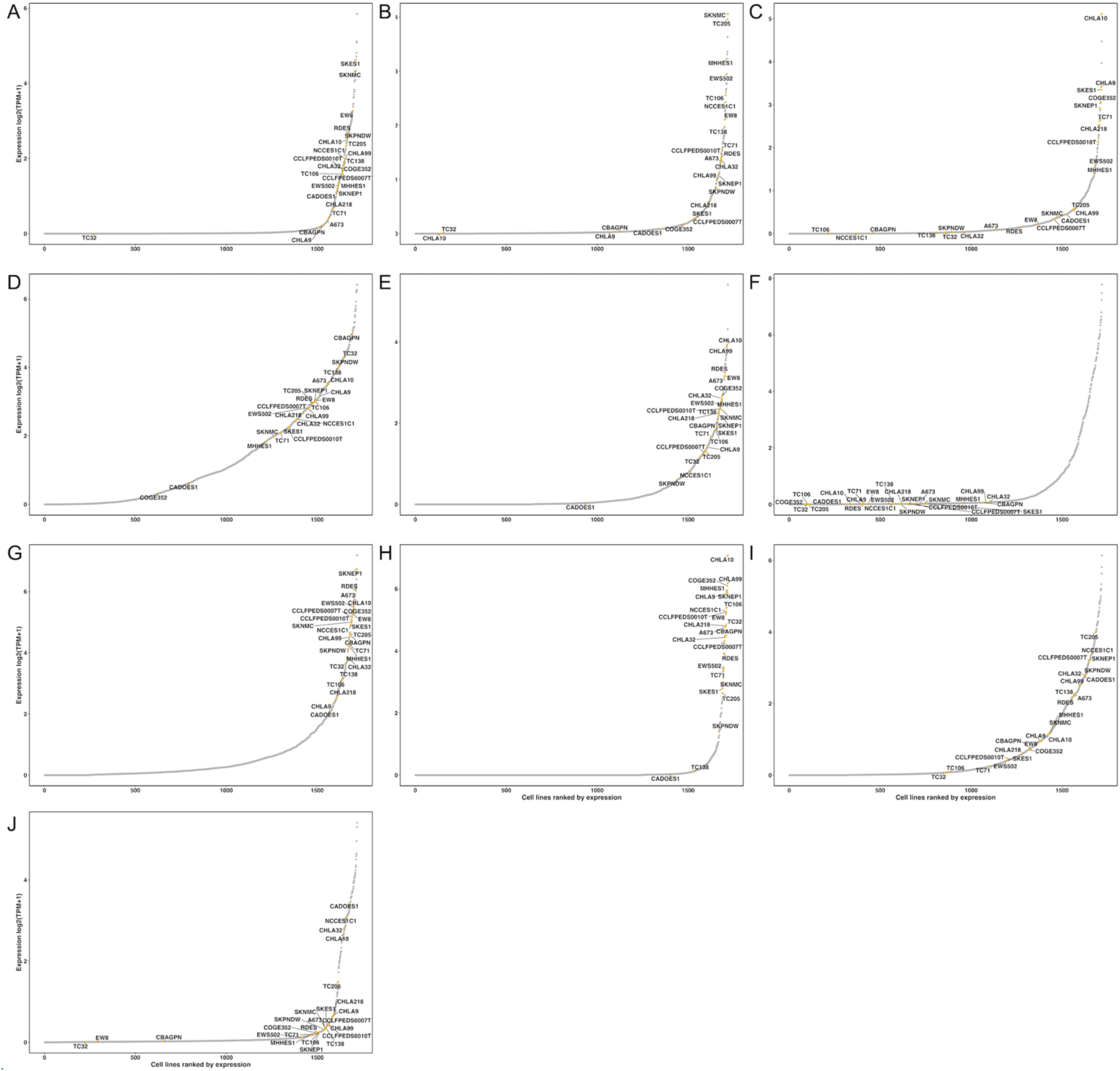
Transcript expression of the 10 putative EwS TAs across DepMap cancer cell lines. RNA expression of (A) *ATP2B3*, (B) *CACNA1I*, (C) *DCHS2*, (D) *DSEL*, (E) *LPAR4*, (F) *LRRC15*, (G) *PRRT4*, (H) *TMEM229A*, (I) *UNC5A*, and (J) *UNC79* was obtained from the DepMap Public 26Q1 short-read RNA-sequencing dataset. Cancer cell lines were ranked according to increasing expression, and transcript abundance is shown as log₂(TPM + 1). EwS cell lines are indicated in gold and labeled, whereas all other cell lines are shown in gray. This analysis places the expression of each TAs candidate in the context of a pan-cancer cell line panel.

**Supplementary Figure 17.**
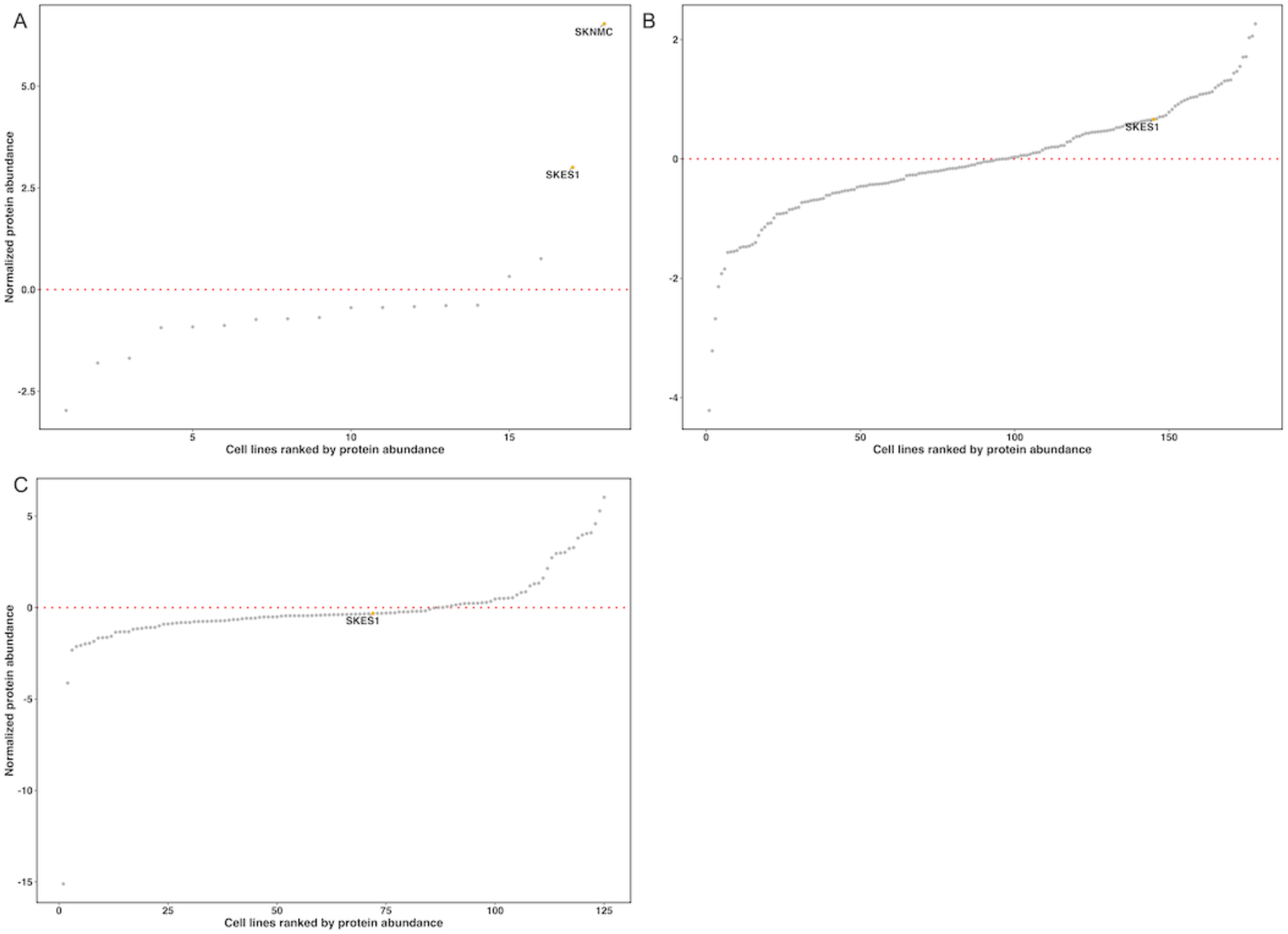
Protein abundance of three putative EwS TAs across DepMap cancer cell lines. Normalized protein abundance of (A) ATP2B3, (B) DCHS2, and (C) LRRC15 (LRRC15_2 isoform) was obtained from the DepMap harmonized CCLE mass spectrometry–based proteomics dataset. Cancer cell lines were ranked according to increasing protein abundance. Each gray dot represents a single cell line, whereas EwS cell lines are indicated in gold and labeled. The red dotted horizontal line indicates a normalized protein abundance value of 0. The harmonized proteomics dataset reports protein abundance values normalized across cell lines, with positive and negative values indicating higher and lower protein abundance, respectively.

## Supporting information

ST1_ALSF_Metadata

ST2_ GSE243347_Metadata

ST3_ Gene_Sets

ST4_Bulk_RNA_Metadata

